# A proton transfer mechanism in the malaria parasite lactate/H^+^ symporter reveals a channel-like transporter without conformational changes

**DOI:** 10.64898/2026.02.05.704099

**Authors:** Ciara Wallis, Kasimir P. Gregory, Stephen J. Fairweather, Giel G. van Dooren, Adele M. Lehane, Ben Corry

**Affiliations:** Research School of Biology, The Australian National University, Canberra, Australia; School of Science and Technology, The University of New England, Armidale, Australia

## Abstract

The malaria parasite *Plasmodium falciparum* relies on anaerobic glycolysis for energy during the intraerythrocytic stage, producing lactate and H^+^ that must be extruded via its formate-nitrite transporter (PfFNT) to prevent death by cytosolic acidification and swelling. Unlike bacterial homologues classified as channels, PfFNT mediates saturable lactate transport and was classified as a transporter, raising questions about its transport mechanism and what distinguishes channels from transporters. Here, we combine over 720 μs of molecular dynamics simulations, quantum chemical calculations, and transport assays in *Xenopus laevis* oocytes to characterize the transport cycle of PfFNT. We show a transport mechanism in which protonation of a central histidine (H230) enables lactate binding, and proton transfer from H230 to lactate is required to release the substrate as neutral lactic acid. This mechanism is supported by data indicating that neutral compounds such as lactamide do not enter the cavity, while anions like iodide, which are unlikely to protonate under physiological conditions, bind in the cavity but are not released and thus block lactate transport. As neither lactate nor lactic acid can diffuse passively through the protein, it is the requirement for proton transfer defines PfFNT as a lactate/H^+^ transporter rather than a channel. Notably, the PfFNT transport process occurs without significant protein conformational changes. Furthermore, our data indicates that under acidic conditions, small neutral molecules such as formic acid can passively diffuse through the protein. This suggests PfFNT challenges standard definitions of channels and transporters and differs from existing examples of channel-like transporters as it i) operates as a transporter without conformational changes and ii) can switch between transporter and channel modes within the same transport pathway.

## Introduction

### What is the difference between transporters and channels?

Proteins that move substrates across a biological membrane are classified into two groups: channels and transporters. While channels may open and close, at some stage they create a continuous water-filled pore that allows for the rapid, often selective, diffusion of solutes down their electrochemical gradient. In contrast, transporters never have a continuous pathway for substrate translocation; instead, they undergo conformational changes after substrate binding to shuttle substrates across the membrane - often against their concentration gradients.^1^ The distinction between channels and transporters dates back to at least the 1950s^2–5^ and is not merely a matter of semantics: correctly identifying modes of substrate transport has important implications for how protein function is interpreted. Mechanistic classification can inform how mutations are expected to alter protein activity, guide the rational design of inhibitors by revealing whether substrates move through an open pore or via conformational cycling, and additionally shape our understanding of how membrane transport proteins evolved to fulfil diverse physiological roles.

While in most cases the distinction between channels and transporters is clear, studies of some protein families have shown that the boundary is not always distinct. For example, the chloride channel (CLC) family contains highly homologous proteins, some of which are passive Cl^-^channels while others are transporters that actively exchange Cl^-^ and H^+^.^6–9^ The SLC26 family also contains both channel and transporter members that share conserved structural features.^10,11^ The cystic fibrosis transmembrane conductance regulator (CFTR) is a member of the ATP-binding cassette transporter family, but functions as a channel for Cl^-^ ions.^12^ Another member of the ATP-binding cassette family, ABCG2, has also been proposed to have a channel-like translocation pathway.^13^ A sperm-specific Na^+^/H^+^ exchanging secondary-active transporter, SLC9C1, was found to have voltage-sensing and cyclic-nucleotide binding domains – features previously only associated with voltage-gated cyclic-nucleotide-modulated ion channels.^14–17^ Glutamate transporters EAAT1 and GltPh from the SLC1A family were identified to have an additional Cl^-^ channel function uncoupled from substrate transport.^18,19^ The human inorganic phosphate exporter, XPR1, has also recently been shown to have a channel-like conduction mechanism, despite being previously thought to be a transporter.^20–22^ While these instances are infrequent, they underscore the formidable challenge of precisely distinguishing between channels and transporters - illuminating the complex nature of their classification.

### The formate-nitrite transporter family of promiscuous pentameric channels

The formate-nitrite transporter (FNT) family comprises integral membrane proteins that mediate the transport of monocarboxylates^23–30^ and certain inorganic ions (like nitrite, chloride, and hydrosulfide)^24,27,31–33^ across membranes of bacteria, protists, and some algae. FNTs are believed to share a highly conserved homopentameric structure.^34–38^ Despite containing the word “transporter” in their name, bacterial members of the family characterized thus far through experimental^26,27,31,39^ and computational^40–44^ studies have been described as channels with minimal selectivity for small, anionic substrates. However, a member of this family from the malaria-causing parasite *Plasmodium falciparum,* known as PfFNT, was proposed to be a transporter^45,46^. This initial classification of PfFNT was predominantly based on the observation that PfFNT-mediated lactate/H^+^ transport was saturable, a kinetic property commonly associated with transporters. Additionally, PfFNT was shown to transport both L-lactate and D-lactate and was reported to have a similar degree of stereoselectivity for L-lactate over D-lactate as seen in MCT4 from the human monocarboxylate transporter family – further drawing comparisons between PfFNT and known lactate transporters.^45–49^ As other FNTs had been previously classified as channels, this raises the question of what differentiates PfFNT from bacterial FNTs, and more broadly, what distinguishes channels and transporters.

### PfFNT: an essential lactate/H^+^ symporter in the malaria parasite

PfFNT has gained interest as a potential antimalarial drug target as it is essential for the parasite, unrelated to human proteins, and has not previously been a target for antimalarials.^50,51^ Clinical symptoms associated with malaria arise during the intra-erythrocytic stage of the parasite’s lifecycle, during which *P. falciparum* relies extensively on anaerobic glycolysis (**Fig. 1a**) to meet metabolic demand - producing L-lactate and protons (H^+^) at a rate up to 100 times greater than that in the cytosol of uninfected erythrocytes.^52–54^ To prevent lethal decreases in cytosolic pH and increases in cell volume due to lactic acid accumulation, the parasite depends on plasma-membrane localised PfFNT to facilitate the removal of L-lactate/H^+^ from the cytosol.^28,55–57^

**Figure 1.**
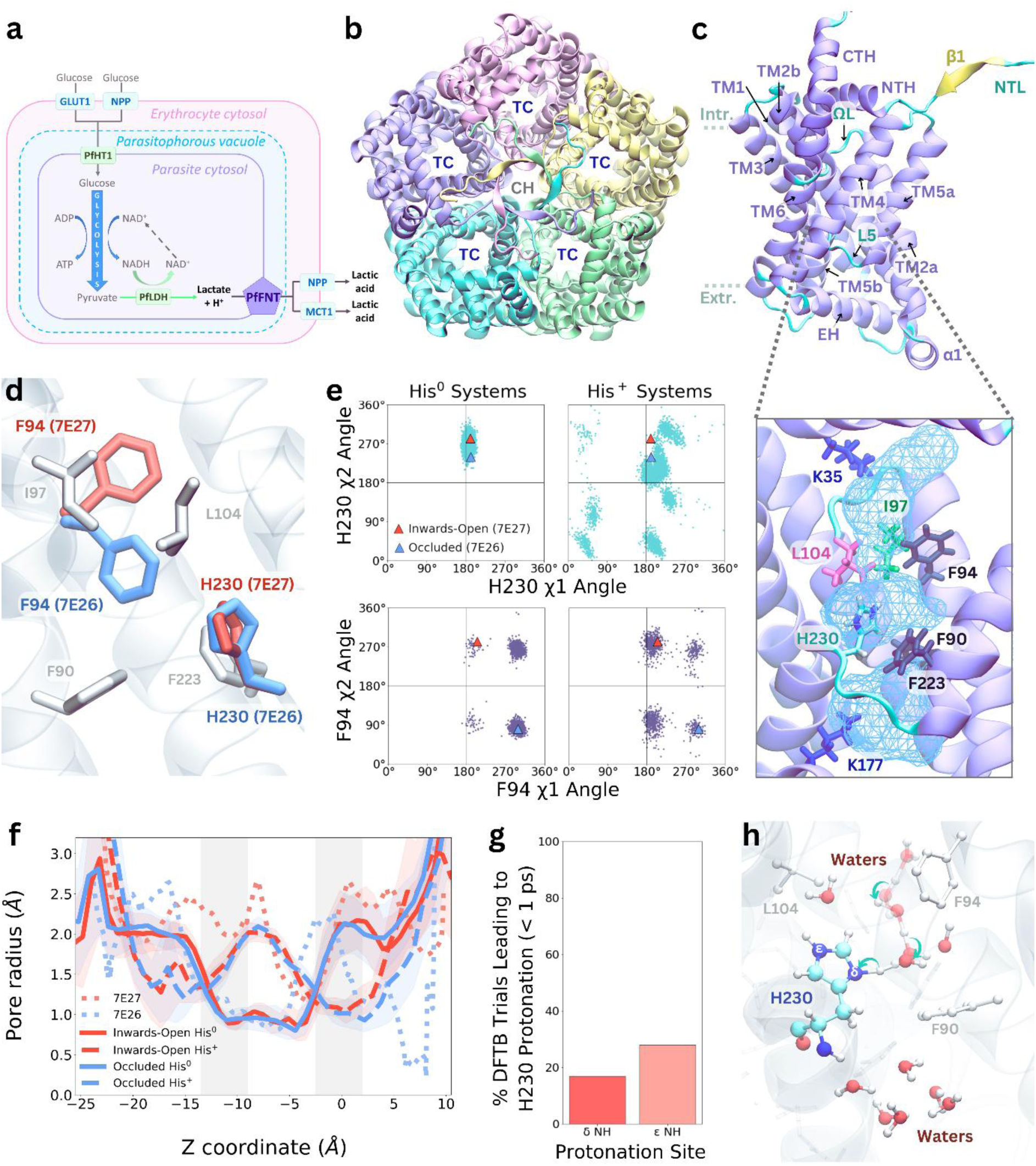
H230 protonates via a Grotthuss mechanism, increasing the radius of the PfFNT transport cavity. **a)** Schematic of PfFNT’s role in removing lactate and H^+^, major waste products from *P. falciparum* metabolism, from the parasite cytosol. Other proteins such as the human glucose transporter GLUT1 and parasite hexose transporter PfHT1 (required for glucose uptake into the host red blood cell and parasite cytosol, respectively), the parasite lactate dehydrogenase enzyme PfLDH (converts glycolytic waste products to lactate and H^+^), and the human monocarboxylate transporter MCT1 (required to remove lactate/H^+^ from the host red blood cell), and the parasite-induced new permeability pathways (NPPs) for nutrient uptake and export are also shown.^74–77^ **b)** Top-down view of the intracellular face of PfFNT (PDB ID: 7E27), with each subunit coloured differently, showing the location of the transport cavities (TC) and central hole (CH). **c)** Topology of a single PfFNT subunit. Transmembrane helices TM1, TM2a, TM2b, TM3, TM4, TM5a, TM5b, and TM6, as well as the C-terminal helix (CTH), N-terminal helix (NTH), extracellular helix (EH), and the juxtamembrane helix (α1) are shown in purple. The Ω loop (ΩL), connecting TM2a and TM2b, loop 5 (L5), connecting TM5a and TM5b, and the N-terminal loop (NTL) are shown in cyan. A beta-sheet (β1), between the NTH and NTL is shown in yellow. A close-up view of the transport cavity within a subunit is also shown. The intracellular constriction residues (F94, I97, and L104) and extracellular constrictions (F90, F223, and H230) are shown in licorice. Two key lysine residues, K35 and K177, are also shown. The shape of the cavity shown in the light blue mesh, is calculated from the area explored by water molecules in MD simulations with charged H230. **d)** Snapshot following alignment of the Inwards-Open (PDB ID: 7E27, red) and Occluded (PDB ID: 7E26, blue) PfFNT structures, showing the sidechain positions of F94 and H230. Other constriction residues are shown in grey. **e)** Janin plots showing the χ1 and χ2 dihedral angles adopted by H230 (teal) and F94 (purple) when H230 is either neutral or charged. Data from all five subunits and all three replicates for both inwards-open and occluded systems were combined on the same plot. Triangles are added to represent the starting dihedral angle from the inwards-open (red) and occluded (blue) cryo-EM structures. **f)** The average pore radius (± SEM) across each frame of the ensemble trajectory for the Inwards-Open His^0^ (red, solid line), Inwards-Open His^+^ (red, dashed line), Occluded His^0^ (blue, solid line), and Occluded His^+^ (blue, dashed line) systems. Light grey shaded regions indicate the locations of the intracellular constriction (“Intr. Constr.”) and extracellular constriction (“Extr. Constr.”). Radii of the cryo-EM structures are shown as blue (7E26) and red (7E27) dotted lines. **g)** Percentage of trials that led to protonation of either the δ or ε nitrogen of H230 within 1 ps of DFTB. **h)** Snapshot from DFTB simulations showing an example route of proton transfer.

Two antiplasmodial compounds from the Medicines for Malaria Venture (MMV) Malaria Box (MMV007839 and MMV000972) and more recently synthesized PfFNT inhibitors (e.g. BH267.meta) were found to potently inhibit PfFNT-mediated lactate/H^+^ transport, resulting in acidification of the parasite cytosol, parasite swelling, and parasite death.^56–58^ However, prolonged *in vitro* exposure of parasites to MMV007839 or BH267.meta led to the emergence of mutations in PfFNT that reduced the potency of known inhibitors.^56,57,59^ Understanding how transport occurs in PfFNT will help illuminate the distinction between channels and transporters and may support the rational design of new PfFNT inhibitors that are less susceptible to resistance mutations.

### Are FNTs channels or transporters?

The discrimination of FNTs as channels or transporters has previously been determined using at least one of two criteria. The first is whether substrate transport is saturable. Relative to channels, transporters generally exhibit a higher substrate affinity (lower K_M_ values) and saturate at much lower substrate concentrations. In contrast, channels tend to have lower affinity and are either non-saturable or saturate only at much higher concentrations.^1^ Measurements of the transport of L-lactate and formate across the plasma membrane in isolated *P. falciparum* trophozoites and in PfFNT-expressing *Xenopus laevis* oocytes yielded K_M_ values in the low mM range (3.8 mM and 7.3 mM for lactate and 1.9 mM and 5.1 mM for formate in the parasites and oocytes, respectively).^28^ However, much lower substrate affinity has been observed for L-lactate transport in PfFNT-expressing *Saccharomyces cerevisiae* (K_M_ 87 mM)^55^, possibly due to variations in the lipid environment and/or in protein folding between expression systems. While the few bacterial FNTs for which transport kinetics have been investigated generally have higher K_M_ values than PfFNT, it is not obvious based on affinity alone where to draw the boundary between transporter and channel members of this family.^27,60^

The second criterion used to distinguish transporters from channels is whether conformational changes are required during transport. Transporters, like the human lactate transporter MCT1, often undergo substantial conformational changes during the substrate transport process.^61,62^ While mobility of the N-terminal helix has been suggested to mediate gating in some bacterial FNTs, no resolved FNT structures show conformational changes associated with substrate transport.^26,39–42^ In all cases, the authors conclude that these bacterial FNTs (EcFocA, VcFocA, StFocA, StNirC, and CdHSC) function as channels.^23,26,31,32,39^ Substrate channelling events have been observed in unbiased MD simulations using structures of EcFocA and CdHSC and a homology model of another FNT, EcYfdC.^43^ Channelling events were also observed in simulations of VcFocA with an imposed concentration gradient.^42^

### Structures of PfFNT and the role of H230

In 2021, two independent groups resolved cryo-EM structures of PfFNT in an apo state (termed ‘occluded’ conformation) and a state with the inhibitor MMV007839 bound (termed ‘inwards-open’ conformation).^63,64^ These revealed a characteristic FNT homopentameric structure with five independent transport cavities, featuring hydrophobic constrictions on both the intracellular (inside the parasite cytosol) and extracellular (inside the parasitophorous vacuole) sides formed by F94, I97, and L104 and F90, F223, and H230, respectively (**Fig. 1b-c, Fig. S1a-c**). The highly conserved H230 is hypothesized to play a critical role in PfFNT activity.^55,63,64^ While some FNTs, such as EhFNT from the amoebiasis-causing parasite *Entamoeba histolytica* and BtFdhC from *Bacillus thuringiensis*, can transport substrates despite not having a cavity histidine^65^, mutation of this residue in PfFNT was shown to abolish L-lactate uptake.^64^

Histidine sidechains can be neutral or positively charged depending on their protonation state, with the likelihood of protonation depending on the surrounding environment. Computational studies on bacterial FNTs implicate the cavity histidine in substrate transport^40–44^, though Schmidt & Beitz (2021) suggested the cavity histidine would be unlikely to be protonated in such a hydrophobic environment^66^. In transport assays in PfFNT-expressing *S. cerevisiae*, lactate uptake increased with decreasing external pH, with optimal transport at pH 3.9.^46^ As this is around the pK_a_ of lactic acid (pK_a_ 3.86), this heightened transport was attributed to the higher likelihood of either H230 or lactate becoming protonated. Notably, transport decreased dramatically at pH < 3.9, where both species are likely to become protonated.^46^

### Proposed mechanisms for transport in PfFNT

In the absence of structures of PfFNT with substrates bound, two main transport mechanisms have been proposed. The dielectric shift model suggests lactate is drawn into the cavity by lysine residues and protonated by water as the environment becomes more hydrophobic, implying a channel-like process requiring neutral lactic acid.^60,67^ The proton transfer model proposes that negatively charged lactate binds to H230, which then protonates it to lactic acid for release.^55,64^ However, the role of H230 and the validity of these mechanisms remain uncertain.

### This study

In this study, we used molecular dynamics (MD) simulations, quantum chemical calculations, and transport assays in *Xenopus laevis* oocytes to investigate how PfFNT transports substrates and whether it functions as a transporter or a channel. Our simulations indicate that anionic lactate, not neutral lactic acid, is the substrate that binds within the transport cavity before becoming protonated by a charged H230 residue to get released - highlighting the importance of H230 for substrate binding and transport. Other known substrates, such as formate, also appear to be transported via a proton transfer mechanism at neutral pH. Iodide, an anion with an extremely low proton affinity that is not expected to protonate under biologically relevant pH conditions, binds within the cavity in our simulations but appears unlikely to get protonated to be released. In transport assays, we show that iodide strongly inhibits PfFNT-mediated lactate transport, likely reflecting iodide blocking the cavity once bound and interfering with lactate entry. While the entire transport mechanism occurs with almost no conformational changes, reminiscent of channels, neither lactate nor lactic acid have a continuous pathway through the transport cavity. We conclude that the requirement for proton transfer for anionic substrates at cytosolic pH allows PfFNT to be defined as a transporter rather than a channel. Interestingly, we observe channel-like permeation of formic acid in our simulations under acidic conditions where both formic acid and H230 are protonated, suggesting that PfFNT may alternate between transporter and channel function depending on the condition and substrate size.

## Results

### Protonation of H230 influences the structural dynamics of the transport cavity

#### Inwards-open and occluded states are not distinct

Given the suggested importance of the central histidine in FNT, we first investigated what protonation state H230 is likely to be in during PfFNT transport by examining how the dynamics of the transport cavity differ between the ‘inwards-open’ and ‘occluded’ states when H230 is either positively charged (His^+^) or neutral (His^0^) (**Fig. S2**). The cryo-EM structures of PfFNT in the ‘inwards-open’ inhibitor-bound and ‘occluded’ apo states are highly similar (RMSD 0.19 Å) and differ subtly by the positioning of the F94 and I98 sidechains (**Fig 1D-E; Fig. S3**). Our simulations show that in the absence of the inhibitor, regardless of H230 protonation state, the transport cavity does not remain in the ‘inwards-open’ conformation. Further, we show that there are no obvious differences between ‘inwards-open’ and ‘occluded’ simulations (**Fig. S2; Fig. S3**). However, the pore radius and orientation of the H230 and F94 sidechains differ appreciably between His^0^ and His^+^ systems (**Fig. 1f**). While the cryo-EM structures appear to represent neutral H230 (as suggested from the sidechain orientation) (**Fig. 1d-e**), the narrow pathway observed in His^0^ systems (< 1 Å radius) would likely hinder substrate entry (**Fig. S2p**). The N-terminal helix, a structure previously suggested to mediate gating in other FNTs^35^, did not move significantly between systems. However, slight differences in the Ω-loop, another structure indicated in FNT gating^34^, were observed between His^0^ and His^+^ systems (**Fig. S2**).

#### H230 can protonate via a Grotthuss mechanism

The widening of the cavity observed in His^+^ systems suggests that protonation of H230 may be required to enable substrates, such as lactate, to enter. We used various methods to predict the likelihood of H230 becoming protonated at cytosolic pH, including PROPKA3.0^68^ and different implementations of continuous constant pH MD simulations^69–73^. These methods yielded pK_a_ estimates ranging between 2.51 and 7.13 (**Fig. S4a-b**), making it difficult to confirm the most likely protonation state. Notably, CpHMD simulations performed with AMBER24 showed that the orientation of the F94 sidechain influenced the pK_a_ of H230 (**Fig. S4b**), indicating the likelihood of H230 protonation is highly sensitive to subtle changes in protein conformation. As no nearby protein residues appear likely to donate protons to H230, we next investigated the possibility of H230 protonating via a Grotthuss mechanism, in which proton transport occurs via protons ‘hopping’ between adjacent, hydrogen-bonded water molecules rather than diffusing as individual ions. To do this, we used density functional tight binding (DFTB) simulations starting from over 200 different MD snapshots of a 5 Å region surrounding H230, where H230 was neutral (protonated at either the δ or ε nitrogen). In ∼20% of cases, within 1 ps of simulation, we observe the proton ‘hopping’ between water molecules into the cavity from either the intracellular or extracellular side, leading to protonation of the δ or ε nitrogen of H230 (**Fig. 1g-h; Fig. S4e; Movie S1**).

### Molecular simulations suggest a proton transfer mechanism for lactate/H^+^ transport

#### Negatively charged Lac^-^ binds to positively charged His^+^

To determine if protonation of H230 and/or the substrate influences binding, we performed MD ‘flooding’ simulations in which a high concentration of either lactate (Lac^-^) or lactic acid (Lac^0^) was distributed across both sides of the membrane in His^0^ and His^+^ systems. In this way, we tested all possible combinations of substrate and H230 protonation state to see whether binding could occur within the timeframes of our simulations. While Lac^0^ was previously suggested to be the species that would more favourably pass the intracellular and extracellular constrictions^78^, we did not observe it entering the cavity regardless of H230 protonation state **(Fig. 2a,e; Fig. S5a**).

**Figure 2.**
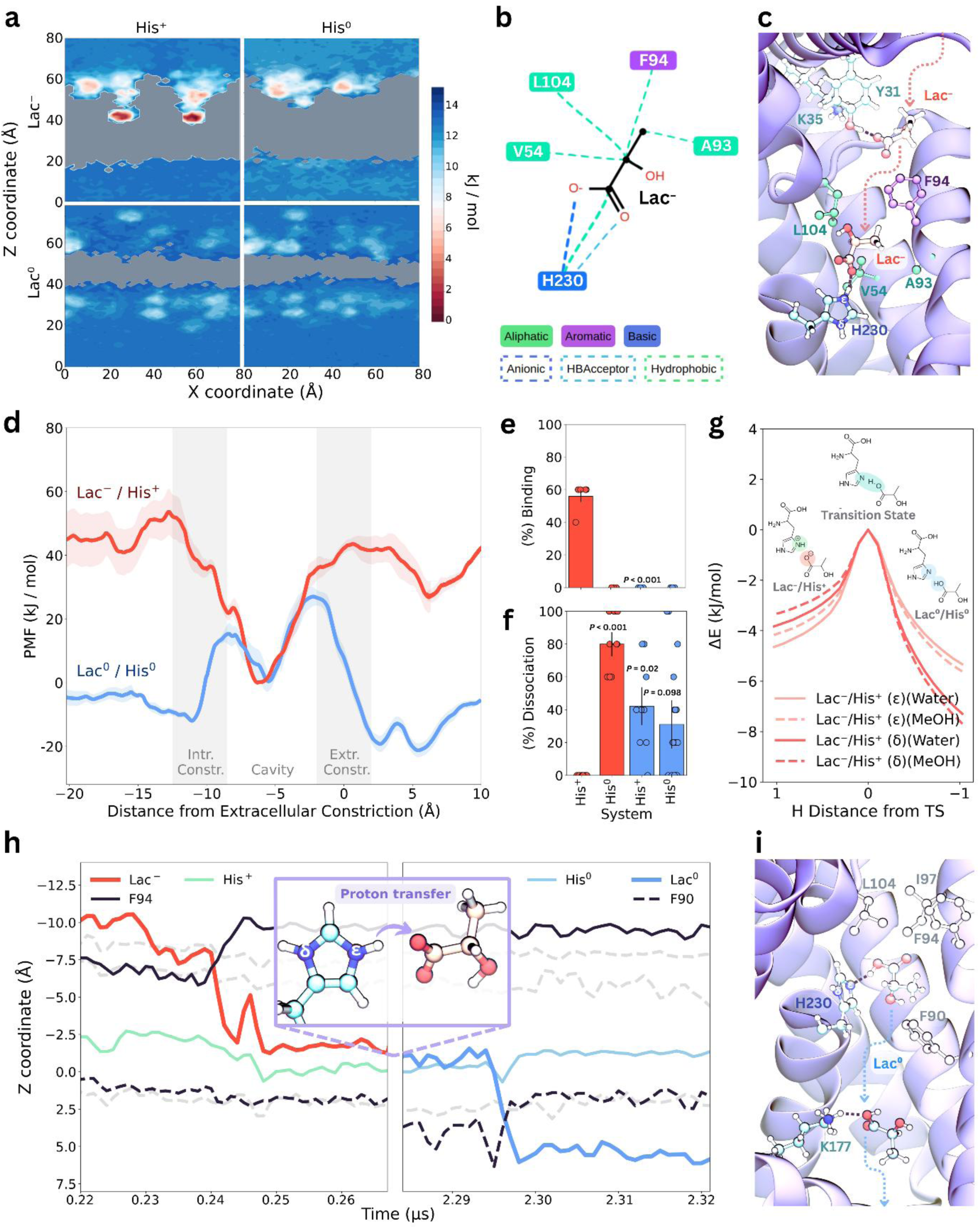
Lac^-^/H^+^ transport in PfFNT occurs via a proton transfer mechanism. **a)** Free energy surfaces (FES) of regions explored by substrates in the X-Z projections of Lac^-^/His^+^, Lac^-^/His^0^, Lac^0^/His^+^, and Lac^0^/His^0^ in MD flooding simulations. For Lac^-^/His^+^, two prominent minima are observed, corresponding to Lac^-^ binding events within the transport cavities of different subunits. Additional minima can be seen near the intracellular vestibule and NTH. Area not occupied by substrates is shown in grey. **b)** Interaction network of residues that interact with Lac^-^ in over 50% of frames from our replicate MD simulations. **c)** Representative snapshot from the top-ranked cluster of Lac^-^ bound to charged H230. V54, F94, A93, and H230, residues that form substantial contacts with Lac^-^, are shown in licorice. A semitranslucent representation shows Lac^-^ transiently interacting with K35, disrupting the salt bridge between K35 and D103, before entering the cavity (indicated by the dotted arrow). Hydrogen bonds between Lac^-^ and H230, Lac^-^ and K35, and K35 and D103 are also shown. In this snapshot, Lac- is bound to the ε NH group of H230, however, Lac- was also observed to interact with the δ NH group. **d)** Average potential of mean force (PMF) (kJ/mol), ± SEM, *n* = 5, determined from 1D and 2D US simulations for the entire transport pathway for the Lac^-^/His^+^ (red) and Lac^0^/His^0^ systems (blue). The distance of the substrate from the extracellular constriction is represented along the X-axis. **e)** Average percentage of subunits bound by substrate out of the total number of binding sites available for Lac^-^ and Lac^0^ flooding simulation systems (± SEM, *n* = 8). **f)** Dissociation events observed from simulations starting with substrates in the cavity, expressed as a percentage of the average total number of subunits without substrates at the end of the simulation over the total number of subunits with a substrate bound at the start of the simulation (± SEM, *n* = 18 for Lac^0^/His^0^, *n* = 6 for all other systems). Statistical significance for **e** and **f** were determined using a two-way ANOVA with a *post-hoc* Tukey test. The number of binding and dissociation events for each system are given in **Tables S2** and **S4**. Binding events for other test compounds are shown in **Fig. S14a**. **g)** Changes in potential energy (ΔE) from quantum chemical calculations of the intrinsic reaction coordinate of His^+^ donating a proton to Lac^-^ from either the δ NH or ε NH group of the H230 sidechain using an implicit water (solid lines) or implicit methanol (MeOH; dashed lines) environment. **h)** Representative traces of Lac^-^ binding and Lac^0^ dissociation in MD simulations, capturing a complete transport process. The Z coordinates for the centre of mass of Lac^-^ (red) and the charged H230 (green), F94 (black, solid), and F90 (black, dashed) sidechains are plotted over time. Other constriction residues are shown in grey dashed lines. After simulating for 2 μs, the proton was manually swapped from His^+^ to Lac^-^ to form Lac^0^ within the cavity, and new simulations were run for 1.5 μs to observe whether Lac^0^ dissociated from the cavity following proton transfer (blue trace). **i)** Snapshot of Lac^0^ transiently interacting with K177 before dissociating from the cavity.

In contrast, multiple binding events were observed for Lac^-^, but only when H230 was charged (**Fig. 2a-c,e; Fig. S5b; Movie S2a-b**). The free energy surfaces (FES) for substrates around the protein obtained from these simulations show a minimal barrier and a deep energy minimum within the cavity in the Lac^-^/His^+^ system, while Lac^0^ seemingly has a larger barrier to enter the cavity and is more stable in the bulk solution (**Fig. 2a**). In our simulations, Lac^-^was only seen to bind from the intracellular side, transiently interacting with K35 and Y31 before entering the cavity and binding tightly to charged H230. Additional hydrophobic interactions with surrounding residues (V54, A93, F94, and L104) help stabilize Lac^-^ in the binding site (**Fig. 2b; Fig. S5b-c**). In simulations with either K35C/K177C or H230N mutations, Lac^-^ binding events were not observed – supporting the importance of these residues for attracting substrates into the cavity (**Fig. S6a**).

We next rigorously quantified the ability of Lac^-^ and Lac^0^ to bind and dissociate from the transport cavity using umbrella sampling simulations. These simulations were performed for the Lac^-^/His^+^ and Lac^0^/His^0^ systems, as these systems represent lactate and proton transport in a physiologically relevant 1:1 stoichiometry. The data show Lac^-^ favourably entering the cavity from both the intracellular and extracellular directions when H230 is charged, revealing relatively low average energy barriers (∼17-20 kJ/mol) to pass the constrictions, and a deep energy well of -43 ± 6 kJ/mol inside the cavity (**Fig. 2d**). Notably, entry of Lac^-^ from the extracellular side appears dependent on the orientation of the H230 sidechain with entry only being favourable if the H230 sidechain is oriented towards the extracellular side (**Fig. S7**). In contrast, Lac^0^ has a significantly larger barrier (43 ± 3 kJ/mol) to enter the transport cavity when H230 is neutral and is more stable in the bulk solution than in the cavity. Lac⁰ permeation was observed only in our simulations after mutating all constriction residues to residues with smaller sidechains, widening the constrictions enough to allow passive diffusion (**Fig. S6b-c; Table S3**).

#### Lac^0^ dissociates if inserted into the cavity

Having shown that lactate will only bind in the cavity when it is negatively charged and His230 is positively charged, we next asked under what conditions the substrate can dissociate from the cavity to complete the transport cycle. In our flooding simulations, we did not observe Lac^-^ dissociating once bound to charged H230 and our umbrella sampling suggests Lac^-^ faces a large barrier to dissociate, while Lac^0^ is more stable in bulk solution and should readily dissociate from the cavity. We thus inserted and equilibrated substrates into the cavity in the Lac^-^/His^+^, Lac^-^/His^0^, Lac^0^/His^+^, and Lac^0^/His^0^ systems, and measured dissociation events over 1.5 µs of simulation per replicate. In all systems except the Lac^-^/His^+^ system, substrates dissociated from the cavity **(Fig. 2f, Table S4**).

Together, our simulations indicate a preference for Lac^-^ to bind within the cavity and a preference for Lac^0^ to exit the protein to enter the bulk solution, leading us to the hypothesis that the transport process requires a change in protonation state of the substrate. Given PfFNT has been suggested to transport lactate and protons in a 1:1 lactate/H⁺ stoichiometry^79^, the most likely mechanism is for charged H230 to donate a proton to Lac⁻. To test whether proton transfer from His^+^ to Lac^-^ is likely to complete transport, we used structures of Lac^-^ bound to His^+^ and performed a manual proton transfer to convert Lac^-^/His^+^ systems to Lac^0^/His^0^ systems. In multiple simulations, the newly formed Lac^0^ dissociated from the cavity, suggesting we can produce an entire transport process for Lac^-^ and H^+^ through PfFNT (**Fig. 2h-i; Movie S3b**). Lac^0^ dissociation events occurred by Lac^0^ passing either the intracellular or extracellular constriction, consistent with PfFNT transporting substrates bidirectionally^79^ (**Table S3**).

#### Proton transfer from H230 to Lac^-^ is favourable

To assess the favourability of proton transfer occurring from the H230 sidechain to Lac^-^, we performed quantum chemical calculations under different solvation conditions within the transport cavity to mimic different dielectric constants of the cavity (**Fig. 2g, Fig. S8**). Calculations of the transition pathway starting from 10 MD snapshots of Lac^-^ bound to charged H230 show that proton transfer occurs favourably in all cases, with small or negligible energy barriers, the size of which are related to the overall dielectric constant of the cavity (**Table S5**). Proton transfer also occurred spontaneously in DFTB MD simulations of a 5 Å region surrounding Lac^-^ bound to H230 within the transport cavity for all starting snapshots tested, further supporting the likelihood of this process (**Movie S3a**). Together, our data indicate that the transport cycle involves (i) protonation of H230 via a Grotthuss mechanism, (ii) binding of Lac^-^ to H230, (iii) transfer of a proton from H230 to form Lac^0^, and (iv) dissociation of Lac^0^; all of which happen spontaneously in our simulations. After proton transfer has occurred, the cavity remains widened in simulations for at least another several hundred ns, rather than immediately occluding to resemble the His^0^ simulations (**Fig. S9**).

### Non-protonatable anionic ions inhibit PfFNT transport

#### PfFNT transports formate via a proton transfer mechanism, but may channel formic acid

To test our proposed Lac⁻/H⁺ transport mechanism, we examined the impact of test compounds differing in size and propensities for protonation. These include formate (For⁻/For⁰), a small monocarboxylate transported by PfFNT and other FNTs^34,35,38,45,80–84^; iodide (I⁻), an inorganic anion; and lactamide, a structural analogue of Lac^0^. As the conjugate acid HI has a pK_a_ of approximately –10, iodide protonation is only likely in superacidic conditions, well beyond those found in biological systems. Lactamide is a neutral compound with negligible propensity for protonation or deprotonation across a broad pH range (pKₐ 13.34). The pK_a_s of all test compounds are described further in **Table 3**.

In flooding simulations, For⁻ bound within the cavity only when H230 was charged (**Fig. 3a**), displaying a deep energy minimum in umbrella sampling as observed for Lac⁻ (**Fig. 3c**). Although For⁻ did not dissociate from His⁺, DFTB calculations indicated favourable proton transfer (**Fig. 3d**). Further, when placed in the cavity, For⁰ molecules dissociated regardless of the protonation state of H230 (**Fig. 3b**). These results suggest that For⁻ is transported via the same proton transfer mechanism as Lac⁻. Notably, For⁻ entry from the extracellular side was more favourable when the H230 sidechain was oriented outward (**Fig. S10**), consistent with our earlier findings for Lac⁻.

**Figure 3.**
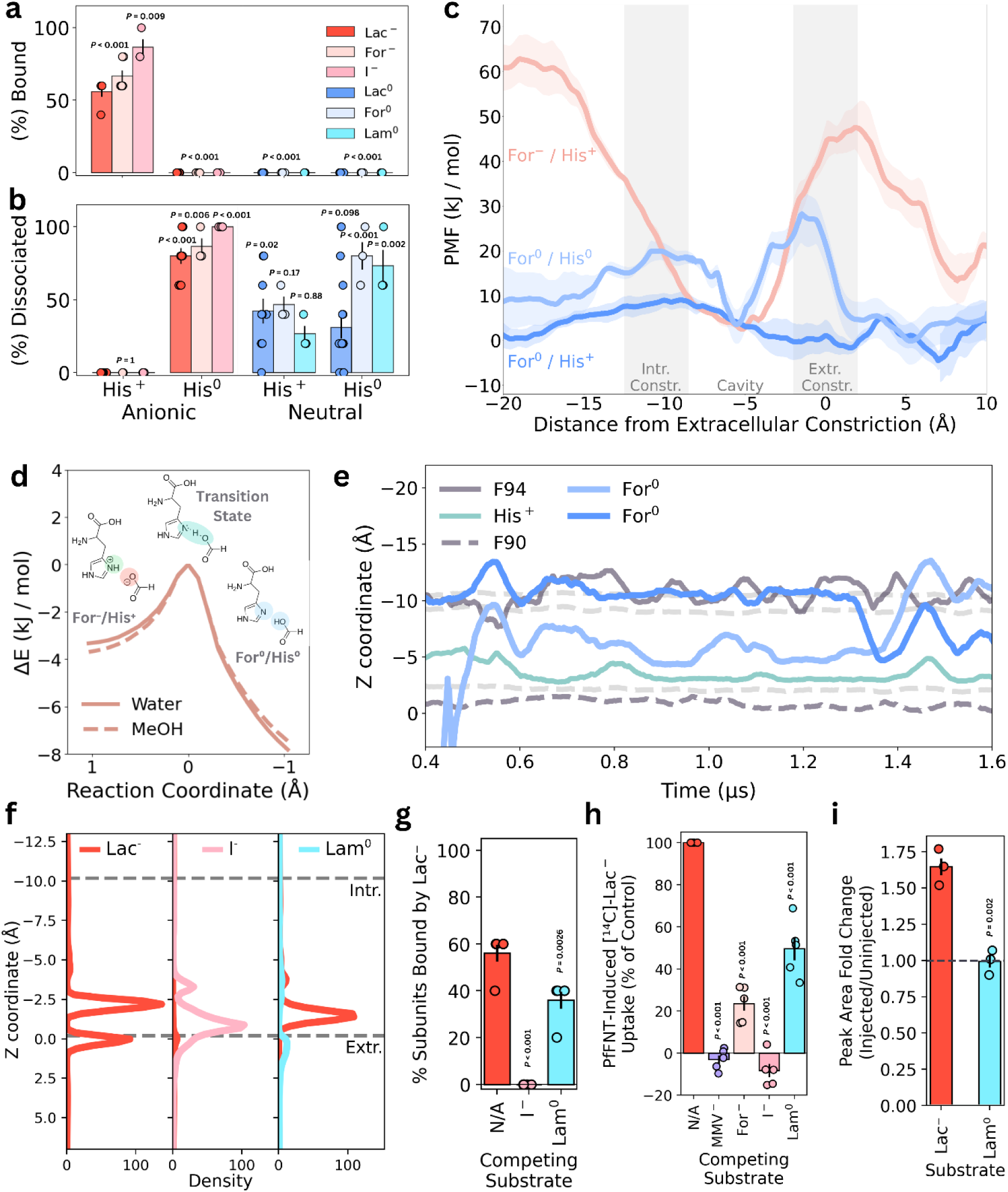
Test compounds can either be transported, channelled, or block lactate transport. Average percentages of **a)** subunits bound by substrate out of the total number of binding sites available and **b)** dissociation events observed from simulations starting with test compounds in the cavity for For^-^, I^-^, For^0^, and Lam^0^ simulation systems (± SEM, *n* = 8 for Lac^-^ and Lac^0^ systems, *n* = 6 for For^-^ and For^0^ systems, *n* = 3 for all other systems). Data for Lac^-^ and Lac^0^ are shown in both panels for reference. Statistical significance for **a** and **b** were determined using two-way ANOVAs with *post-hoc* Tukey tests. **c)** Average potential of mean force (PMF) (kJ/mol) (± SEM, *n* = 5), determined from 1D and 2D US simulations for the entire transport pathway for the For^-^/His^+^ (red), For^0^/His^+^ (grey), and For^0^/His^0^ systems (blue). The distance of the substrate from the extracellular constriction is represented along the X-axis. **d)** Changes in potential energy (ΔE) from quantum chemical calculations of the intrinsic reaction coordinate of His^+^ donating a proton to For^-^ from either the δ NH or ε NH group of the H230 sidechain in an implicit water (solid lines) or implicit methanol (MeOH; dashed lines) environment. **e)** Representative traces of For^0^ permeation events from His^+^ systems. The Z coordinates for the centre of mass of For^0^ (blue) and the charged H230 (green), F94 (black, solid), and F90 (black, dashed) sidechains are plotted over time. Other constriction residues are shown in grey dashed lines. **f)** Average substrate density across z for systems with Lac^-^ only, Lac^-^ and I^-^, and Lac^-^ and Lam^0^. **g)** Average % of subunits bound by Lac^-^ in flooding simulations with Lac^-^ only, Lac^-^ and I^-^, and Lac^-^ and Lam^0^ (± SEM, *n* = 8 for Lac^-^ only, *n* = 5 for all other systems). Statistical significance was determined using one-way ANOVA with a *post-hoc* Tukey test. **h)** The effect of different test compounds on PfFNT-mediated lactate uptake (1 mM; of which a small portion was [¹⁴C]-labelled) in *X. laevis* oocytes from five independent experiments performed on different days using oocytes from different frogs. Uptake assays were performed at pH 6.4 where both the 1 mM lactate and the test compound (10 mM in the case of formate, iodide, and lactamide or 2 μM in the case of MMV007839) were added simultaneously. For each condition, uptake was measured after a 15-minute incubation in oocytes injected with cRNA encoding PfFNT or in non-injected oocytes, and the final reported value is the level of uptake in injected oocytes minus the level of uptake in non-injected oocytes – expressed as a percentage of the total amount of lactate uptake in the absence of test compounds/MMV007839. For all conditions, the test compound was added to the oocytes at the same time as the radiolabelled substrate. Each independent experiment was performed using 8-10 oocytes per condition. Statistical significance was determined using a one-way ANOVA, blocking by experimental day, followed by a *post hoc* Tukey test. Comparisons with other substrates are shown in **Fig. S14b**. Raw data is given in **Table S6**. **i)** Average fold-change in lactate and lactamide uptake levels in oocytes measured from LC-MS experiments. PfFNT-expressing and non-injected oocytes were incubated with either lactate or lactamide for 4 hours prior to sample preparation. Fold-change was calculated as the peak area of the substrate in injected oocytes divided by the peak area in non-injected oocytes. Experiments were performed on three separate days using oocytes from three different frogs, with 40 oocytes per condition. Statistical significance was assessed using an independent two-tailed t-test. Extracted ion chromatograms for lactate and lactamide are shown in **Fig. S15**. Raw peak area data is given in **Table S7**.

In contrast to Lac^0^, formic acid (For⁰) – which is smaller in size than Lac^0^ – was shown to enter the cavity and transiently diffuse along the entire transport pathway when H230 was charged (**Fig. 3e; Table S3**). Such permeation events did not occur when H230 was neutral. This is supported by the umbrella sampling data that shows that For^0^ has negligible barriers for permeation when H230 is charged but has larger barriers to enter the cavity when H230 is neutral (**Fig. 3c**).

#### Iodide inhibits PfFNT-mediated lactate/H^+^ transport

Neutral lactamide (Lam⁰) did not bind in the cavity during flooding simulations (**Fig. 3a**) but rather was observed to transiently interact with the vestibule regions above and below the cavity (**Fig. S14c,e**). In contrast, iodide (I⁻) bound tightly within the cavity when H230 was charged (**Fig. 3a**) and did not dissociate within the timescales of our simulations (**Fig. 3b**). As I⁻ is unlikely to get protonated by H230, we expect that it is unable to be transported and would most likely inhibit PfFNT-mediated Lac⁻ transport. Co-flooding simulations showed that I⁻ blocked Lac⁻ binding in all five subunits across all replicates (**Fig. 3g; Fig. S14b; Table S2**). Notably, a significant reduction in Lac^-^ binding events was also observed from co-flooding simulations with Lam^0^.

We further explored whether the presence of known or potential test compounds inhibit PfFNT-mediated lactate/H^+^ transport by expressing PfFNT in *Xenopus laevis* oocytes – a system previously used for studying PfFNT transport activity ^28,30,57^. We tested PfFNT-mediated lactate uptake (1 mM; of which a small portion was [¹⁴C]-labelled) oocytes in the presence of 10 mM of a single test compound. The test compounds included the monocarboxylates formate, acetate, propionate, and pyruvate; the tricarboxylate citrate; the inorganic anions nitrate and iodide; and the monocarboxylic acid amides formamide, acetamide, propionamide, and lactamide. Lactate uptake in the presence of 2 μM of the known PfFNT inhibitor MMV007839 was tested as an additional control. All the test compounds caused a decrease in [^14^C]-lactate uptake (**Fig. 3h; Fig. S14b; Table S6**).

In simulations of the test compounds, lactamide, propionamide, acetamide, formamide, and citrate did not enter the cavity despite significantly decreasing PfFNT-mediated [^14^C]-lactate uptake (**Fig. S14a-e**). However, formamide was capable of entering the transport cavities in simulations. Lactamide, propionamide, acetamide, and citrate were all observed to interact with the vestibule regions above and below the cavity, suggesting that they hinder lactate permeation by transiently obscuring the transport pathway rather than competing directly for binding to H230 (**Fig. 3h; Fig. S14**). We further investigated the likelihood of PfFNT transporting lactamide by incubating PfFNT-expressing and non-injected oocytes in either lactate or lactamide for 4 hours before extracting the intracellular solution for LC-MS analysis. While lactate and lactamide could both successfully be detected in oocyte samples, the level of lactamide uptake was the same in PfFNT-expressing and non-expressing oocytes, while lactate uptake was approximately 1.5-fold higher in PfFNT-expressing oocytes than in non-expressing oocytes (*p =* 0.002, independent t-test) (**Fig. 3i; Fig. S14e**). These results indicate that PfFNT is transporting lactate but not lactamide.

Of all the tested compounds, only iodide showed an almost-complete impairment of [¹⁴C]-lactate uptake (**Fig. 3h**). In contrast, nitrate, another small inorganic anion with a low pKₐ, reduced [¹⁴C]-lactate uptake by a similar degree as formate but not as severely as iodide. In our simulations, nitrate binds to charged H230 within the cavity (**Fig. S14a, c**), but unlike iodide, it can be protonated by H230 in quantum chemical calculations (**Table S5**). As iodide binds only to charged H230 and cannot be protonated for release, we propose that it persistently remains bound within the cavity, blocking lactate/H^+^ transport.

## Discussion

### PfFNT is the first example of a transporter without conformational changes

While the existence of protein families with both channel and transporter members is rare, there has been an increasing amount of evidence over the last few decades of proteins/protein families that blur the boundaries between these once distinct classes of membrane transport proteins.^1,7^ Among these are the FNT family^85^, with our work showing PfFNT to have characteristics that challenge traditional definitions of what constitutes channels and transporters. Unlike other proteins suggested to be borderline between transporters and channels, we suggest a unique example of a protein that does not require major conformational changes for transport and is likely able to function as both a transporter and a channel without requiring separate pathways for the different modes of substrate transport. **Our data indicate firstly that PfFNT is a transporter for lactate/H^+^ under physiological conditions, as neither lactate nor H^+^ has a continuous pathway through the entire transport pathway as would be expected for a channel. However, PfFNT does not need to undergo the large-scale conformational changes usually associated with transporters during the transport cycle.** We have shown that Lac^-^/H^+^ symport occurs through a process where: **1)** H230 becomes protonated via a Grotthuss mechanism, **2)** Lac^-^ enters the cavity and binds tightly to charged H230, **3)** H230 transfers a proton to Lac^-^, and **4)** the newly formed neutral Lac^0^ dissociates (**Fig. 4**). These data support a previously proposed proton transfer mechanism.^55,64^ Throughout the transport process, the protein does not undergo substantial conformational changes, with only the sidechains of H230 and F94 having noticeable movement during transport. **Secondly, our data suggest PfFNT may shift between transporter- and channel-like transport depending on pH and substrate size**. Our simulations indicate For^-^ is also transported by a proton transfer mechanism, while For^0^, which is a smaller molecule than Lac^0^, can passively channel through PfFNT when H230 is charged (**Fig. 3a-e; Movie S4**). While the saturable kinetics observed for PfFNT-mediated For^-^ transport at neutral pH remain consistent with PfFNT functioning as a For^-^/H^+^ transporter, we propose that PfFNT can function as a channel for formate at low pH, when both H230 and the substrate are likely to be protonated.

**Figure 4.**
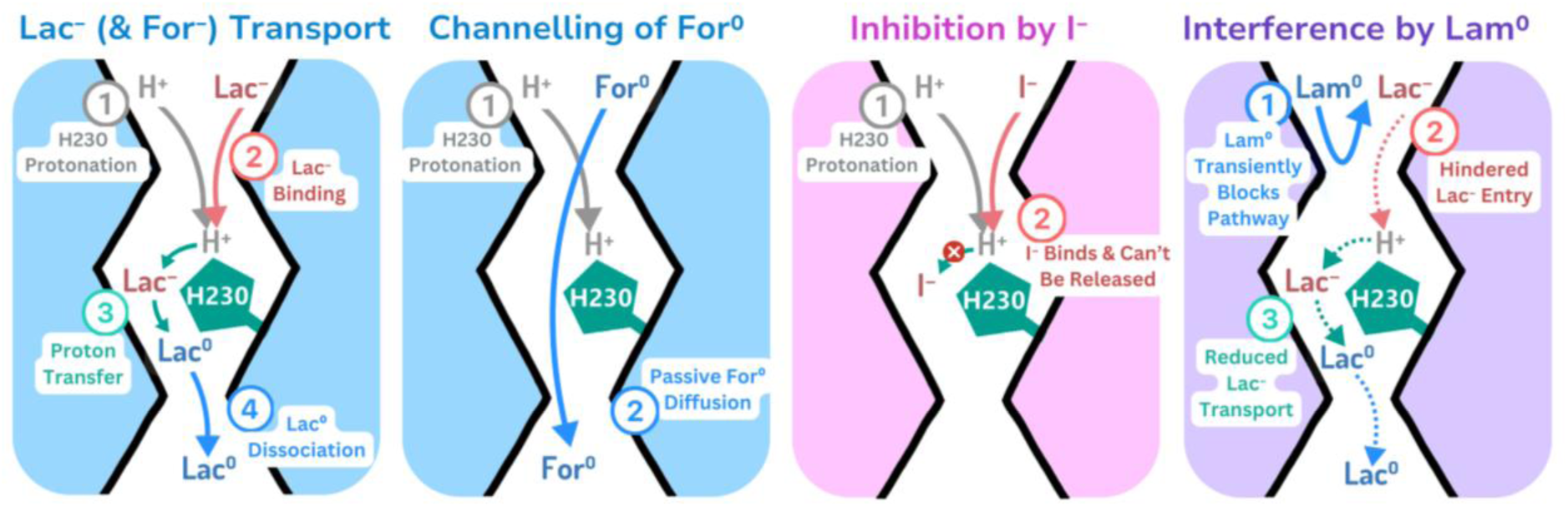
Proposed mechanisms for PfFNT-mediated Lac^-^ and For^-^ transport, channelling of For^0^, and inhibition by I^-^ and Lam^0^. For Lac^-^ and For^-^ transport, the steps of the proposed mechanism have been labelled 1 to 4. **1)** Neutral H230 is protonated via a Grotthuss mechanism and becomes positively charged. **2)** Either Lac^-^ or For^-^ enter the cavity. **3)** To destabilize the tight binding of the substrate within the cavity, charged H230 favourably donates a proton to bound substrate. **4)** Subsequently, the newly formed Lac^0^ or For^0^ molecule then dissociates from the transport cavity, completing the transport cycle. Channelling of For^0^ is a much simpler process – requiring only the protonation of H230 for For^0^ molecules to passively diffuse through the transport pathway. Similarly to Lac^-^ and For^-^, I^-^ binds to charged H230. However, as I^-^ cannot get protonated to be released from the cavity, I^-^ remains bound to charged H230 and inhibits PfFNT transport. Lam^0^ does not enter the cavity but transiently interacts with residues in the vestibules, hindering Lac^-^ entry and reducing transport.

### Evidence for the proposed transport mechanism

Our findings support a proton transfer mechanism in which Lac⁻ binds within the transport cavity and is protonated by H230 to allow release. Across our simulations, Lac⁻ consistently bound only when H230 was charged, whereas neutral Lac⁰ faced large energy barriers to enter the cavity and was more stable in the bulk solution. Further, we showed Lac^-^ bound tightly within the cavity and could not be released unless H230 transferred a proton to convert it to Lac^0^. Once formed, Lac^0^ readily dissociated toward either side of the membrane, consistent with PfFNT functioning as a bidirectional transporter.

Our proposed mechanism is supported by our experimental transport assays that indicate iodide inhibits PfFNT-mediated [^14^C]-lactate uptake in *X. laevis* oocytes (**Fig. 3c, f**). In combination with our computational observations that iodide only binds to charged H230 but cannot get protonated by H230 (**Table S5**), these results suggest a transport mechanism in which negatively charged compounds bind tightly to positively charged H230 and require proton transfer to dissociate from the cavity. However, our observation that iodide inhibits lactate uptake more strongly than formate at the same concentration does not fully exclude the possibility that iodide may bind to H230 and subsequently be released, despite our simulations suggesting this is unlikely to occur. Additional biochemical assays would thus help to confirm how iodide inhibits PfFNT-mediated lactate transport.

### pH dependent transport and H230 protonation

Our proposed proton transfer mechanism is consistent with previous reports of the pH dependence of PfFNT transport. Wu et al. (2015) observed that maximal PfFNT-mediated lactate/H^+^ transport activity occurs at pH 3.9 in a PfFNT-expressing strain of *S. cerevisiae*, in which the endogenous monocarboxylate transporters were knocked out. Although our H230 pK_a_ estimates vary, we expect pH 3.9 to be between the pK_a_s of lactate and H230 – suggesting transport occurs most efficiently at the pH at which there is the highest probability of H230 being protonated and lactate being deprotonated. At more acidic and more basic pH, transport rates decline, consistent with increased probability of both or neither species being protonated, respectively.

As pH 3.9 is well below the pH of the *P. falciparum* cytosol, this raises the important question of how H230 becomes protonated under physiological conditions to enable substrate binding. While it has been previously suggested that charged molecules (like protons) are unlikely to pass the hydrophobic constrictions of the PfFNT cavity^78^, several proteins exist that have hydrophobic gates along their transport pathways yet are known to conduct protons. These include the M2 proton channel from the influenza A virus^86^, the potassium and proton channel TMEM175^87^, the voltage-gated proton channel H_v_1^88,89^, and the otopetrin proteins Otop1 and Otop3^90^. In these proteins, it has been proposed that protons move along the transport pathway via a Grotthuss ‘proton-hopping’ mechanism – a mechanism that our DFTB simulations suggest also occurs in PfFNT (**Fig. 1g-h**). It is important to note that these DFTB simulations rely on the Born–Oppenheimer approximation, which assumes that the movement of atomic nuclei and their associated electrons in the system can be treated separately, as nuclei are generally much heavier and slower-moving than electrons.^91–93^ However, in proton transfer modelling, the proton (the hydrogen nucleus) has a small mass and the fact that its migration creates a transient excess-proton defect that requires ultrafast reorganisation of the surrounding water molecules’ electrons means that this separation of timescales may break down which is a limitation of DFTB in these conditions. .^91,92^ Nonetheless, we believe a Grotthuss mechanism remains the most plausible route for H230 protonation. While other computational approaches, such as *ab initio*^94^ or path-integral MD^95^, may more accurately represent proton behaviour, these simulations would likely be too computationally challenging to perform on a protein system of this scale.

During the trophozoite stage, cytosolic accumulation of lactate and protons likely provides a driving force for H230 protonation. Subsequent re-protonation of H230 after transport may then be facilitated by the cavity remaining wider and more hydrated following proton transfer, resulting in a lowered barrier for proton (and subsequently lactate) entry (**Fig. S9b**). As substrate binding and dissociation occur on the 0.2–1.5 μs timescale (**Fig. 2h**) and proton transfer from H230 to lactate occurs on a sub-picosecond timescale, the time taken for H230 to become protonated, more specifically the time for delivery of a proton from the bulk solution to the central transport cavity, is likely the rate-limiting factor for PfFNT transport.

### Conserved residues and transport mechanism

The transport mechanisms of human monocarboxylate transporters MCT1 and MCT4 involve proton transfer from a buried lysine that drives an alternating-access cycle.^96–99^ In contrast, our simulations indicate that PfFNT functions as a channel-like transporter without any major conformational changes. The residues that we and others have suggested are important for PfFNT-mediated lactate/H^+^ transport are highly conserved in FNTs from the other five *Plasmodium* species that cause malaria in humans, as well as in FNTs from related Apicomplexan parasites like *Toxoplasma gondii* (**Fig. S1c**), suggesting that these closely related FNTs may share the same proton transfer mechanism.

### PfFNT may alternate to functioning as a channel depending on the substrate and condition

Although PfFNT appears to function as a Lac⁻/H⁺ and For⁻/H⁺ transporter under physiological conditions, our simulations show channel-like permeation of formic acid when H230 is charged (**Movie S4**). In contrast, no permeation events were observed for the larger, neutral substrate Lac⁰ without mutating the constriction residues (**Fig. S6; Table S3**). Based on these observations, we hypothesised that PfFNT may be able to function as either a transporter or a channel depending on both the pH and the size and charge of the substrate being transported. Consistent with this hypothesis, while the larger, neutral molecule lactamide significantly reduced PfFNT-mediated lactate uptake in oocytes, the lack of difference in lactamide uptake between non-injected and injected oocytes from the LC-MS data indicates that lactamide is not appreciably being transported or channelled via PfFNT. These results suggest that, if PfFNT can channel neutral substrates, it likely does so only for small molecules like For⁰.

PfFNT-mediated For⁻ transport at neutral pH displays saturable kinetics^6^ – consistent with the protein functioning as a formate/H⁺ transporter. However, we propose that at more acidic pH PfFNT could function as a channel for formic acid. As the pK_a_ of both formate (3.75) and H230 (**Fig. S4**) suggest it is unlikely for both to be protonated except at conditions more acidic than the *P. falciparum* cytosolic pH range, it is unclear whether such a change in functionality could be physiologically relevant to the parasite. Nonetheless, the potential for PfFNT to change between functioning as a transporter and a channel further exemplifies the blurred boundary between these once-distinct classes of membrane transport proteins. Although the bacterial FNT StFocA was hypothesised to switch from a passive formate channel to an active formate/H^+^ symporter under acidic conditions^100^, there has not previously been any direct evidence to indicate StFocA or other members of the FNT family can switch between transporter and channel function. Here, we suggest PfFNT acts as a transporter without conformational changes yet can switch to functioning as a channel under acidic pH for small, neutral substrates. Given the high sequence similarity between PfFNT and FNTs from other *Plasmodium* species, as well as from closely related parasites such as *Toxoplasma gondii* (**Fig. S1d**), it is likely that other FNTs also have the capacity to switch between transporter- and channel-like modes.

### PfFNT as a drug target

Beyond its fundamental role at the channel–transporter interface, PfFNT is also an attractive antimalarial drug target due to its essential function in exporting lactate and protons from the parasite cytosol. Inhibitors such as MMV007839, MMV000972, and other structural analogues of these compounds have previously been shown to potently kill parasites and/or block PfFNT transport, however, resistance mutations to these compounds have already been identified, including G107S and V196L along the transport pathway and G21E in the N-terminal helix.^50,51,101^ Understanding which residues are essential for substrate binding and which may serve as potential resistance hotspots is therefore critical for rational inhibitor design. Our data show that binding sites for Lac⁻ and For^-^ overlap with one another and with the cryo-EM binding site of MMV007839, with common interactions at H230, V54, F94, and L104 (**Fig. S5b–d; S16a–b**). In contrast, several transport pathway lining residues surrounding the MMV007839 binding site do not have substantial interactions with substrates (**Fig. S16**). Among these residues are G107 and V196, sites of identified PfFNT resistance mutations. The other residues that fall into this category, Y31, I98, V200, and V220, may therefore represent potential sites where mutations could occur that confer resistance to PfFNT inhibitors and should be investigated in further studies.

### Conclusions: What is a channel, what is a transporter?

In this study, we present a mechanism for substrate transport in PfFNT that does not require any substantial conformational changes during the transport process. While we agree with the previous characterization of PfFNT as a transporter for lactate and H^+^ under physiological conditions, our data challenge the canonical definition of a transporter being a protein that undergoes conformational changes during transport. We propose instead that the key distinction between transporters and channels is that transporters require a cycle of changes – either in conformation or in a chemical change of protein and/or substrate (such as through proton transfer) - to enable the movement of substrates across membranes. In the case of PfFNT, transport of Lac^-^ and For^-^ is dependent first on H230 protonation, followed by proton transfer from H230 to enable release of the bound substrate. Subsequently, re-protonation of H230 is required for the next substrate to be transported. Comparatively, channels act like ‘on-off’ gates – channel activation creates a continuous pathway for rapid substrate conduction without requiring ‘resetting’ in between each permeation event. This is observed with For^0^ channelling in PfFNT, where permeation occurs following H230 protonation and does not require any other changes in conformation or protonation state in between conduction events - unlike what is seen when PfFNT is transporting substrates via the proton transfer mechanism. More broadly, our data indicate that rather than channels and transporters always being distinct classes of membrane transport proteins, single proteins like PfFNT can alternate between these two methods of transport, depending on factors such as pH and the substrate being transported.

Further, the difference in PfFNT transport kinetics observed across different studies^45,46,48^ warrants additional investigation into whether other factors, such as the lipid environment, may also be modulating whether PfFNT functions as a transporter or a channel. Recent studies have shown that some protein families include distinct members functioning as either channels or transporters, and even separate channel-like and transporter-like pathways within a single protein.^102–114^ However, this is the first example of a protein that (i) can be defined as a transporter, despite not undergoing conformational changes, and (ii) uses the same pathway to switch between transporter and channel modes depending on the substrate and environmental conditions. Both features challenge the commonly accepted definitions of transporters and channels, suggesting membrane transport mechanisms may not always be fundamentally distinct. By indicating that a protein can transition between channel-like and transporter-like mechanisms within a single pathway without mutation or substantial structural rearrangements, our findings blur a long-standing conceptual boundary in membrane transport – suggesting the evolutionary divide between channels and transporters may be more fluid than previously thought. We propose that, rather than dividing membrane transport proteins into channels and transporters, their mechanisms should be viewed as spanning a spectrum between these two formerly discrete modes of substrate transport. Recognising proteins like PfFNT that exist somewhere along (and perhaps even transition between) the transporter and channel ends of this spectrum may help reshape how membrane transport processes are classified, studied, and targeted therapeutically.

## Materials and Methods

### General Simulation System Set-Up

The Cryo-EM structures of PfFNT used in this study were obtained from the Protein Data Bank (PDB) for both the occluded (PDB ID: 7E26, 2.29 Å resolution) and inwards-open (PDB ID: 7E27, 2.29 Å resolution) conformations. Residues 1 to 6 of the N-terminal loop and residues 294 to 309 of the C-terminal helix of each subunit were not resolved in either structure and were not modelled in this study. Initial predictions of the ideal rotameric states of all histidine residues within the protein were performed using Schrodinger’s Maestro^115^ for both the occluded and inwards-open structures at a pH of 7.4. The inwards-open and occluded structures were then sourced from the Orientations of Proteins in Membranes (OPM) Database^116^ for use with the CHARMM-GUI^117,118^ membrane generator to create systems ready for simulation. All non-protein molecules were removed from the structures, including water molecules from both structures and the inhibitor MMV007839 from the inwards-open structure. The protonation states and rotamers of all histidine residues, except for H230, were set as predicted by Maestro^115^ (**Table 1**).

**Table 1.** Initial predictions of pK_a_ and protonation state for all histidines within PfFNT. Protonation states of each histidine residue were assigned from predictions of pK_a_ and rotameric state using Maestro. Mean pK_a_ for each residue is listed ± standard deviation (n=5).

| Residue | Predicted pK<sub>a</sub> (Maestro) | Predicted Protonation State |
| --- | --- | --- |
| 37 | 6.01 ± 0.00 | Neutral ($\delta$) |
| 59 | 3.28 ± 0.02 | Neutral ($\delta$) |
| 71 | 5.74 ± 0.00 | Neutral ($\epsilon$) |
| 159 | 6.23 ± 0.00 | Neutral ($\delta$) |
| 165 | 4.16 ± 0.01 | Neutral ($\delta$) |
| 179 | 6.28 ± 0.00 | Neutral ($\delta$) |
| 180 | 5.29 ± 0.01 | Neutral ($\epsilon$) |
| 230 | 3.51 ± 0.00 | Neutral ($\epsilon$) |
| 284 | 5.61 ± 0.01 | Neutral ($\epsilon$) |

For both the occluded and inwards-open structures, simulation systems were created where H230 was either neutral (protonated only at the ε nitrogen, or only at the δ nitrogen where specified otherwise) or positively charged (protonated at both the δ and ε nitrogen). In each simulation system, the whole PfFNT pentamer was embedded in a lipid bilayer comprised of pure 1-palmitoyl-2-oleoyl-sn-glycero-3-phosphocholine (POPC) lipids, as the precise lipid composition of the *P. falciparum* plasma membrane is not known. POPC is a commonly used model membrane in MD simulations, including simulations of other malaria transporters^119,120^ and related members of the FNT family^121–123^.

In our preliminary simulations of PfFNT without lipids inserted into the hydrophobic central hole, we noted the formation of a vacuum of space where no water or ions would enter. Using VMD, we generated structures where either one, two, or three lipids were inserted into the hydrophobic central hole in the inwards-open His^0^ system. MD simulations were run in triplicate for 100 ns per replicate. In all systems, there is minimal change in RMSD of the residues surrounding the central hole (**Fig. S2a-e**).

To maintain consistency with other MD simulation studies of FNTs, two POPC lipids were manually inserted into the central hole of the protein for all following simulations (**Fig. S2a-f**).^121–124^ Systems were solvated with water molecules and 150 mM KCl. K^+^ and Cl^-^ are expected to be the most abundant ions in the parasite cytosol, with estimated concentrations of 130-149 mM^125,126^ and 48 mM^127^, respectively. Total system dimensions were approximately 140 Å by 140 Å by 120 Å with around 222,220 atoms per system. Systems were visualized using the Visual Molecular Dynamics (VMD) software^128^.

Topology and coordinate files for each system were generated using TLEaP with the AMBER ff19SB protein forcefield, lipid17 lipid forcefield, OPC water model, and 12-6-4 ion parameters. ParmEd was used to repartition the masses of heavy atoms onto the hydrogen atoms to 3.024 Da for non-water molecules to allow a 4 fs timestep to be used. Subsequently, each system was prepared for MD simulations by (1) energy minimization, to eliminate steric clashes and unfavourable molecular interactions and allow the system to be brought to a local energy minimum, (2) heating, using the Langevin thermostat to slowly increase the temperature of the system to 310 K (36.85°C) over 500 ps, and (3) pressurizing, using the Monte Carlo barostat to maintain the system pressure at 1 atm over 200 ps, with the Berendsen barostat having been used for all prior preparatory phases. Throughout each of these preparatory phases, a 1 kcal/mol·Å^2^ harmonic restraint was kept on the α-carbon atoms of the protein, enabling lipids and water to accommodate the protein structure. Subsequently, restraints were incrementally released over a timeframe of 5 ns. All simulations were performed using AMBER20 (unless specified otherwise). Energy minimization and an initial heating step between -273.15 and -242.15 °C was performed using the pmemd.MPI parallel CPU implementation, while all other phases, including production, were performed using the pmemd.cuda GPU implementation. Hydrogen bonds were constrained with SHAKE. Electrostatic interactions were calculated using the Particle Mesh Ewald (PME) summation with a real-space cut-off of 10 Å. All system preparatory phases and production simulations were performed in triplicate for each system. K^+^ and Cl^-^ ions were prevented from entering the transport cavities to avoid potential competition with compounds using a spherical restraint of 1 kcal/mol·Å^2^ starting 15 Å from the centre of each cavity. Production simulations for each system without substrates present were run in triplicate for 1 μs per replicate. Additional simulations were performed for the His^+^ and His^0^ inwards-open systems, where a 1 kcal/mol·Å^2^ restraint was maintained on the α-carbon atoms of the protein backbone during production simulations. A summary of the main simulations performed in this study, including the number of replicates and the combined length of each simulation, is given in **Table 2**.

**Table 2.**
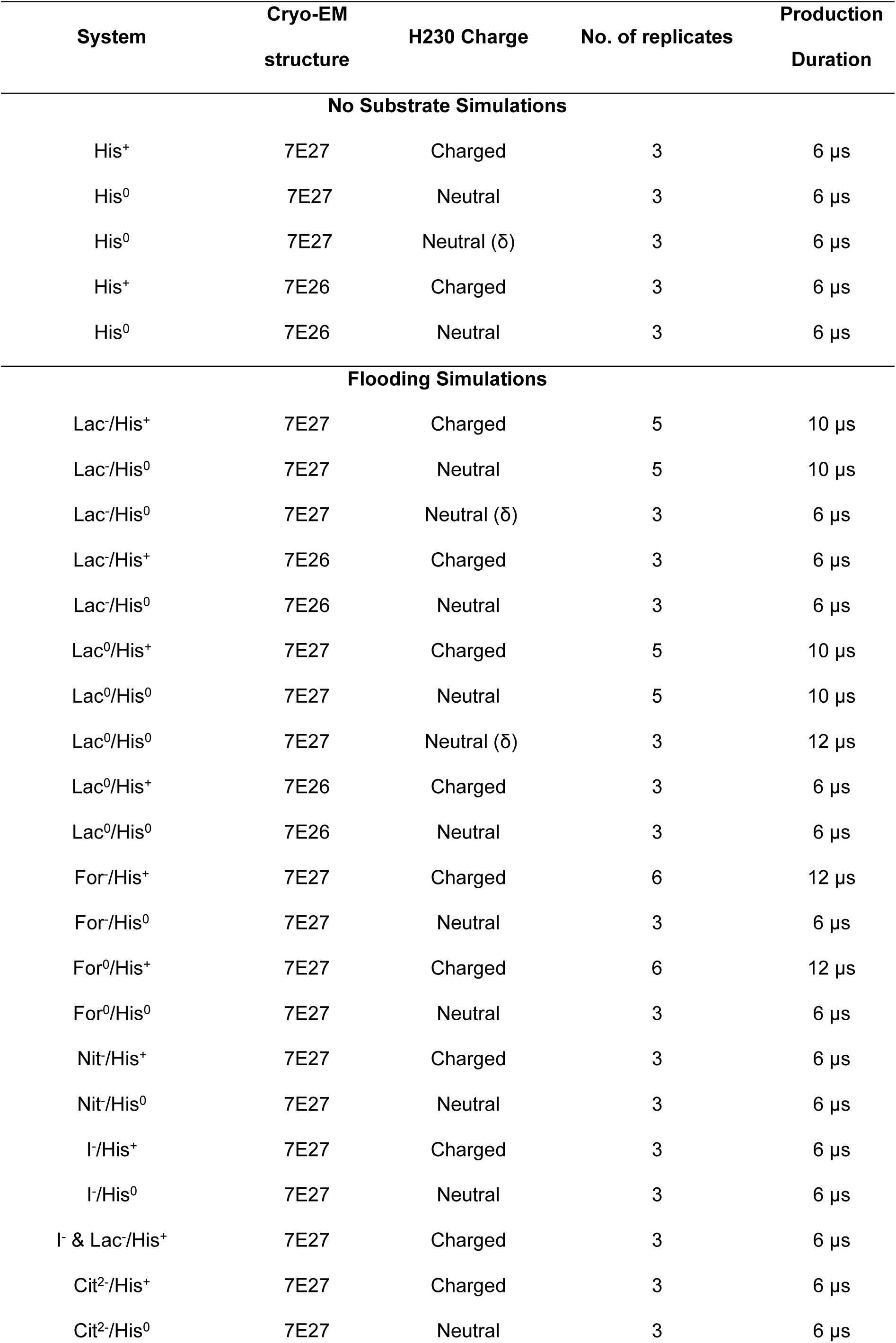
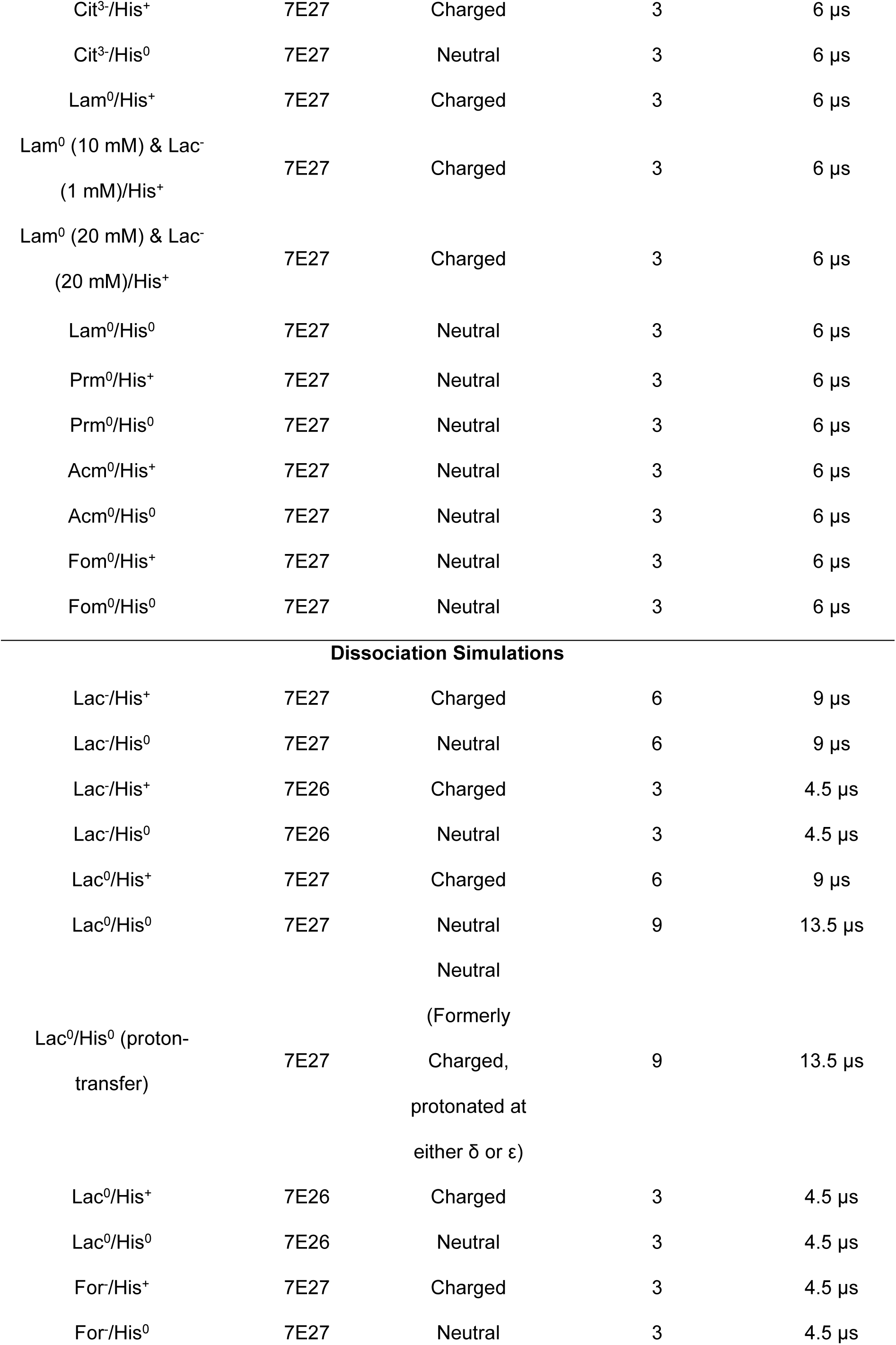
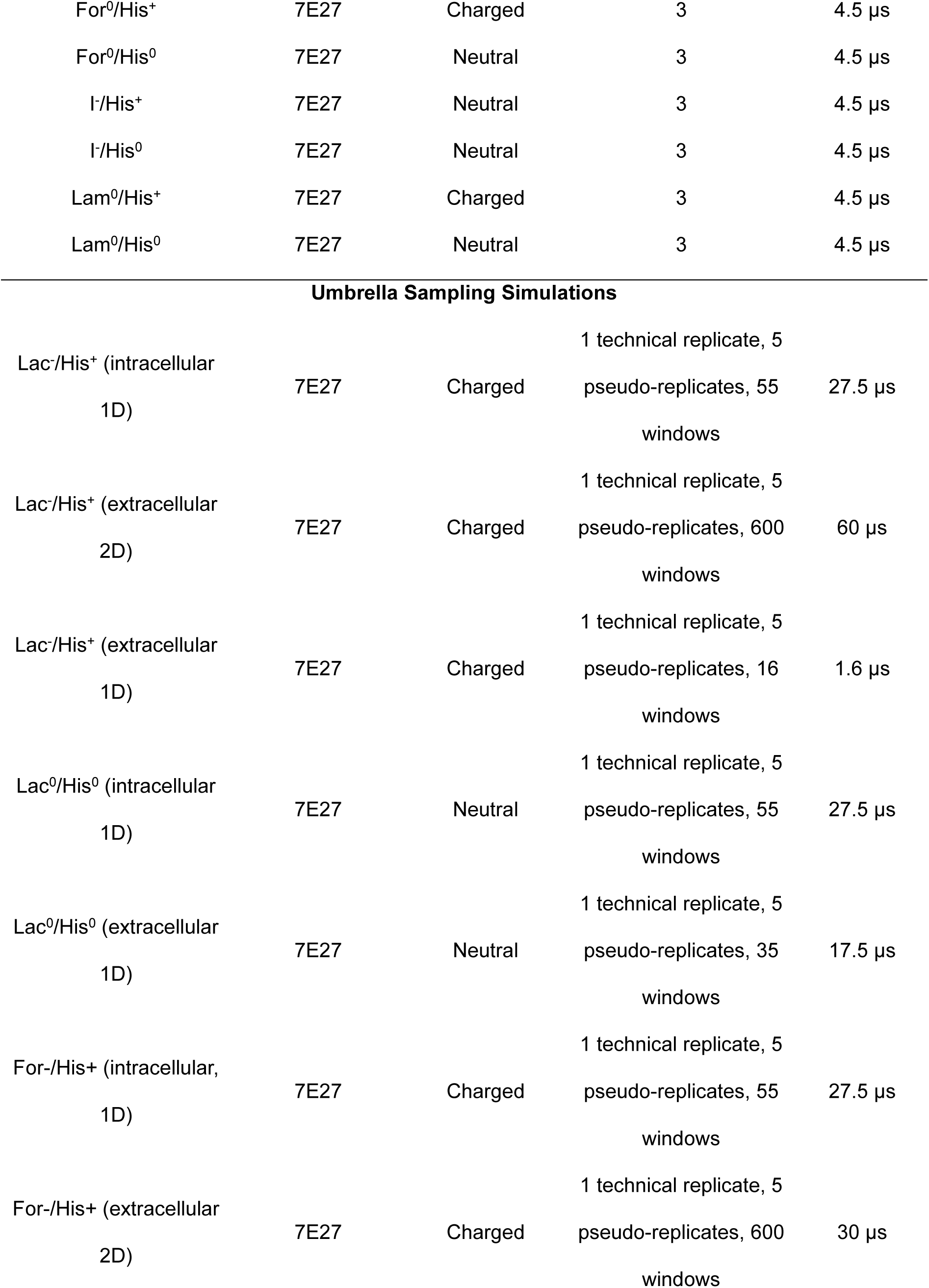
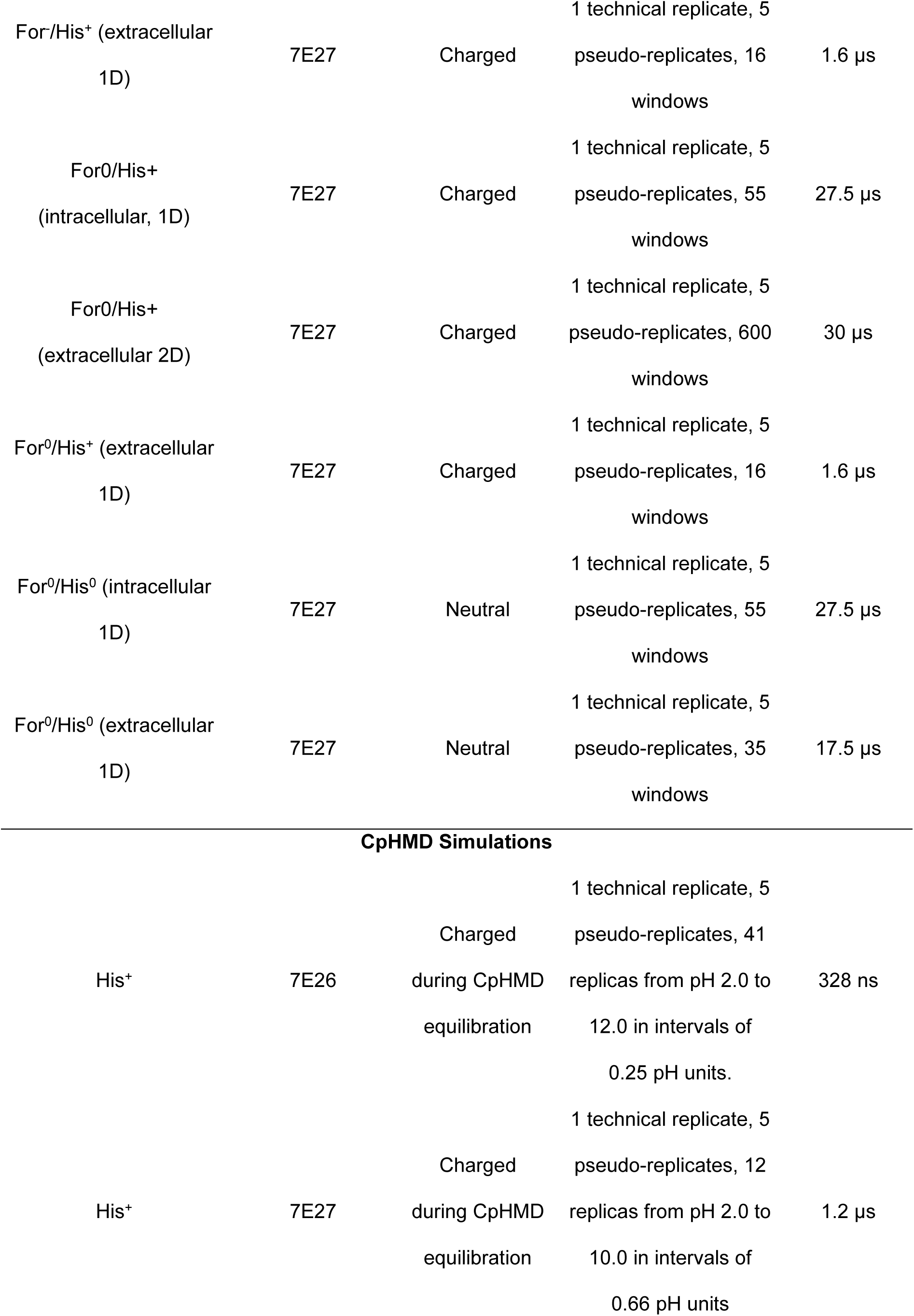
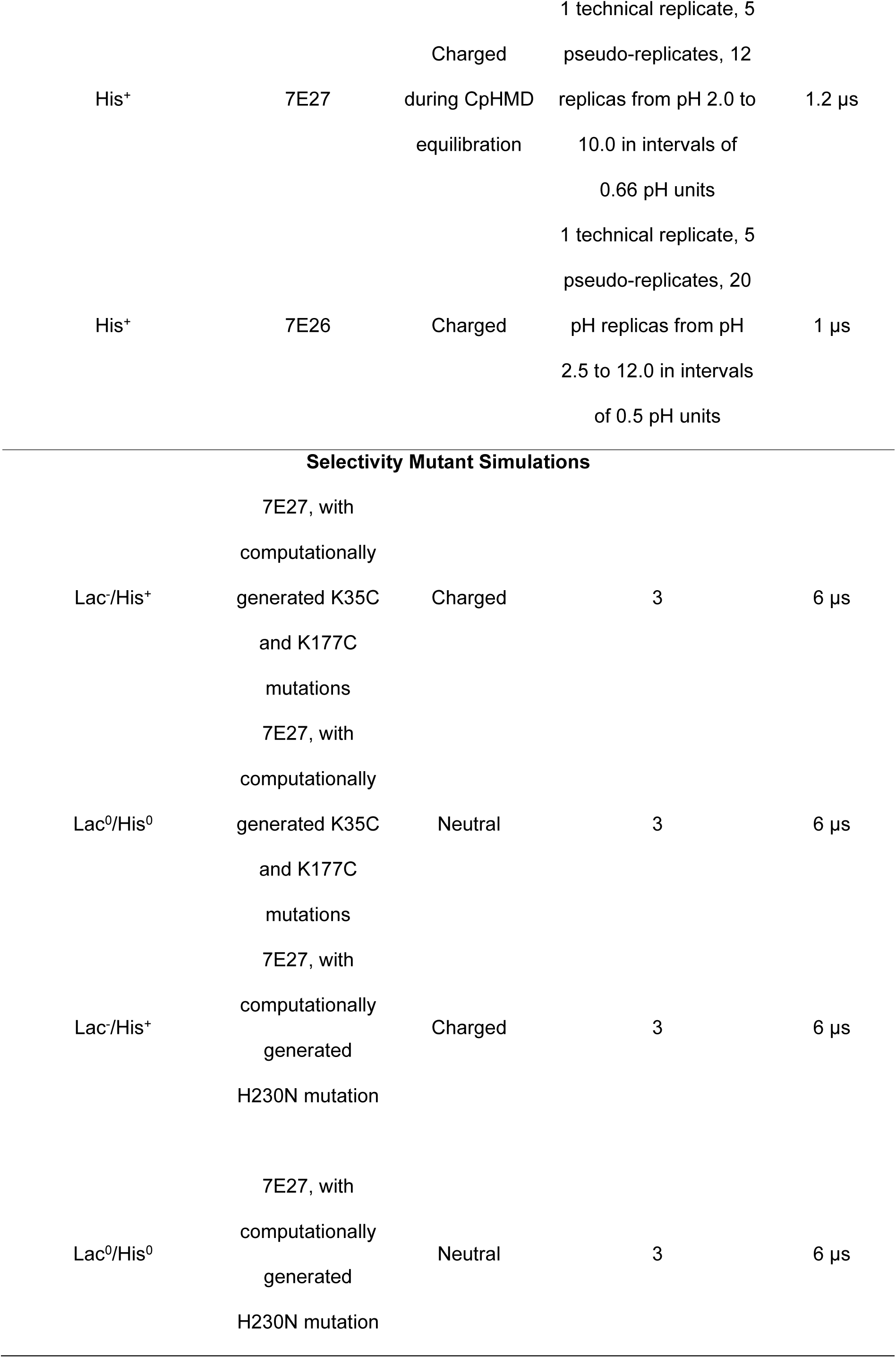
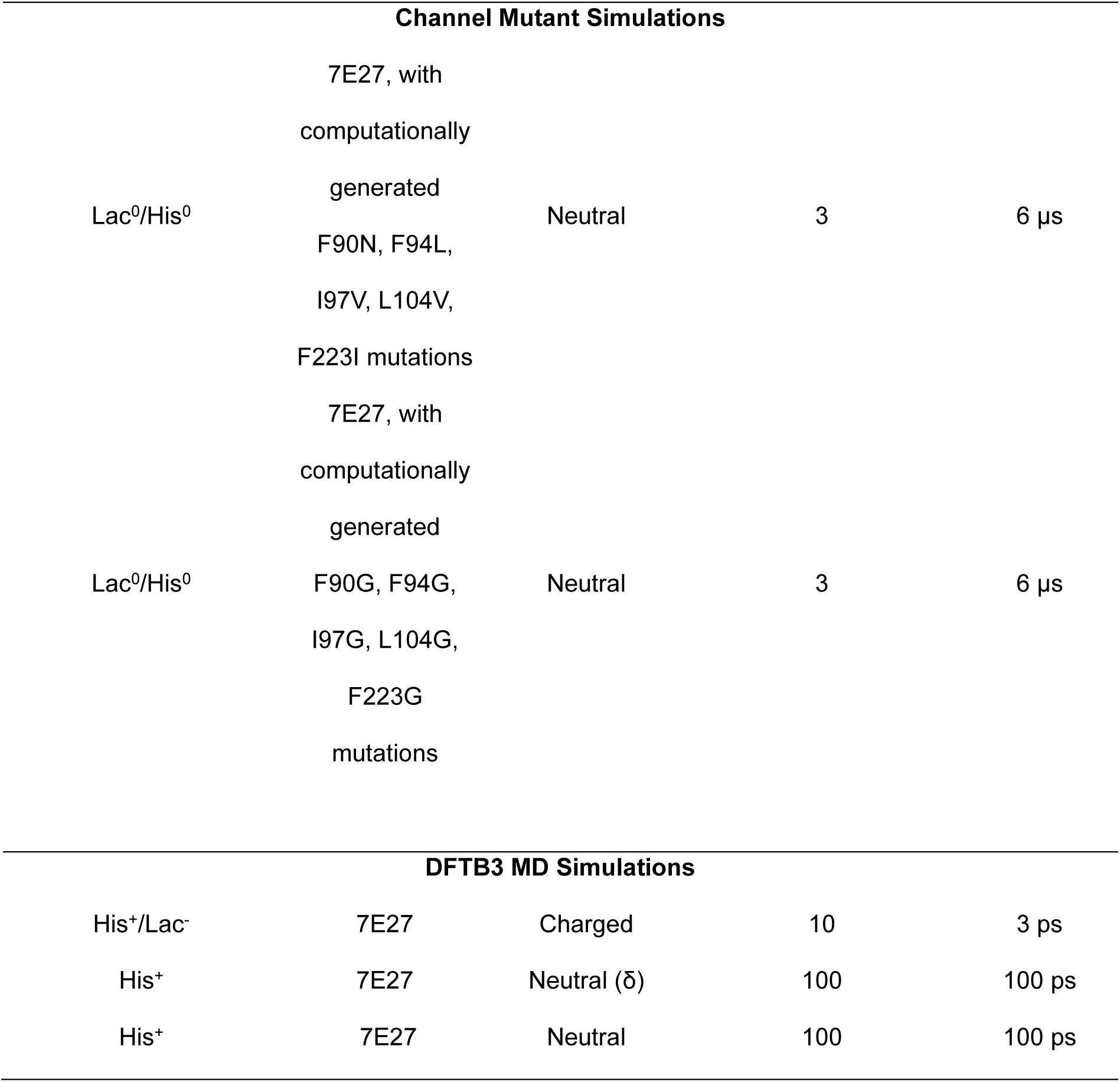
Main simulations performed in this study. A summary of the system, including the starting cryo-EM structure (inwards open = PDB ID: 7E27, occluded = PDB ID: 7E26), H230 charge, substrate charge (where applicable), number of replicates, and total production duration is given for the different simulations run in this study. Simulations are divided into categories based on the type of simulation being performed.

### Flooding Simulations

To investigate how substrate binding and transport occurs in PfFNT, we set up simulations with a high concentration of known or potential compounds with all possible combinations of H230 and compound protonation state (**Table 3**, **Fig. 5a**). The 3D molecular structures of L-lactate (C_3_H_5_O_3_^-^, CID 5460161), L-lactic acid (C_3_H_6_O_3_, CID 612), formate (CHO_2_^-^, CID 283), formic acid (CH_2_O_2_, CID 284), citrate^3-^ (C H O ^3-^), lactamide (C_3_H_7_NO_2_, CID 94220), propionamide (C_3_H_7_NO, CID 6578), acetamide (C_2_H_5_NO, CID 178), and formamide (CH_3_NO, CID 713) were downloaded from PubChem. The structure of citrate^2-^ (C_6_H_6_O_7_^2-^) was prepared by adding a proton to one of the tricarboxylate groups of citrate^3-^ using Molefacture plugin within VMD.^128^

**Table 3.**
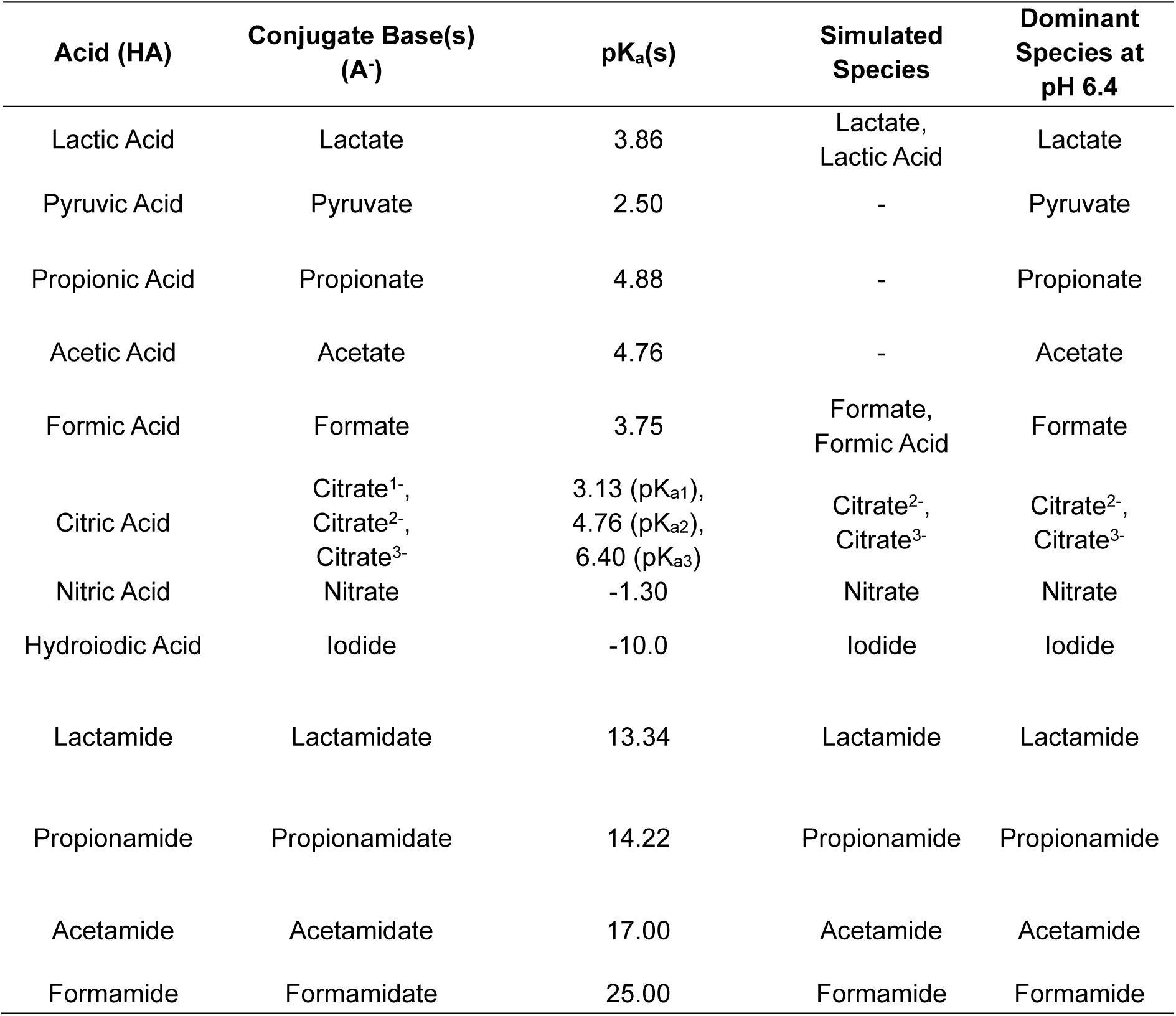
pK_a_ values of key compounds in this study. A list of the compounds, including the parent acid, conjugate base, pK_a_(s) of the acid form, the species simulated in MD, and the dominant species expected at pH 6.4 - the pH at which the *Xenopus laevis* transport assays were performed.

**Figure 5.**
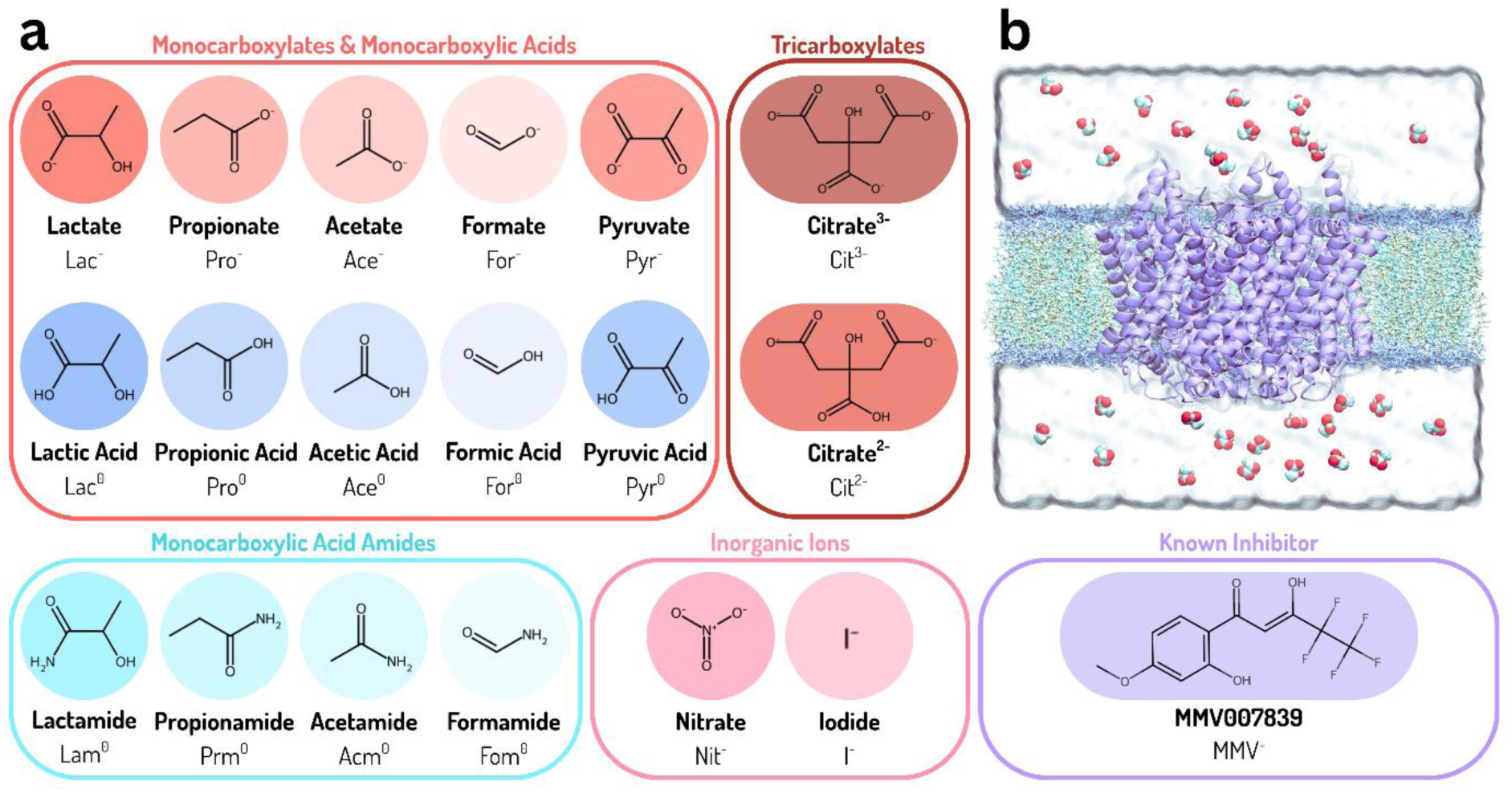
Overview of flooding simulations. **A)** Structures of compounds tested computationally and experimentally in this study. **B)** Visualization of a lactate flooding simulation system with 30 substrates randomly distributed across both sides of the membrane. Lactate molecules are shown in a Van der Waals representation and are coloured by atom type, cyan for carbon, red for oxygen, and white for hydrogen. The PfFNT pentamer is shown in a purple cartoon representation, POPC lipids are shown in licorice, and space occupied by water and ions by the grey surface.

The structure file for each compound was converted to PDB format using OpenBabel^129,130^, before being parameterized using Antechamber and the General AMBER Forcefield, GAFF2.^131^ The structure and parameters for nitrate (NO_3_-) were obtained from quantum chemical calculations performed by Baaden et al.^132^ Each compound was simulated in a vacuum following parameterization to ensure the structures were stable. Briefly, each compound’s structure was energy minimized as a lone molecule in a 40 Å × 40 Å × 40 Å box prior to heating to 310 K over 500 ps using SANDER under periodic boundary conditions with a cut-off of 10 Å. Following heating, structures were simulated using PMEMD with a 2 fs timestep for 5 ns without periodic boundary conditions using a cut-off of 9999 Å.^133^ Hydrogen bonds were constrained using SHAKE.

The final frame from the equilibration simulations of systems without substrates was used as a starting coordinate for flooding simulations. The AMBERTools ‘AddToBox’ program was used to randomly distribute ∼20 mM of the chosen compound (e.g., 30 molecules of lactate) throughout the system (**Fig. 5b**). Any overlapping water molecules where compounds were inserted were removed. TLEaP’s ‘AddIonsRand’ command was used to randomly insert ions to restore the charge of the system to neutral and maintain ion concentrations around 150 mM for all systems. Systems underwent the same preparatory procedure described above of (1) minimization, (2) heating, (3) pressurization, and (4) relaxing of restraints on the protein backbone, before starting production simulations. All steps of simulation set-up and production were performed using three to six replicates, with production simulations lasting 2 μs per replicate (see **Table 1**).

Analysis of substrate binding events was performed using ProLIF^134^. Frames from each binding event were aligned and different subunits were superimposed into an ensemble trajectory of a single subunit used for ProLIF analysis to create an average interaction map. Free-energy surfaces (FES) were created for each simulation system using python scripts adapted from Tanner et al.^135^ For each system, an ensemble trajectory of the whole PfFNT pentamer was made by combining all trajectory frames from all replicates. In cases where both the inwards-open and occluded structures were used as starting points, the data were combined for analysis. The 3D space of each system was divided into a series of grid points, and the probability of a ligand occupying any of those grid points (p_grid_) was calculated. Following calculation of p_grid_ for each grid point, the data were converted to a measure of the free energy (kJ/mol) of the substrates across the system using the following equation:

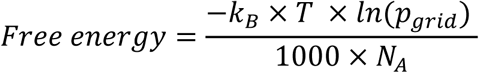

Where k_B_ is the Boltzmann constant, T is the simulation temperature (310 K), and N_A_ is Avogadro’s number.

### Dissociation Simulations

To examine the ability of different compounds to dissociate from the cavity we set up simulations with each compound bound in the cavity for each H230 protonation state. To do this, we extracted the final frame of each Lac^-^, Lac^0^, For^-^, For^0^, I^-^, and Lam^0^ flooding system and manually inserted a compound into the transport cavity of each subunit. Any compounds not within the transport cavities were deleted, and ions were added to maintain a charge-neutral system. Additional 1 kcal/mol·Å^2^ harmonic restraints were placed on the sidechain atoms of the protein as well as on the ligands to allow the substrate to equilibrate in the cavity. Restraints were released by first incrementally reducing the restraints on the protein sidechains for 25 ns and then incrementally reducing the restraints on the protein backbone over a further 25 ns, keeping the restraints on the ligand the whole time. To ensure the substrates started in energy-minimum binding positions in the cavity, a simulated annealing process was performed in triplicate for each system involving rapidly heating the system from 310 to 360 K over 5 ns before gradually cooling the system down to 310 K for 50 ns. Following annealing, the restraints on the ligand were gradually released for 5 ns. Production simulations were then run in triplicate for 1.5 μs per replicate to observe whether substrate dissociation occurred.

To mimic the effect of proton transfer, the final frames from the Lac^-^/His^+^ flooding simulations where binding events occurred and simulations where lactate was inserted into the transport cavity were extracted using CPPTRAJ^136^, and the PDB files were edited to rename the proton on the δ or ε nitrogen of H230 as if it belonged to lactate. The system was then re-parameterized using the parameters for lactic acid instead of lactate. Additional 1 kcal/mol·Å^2^ harmonic restraints were placed on the sidechain atoms of the protein as well as on the ligand. Restraints were released by first incrementally reducing the restraints on both the ligand and the protein sidechains for 25 ns, and then incrementally reducing the restraints on the protein backbone over a further 25 ns. Production simulations were run in triplicate for 1.5 μs per replicate to observe whether substrate dissociation occurred.

### Steered MD and Umbrella Sampling Simulations

To generate starting coordinates for umbrella sampling (US), steered MD (SMD) simulations were performed for the Lac^-^/His^+^, Lac^0^/His^0^, For^-^/His^+^, For^0^/His^+^, and For^0^/His^0^ systems. For each system, the final frame of one replicate of the simulations where substrates were inserted into the cavity was used as a starting frame for SMD. As the PfFNT transport cavity is narrow, SMD was performed in four individual ‘pulling’ events – intracellular bulk to intracellular constriction, cavity to intracellular constriction, cavity to extracellular constriction, and extracellular bulk to extracellular constriction - rather than one continuous pulling event to prevent distortions to the constriction residues. For both SMD and US, the collective variable used for the intracellular side of the protein was the distance between the centre of mass of the backbone atoms of extracellular constriction and the centre of mass of the substrate. Similarly, the collective variable used for the extracellular side of the protein was the distance between the centre of mass of the backbone atoms of the intracellular constriction and the centre of mass of the substrate.

For all SMD, a force constant of 25 kcal/mol·Å^2^ was used to move the substrate in 0.5 Å increments every 2 ns in all five subunits simultaneously. AMBER restart files were written every 2 ns of simulation to obtain starting coordinates for US along the pathway from the cavity to bulk. Before starting US simulations, simulated annealing was run on each of the restart files from the SMD with the 25 kcal/mol·Å^2^ restraint maintained throughout all heating and cooling phases to relax the protein and substrate. For the Lac^0^/His^0^ and For^0^/His^0^ systems, two separate one-dimensional US simulations were run to sample the entirety of the transport process from bulk to bulk using a combined total of 90 windows. Each window was simulated for 500 ns. All windows were spaced 0.5 Å and used a force constant of 15 kcal/mol·Å^2^, except for eight windows near the intracellular constriction where extra windows were inserted where necessary to ensure sufficient overlap and were restrained with a 50 kcal/mol·Å^2^ force constant. For the Lac^-^/His^+^, For^-^/His^+^, and For^0^/His^+^ systems, two separate one-dimensional US and one two-dimensional US were used to sample the entirety of the subunit from bulk to bulk. The one-dimensional intracellular US used the window spacing and force constants as used for the Lac^0^/His^0^ and For^0^/His^0^ systems, however, the extracellular one-dimensional US only used 16 windows covering 8 Å of space in each subunit from bulk on the extracellular side into the vestibule above the extracellular constriction.

The remaining space – cavity to vestibule – was sampled using two-dimensional US where an additional collective variable was added to bias the χ1 dihedral angle of H230 to capture whether the sidechain of H230 ‘swings’ down towards the extracellular constriction when lactate approaches the cavity. For each substrate window, a short SMD was performed where the H230 χ1 dihedral angle was moved in 0.25 rad increments every 0.5 ns from the sidechain pointing ‘up’ towards the intracellular constriction to ‘down’ towards the extracellular constriction, using a force constant of 50 kcal/rad Å^2^ to create starting coordinates for 2D US. In total, 25 substrate windows spaced 0.5 Å apart along the reaction coordinate between the cavity and the extracellular vestibule were used for 2D US, with each substrate position having 24 χ1 windows sampling the H230 sidechain movement – giving a total of 600 individual 2D US windows. Each window was simulated for either 50 ns (For^-^/His^+^ and For^0^/His^0^ systems) or 100 ns (Lac^-^/His^+^ system). All substrate restraints in steered MD, 1D US, and 2D US were set up using the AMBER ‘COM_DISTANCE’ restraint. H230 χ1 restraints were set up using the AMBER ‘TORSION’ restraint. Steered MD and umbrella sampling codes were activated by setting the non-equilibrium free energy method (infe) to 1 in the input file.

Similarly to 1D US, 2D US was performed with five pseudo replicates by simultaneously performing 2D US in each of the five subunits. All US data were unbiased, and the potential of mean force (PMF) was determined using a version of the Weighted Histogram Analysis Method (WHAM) implemented by Alan Grossfield. For all 1D US simulations, nonperiodic WHAM calculations were performed with 500 bins and a convergence tolerance of 0.00000001. The minimum and maximum bounds of the histogram were set to the respective minimum and maximum distances of the substrate along the reaction coordinate. To assess whether the PMF has converged and determine how much equilibration time can be discarded from the start of the simulations, we recalculated PMFs for each subunit after incrementally removing data from either the start or end of the simulation. For all 1D umbrella samplings, the first 100 ns of data was removed, and the data collected between 100 and 500 ns were used to generate the final PMFs (**Fig. S7, S10, S11, S12**). To obtain 2D PMFs, the 2D implementation of WHAM was used to analyze data collected between 10-50 ns (For^-^/His^+^ and For^0^/His^0^ systems) or 10-100 ns (Lac^-^/His^+^ system) from 2D US simulations. For the H230 χ1 dihedral angle reaction coordinate, a periodicity of 2*pi was specified, while the reaction coordinate for lactate was treated as nonperiodic. In both dimensions, calculations were performed with 50 bins, and the minimum and maximum bounds of the histogram were set to the minimum and maximum values along the respective reaction coordinates. The 2D PMFs obtained from each subunit are shown in **Fig. S7** for the Lac^-^/His^+^ system, **Fig. S10** for the For^-^ /His^+^ system, and **Fig. S12** for the For^0^/His^+^ system. Average 2D PMFs were created by combining data from all five subunits prior to running WHAM. The minimum free energy pathways (MFEP) were extracted from each of the five 2D US PMFs using a two-basin potential implementation of the string method developed by Weinan et al.^137^ and were averaged to obtain a 1D PMF.

### Simulations of computationally generated mutant structures

Several mutant structures of PfFNT were computationally generated using AMBERTOOLS’ PDB4AMBER to investigate (i) mutations that abolish lactate binding (selectivity mutants) and (ii) mutations that could convert PfFNT into a channel for lactate or lactic acid (channel mutants).

Selectivity mutants were chosen based on previous mutagenesis studies in PfFNT and other FNTs, which highlighted the importance of lysine residues above and below the cavity and the central histidine for substrate transport. Two selectivity mutant structures were prepared: K35C/K177C and H230N.^124,138,139^ Each was subjected to triplicate flooding simulations with either lactate or lactic acid for 2 μs, using the same simulation set-up protocol described above for wild-type PfFNT.

Two channel mutants, PfFNT-F94I/I97C/L104V/F90N/F223I (chosen by analysing natural variants in constriction residues across the FNT family) and PfFNT-F94G/I97G/L104G/F90G/F223G were computationally generated for running flooding simulations with Lac^0^. In all simulations, H230 was neutral. Flooding simulations were run in triplicate for 2 μs per replicate. All simulation set up was performed as described above for PfFNT-WT systems. Analysis of permeation events was performed using a script modified from Adamson et al.^140^

### Quantum chemical proton transfer calculations

To determine the energetic favourability for H230 to donate a proton to various substrates (lactate, formate, nitrate, and the known inhibitor MMV007839) and identify potential mechanisms by which H230 can become protonated, we used quantum chemical calculations. The initial geometries for each substrate were extracted from the top-ranked cluster of frames of the combined trajectories for all binding events. Where possible, frames were taken for both δ NH and ε NH binding events. To investigate H230 protonation we took random snapshots from trajectories where H230 was charged and water was present within the transport cavity, and transferred a proton from H230 to the nearby water molecule. The coordinates for H230 and the bound substrate/water molecule were then extracted from each structure and processed using a Python script with PyMOL^141^ and RDKit^142^ to protonate both the carbon and the nitrogen atom of H230 that would otherwise be involved in a peptide bond to maintain a charge-neutral backbone.

Each unique dimer structure between the substrate and histidine was reoptimized with the SOGGA11-X functional^143^ with third order Grimme dispersion^144^ and Becke-Johnson Damping^145^ (D3(BJ)), using the Def2-TZVPP basis set^146^ in the Gaussian 16 program.^147^ SOGGA11-X was chosen as it has been shown to be a versatile hybrid functional method for barrier heights.^148^ The optimizations were repeated gas-phase, or with a water, methanol (MeOH), ethanol (EtOH), or nonanol polarisable continuum medium. For many substrate-histidine dimers, a spontaneous proton transfer occurred in the gas-phase. The geometry of these optimized gas-phase structures were reoptimized using an implicit solvent. The coordinates from the solvent-optimized structures with the proton on either H230 or the substrate were used as the “reactant” and “product” for 2-Structure Synchronous Transit-Guided Quasi-Newton Method (QST2) calculations^149^ to find a transition state. Where a transition state was able to be found, an intrinsic reaction coordinate calculation (IRC)^150,151^ was performed to observe the transition state energy barrier.

### Density functional tight binding (DFTB) molecular dynamics simulations

To investigate whether proton transfer can occur within the cavity, we extracted frames of lactate bound at either δ or ε NH groups where the H230 sidechain was either pointing ‘down’ or ‘up’. Similarly, to determine whether H230 protonation could occur within the cavity, we extracted frames where water was present within the cavity and H230 was charged. In the latter case, a proton from H230 was transferred to 100 distinct locations spread throughout the cavity for each of the δ or ε NH. For all frames, the coordinates for all protein, substrate, and water molecules within 5 Å of H230 were obtained and processed to protonate the carbon and nitrogen atoms of H230 that would otherwise be involved in the peptide bond without the 5 Å cut-off. Using each of these structures, MD simulations were performed using 3rd-order density functional tight binding (DFTB3)^152^, which was computed on-the-fly at each timestep using Grimme’s D3 dispersion^144^ with Becke-Johnson damping^145^ (D3(BJ)) and the 3ob-3-1 parameter set^153^ made for organic and biological systems. All DFTB simulations were performed using the DFTB+ software package (v. 23.1)^154^.

### CpHMD Simulations

To estimate the pK_a_ of H230, we performed Continuous Constant pH MD (CpHMD) simulations using the GPU-accelerated generalized Born (GB) based asynchronous pH replica exchange CpHMD method implemented in AMBER18, and the all-atom Particle Mesh Ewald CpHMD methods implemented in AMBER24 and GROMACS (v.2021).

For the AMBER18 CpHMD simulations^71,72^, the starting structure was obtained from our MD simulations of the ‘occluded’ structure (PDB ID: 7E26) with neutral H230 and was prepared for simulation using the CpHMD preparation script developed by the Shen Lab^155^. All glutamate, aspartate, and histidine residues were set to be titratable, and residues were changed to the GL2, AS2, and HIP protonation states. The system was energy minimized for 1000 cycles, with a 50 kcal/mol·Å^2^ restraint held on all non-hydrogen atoms. Subsequent equilibration simulations were performed using a 2 fs timestep, where restraints on the protein were incrementally released over a period of 4 ps. Simulations were performed using a PME cut-off of 999 Å. All simulations were performed at 310 K with an implicit salt concentration of 150 mM using the ff14SB protein forcefield and the GBNeck2 generalized Born (GB) implicit-solvent model. Hydrogen bonds were constrained using SHAKE. During equilibration, the implicit pH was set to 7.0. Following equilibration, CpHMD-pHREX replicas were simulated at 41 different pH conditions ranging from pH 2.0 and pH 12.0, spaced 0.25 pH units apart using the CpHMD-pHREX script developed by the Shen Lab^156^. Exchanges were attempted between replicas every 1000 steps, i.e., 1 ps. Each replica was simulated for a total of 8 ns. Simulation convergence was assessed by measuring the change in unprotonated fraction (i.e., the percentage of frames where the residue is unprotonated out of the total number of frames) for H230 at each pH replica over time until substantial changes could no longer be observed with increasing simulation length. Once converged, the predicted pK_a_ for H230 from each subunit was determined by fitting the unprotonated fraction value from each simulated pH condition to the generalized Henderson-Hasselbalch equation to obtain the best-fit titration curve. The overall exchange frequency between all replicas was 24.92% (**Fig. S11a-f**).

For CpHMD simulations with AMBER24^73,157,158^, starting structures from both the ‘inwards-open’ (PDB ID: 7E27) and ‘occluded’ (PDB ID: 7E26) conformations were used for simulation. To minimize computational cost, only one subunit from each system was simulated. Systems were prepared using the CHARMM-GUI Constant-pH Simulator tool.^159^ Each subunit was embedded in a 60.5 × 60.5 Å model POPC bilayer and was solvated with 150 mM NaCl. Total system dimensions were 60.5 × 60.5 × 112.8 Å, and consisted of ∼38,315 atoms. All glutamate, aspartate, and histidine residues were set to be titratable. Systems were energy minimized for 5000 cycles with 10 kcal/mol·Å^2^ and 2.5 kcal/mol·Å^2^ restraints placed on the protein backbone and lipid headgroups, respectively. Equilibration simulations were performed where the restraints on the protein backbone and lipid headgroups were incrementally released over a period of 2 ns. These simulations were run using a 1 fs timestep for the first 500 ps, before switching to a 2 fs timestep for a remaining 1.6 ns. Only the sidechain of F94 remained restrained after equilibration, using a 5 kcal/mol·Å^2^ restraint. All simulations were performed at 310 K, using a PME cut-off of 9 Å. Simulations were parameterized using the ff14SB protein forcefield, TIP3P water model, and lipid21 lipid forcefield. Each system was simulated using 12 different pH windows – pH 2.00, 2.73, 3.45, 4.18, 4.91, 5.64, 6.36, 7.09, 7.82, 8.55, 9.27, and 10.00. Each window was simulated for 100 ns.

For CpHMD simulations run using GROMACS, a specialized version of GROMACS (v.2021) was used with constant pH MD module implemented.^160^ For CpHMD simulations, the modified CHARMM36 forcefield^161^ was used to parametrize interactions of the system. λ-dynamics based constant pH MD was applied to interpolate the hamiltonians of the protonation states of titratable residues. Multisite representation was used for histidine residues. Steepest descent energy minimization was conducted followed by six sequential steps of equilibration (112 ns in total) with a gradual decrease in the restraining force applied to protein backbone and sidechain atoms. Protein, lipids, and ion-water groups including buffer particles were treated independently to increase accuracy. Buffer ions were added into system by replacing water molecules for maintaining the charge neutral condition during simulation, the minimum distance between protein/lipid atoms and buffer ions was 1.0 nm, and those buffer ions were fixed in space during titration calculation. The pH range of the simulation was from 2 to 10.5 with an interval of 0.5; three independent replicas under each pH condition were run for 50 ns. The Henderson–Hasselbalch equation was used to calculate the pKa values of titratable residues.

### General simulation analysis

AMBER trajectories were post-processed using AMBERTools’ CPPTRAJ.^136^ Unless specified otherwise, all analysis was performed in Python using MDAnalysis^162^, with the use of the NumPy^163^ and Pandas^164^ libraries. All graphs and figures were created using Matplotlib^165^. Statistical analysis was performed using the SciPy library^166^. Unless specified otherwise, statistical significance was determined using one-way or two-way ANOVAs with *post-hoc* Tukey tests. The radius throughout the transport cavity was measured for each system using HOLE2.0 implemented in MDAnalysis.^167^ For each system, an ensemble trajectory was created by first aligning all trajectories from the three replicates of each system together and then superimposing the five subunits onto each other to create a single subunit trajectory. The CVECT function [0,0,1] was used to find pathways along the Z-axis through the centre of each subunit. The pK_a_ of H230 was predicted from different systems using PROPKA3.0.^68^ Cluster analysis was performed using the WMC PhysBio clustering plugin in VMD.^168^ All 2D chemical structures presented throughout were drawn using RDKit within python.^142^

#### Compounds used in competition assays

MMV007839 and lactamide were purchased from MolPort, and all other compounds were obtained from Sigma-Aldrich. The test molecules citrate, pyruvate, propionate, acetate, formate, nitrate, and iodide were added as sodium salts, whereas lactamide, propionamide, acetamide, formamide, and MMV007839 were pure compounds. Initial 1 M stock solutions of each compound (excluding MMV007839) were prepared in Milli-Q water and subsequently diluted to a final concentration of 1 mM (for lactate) or 10 mM (for all other test compounds) in ND96 buffer (96 mM NaCl, 2 mM KCl, 1 mM MgCl₂, 1.8 mM CaCl₂, 10 mM MES, and 10 mM Tris-base; pH 6.4 adjusted with NaOH). For MMV007839, a 200 μM stock solution was prepared in DMSO and diluted to 2 μM in ND96. The final concentrations of Milli-Q water or DMSO in the assay solutions were approximately 1%. All final solutions in ND96 had a pH of approximately 6.4.

#### Ethics approval for working with *Xenopus laevis* frogs

Ethical approval of the work performed within this study with adult female *Xenopus laevis* frogs was obtained from the Australian National University (ANU) Animal Experimentation Ethics Committee (Animal Ethics Protocol Number A2023/24) in accordance with the Australian Code of Practice for the Care and Use of Animals for Scientific Purposes.

#### Preparation of wildtype and HA-tagged PfFNT cRNA for *X. laevis* oocyte experiments

The wildtype (3D7) untagged *pffnt* oocyte expression vector was previously prepared by Marchetti et al. (2015). Plasmid DNA was extracted using the GeneJET Plasmid Miniprep Kit. The *pffnt* insert was sequenced using the T7 forward and reverse primers to confirm the presence of full-length wildtype PfFNT. Sequencing was performed on an AB 3730xl DNA Analyzer (at the Genome Discovery Unit - ACRF Biomolecular Resource Facility, The John Curtin School of Medical Research, Australian National University) following the manufacturer’s protocol (Applied Biosystems 2002).

Plasmid DNA was linearized with the NotI restriction endonuclease (New England Biolabs), and successful linearization was verified by agarose gel electrophoresis, which showed a single band at the expected size. The linearized DNA was purified by phenol:chloroform:isoamyl alcohol (25:24:1) extraction and used as a template for in vitro cRNA synthesis with the mMESSAGE mMACHINE T7 Transcription Kit. The resulting cRNA was purified again using phenol:chloroform:isoamyl alcohol extraction and quantified on a nanodrop spectrophotometer before injection into *Xenopus laevis* oocytes.

#### Selection and microinjection of *X. laevis* oocytes

*X. laevis* oocytes were surgically removed from adult female frogs via an abdominal incision, and were digested for 8–10 hours at 18 °C in 0.5 mg ml⁻¹ collagenase B (Roche) prepared in Ca²⁺-free oocyte resuspension buffer (96 mM NaCl, 2 mM MgCl₂, 1.8 mM KCl, 1.9 mM Na₂HPO₄, 5 mM HEPES, and 50 µg ml⁻¹ gentamycin; pH 7.8 adjusted with NaOH). Following digestion, oocytes were washed extensively and transferred to resuspension buffer supplemented with 1.8 mM CaCl₂. Stage V–VI oocytes were then manually selected for experiments. PfFNT cRNA (30 ng) was microinjected into oocytes within 24 hours post-surgery, consistent with prior studies.^45,51,169^ Oocyte injections were performed using a Micro4 micro-syringe pump controller and A203XVY nanolitre injector (World Precision Instruments, Sarasota, FLA, U.S.A.).

#### L-[^14^C]lactate uptake and competition assays in *X. laevis* oocytes

Radiolabelled uptake and competition assays were performed at 27.5 °C on days 4–5 post-injection on five different days with oocytes from different frogs. Oocytes were washed four times at room temperature with 4 ml ND96 buffer (96 mM NaCl, 2 mM KCl, 1 mM MgCl₂, 1.8 mM CaCl₂, 10 mM MES, and 10 mM Tris-base; pH 6.4 adjusted with NaOH). After the final wash, the buffer was removed and replaced with ND96 buffer warmed to 27.5 °C containing 1 mM sodium L-lactate and 0.66 µM L-[¹⁴C]lactate (sodium salt, 150.6 mCi mmol⁻¹). Each assay tube contained 8–10 oocytes per timepoint.

For competition assays, the assay buffer was supplemented with 10 mM of the indicated test compounds (citrate, pyruvate, propionate, acetate, formate, nitrate, iodide, lactamide, propionamide, acetamide, or formamide) or 1 µM MMV007839, added simultaneously with L-lactate.

Oocytes were incubated in a 27.5 °C water bath for 15 min, then washed three times with ice-cold ND96 quenching buffer (96 mM NaCl, 2 mM KCl, 1 mM MgCl₂, 1.8 mM CaCl₂, 10 mM MES, 10 mM Tris-base, pH 6.4 adjusted with NaOH). To prevent L-[¹⁴C]lactate being effluxed by PfFNT at the end of the experiment, 2 µM of the known PfFNT-inhibitor MMV007839 was added to the ice-cold ND96 quenching buffer. A time-course was performed with PfFNT-expressing and non-injected oocytes where oocytes were either washed once in buffer without MMV007839 present or four times in buffer containing 2 µM MMV007839 to confirm that the addition of MMV007839 into the final washing buffer did not significantly affect uptake (**Fig. S13**).

As observed previously^45,51^, there was considerable variability between experiments in the magnitude of PfFNT-mediated L-[^14^C]lactate uptake relative to non-injected oocytes. Data were included only from experiments performed with oocyte preparations where L-[^14^C]lactate uptake by PfFNT-expressing oocytes, in the absence of any test compounds, exceeded that of non-injected oocytes by more than two-fold (**Table S6**). Individual oocyte data points >2 SD from the mean were treated as outliers and excluded.

#### Detection of lactate and lactamide using LC-MS

To investigate whether lactamide could be transported by PfFNT, oocytes were prepared for liquid chromatography–mass spectrometry (LC-MS) analysis. Oocytes were collected from three separate frogs on three different days, 4–5 days post-injection, and 40 oocytes per condition were used to increase the likelihood of detecting metabolites. Non-injected and PfFNT-expressing oocytes were washed three times in room-temperature ND96 before incubation for 4 hours at 27.5 °C in ND96 containing either 10 mM lactate or 10 mM lactamide. Experiments were quenched by removing external lactate/lactamide by washing three times in ice-cold ND96 buffer containing 2 µM MMV007839, and metabolites were isolated via a two-stage liquid-liquid phase extraction.^170,171^ Samples were dried under vacuum and reconstituted in 15:85 mobile phase A (10 mM ammonium formate + 0.15% formic acid) to mobile phase B (acetonitrile) prior to transfer into LC-MS vials.

Chromatographic separation was performed on an Ultimate 3000 RSLC nano UHPLC system (Dionex) using hydrophilic interaction liquid chromatography on a ZIC-cHILIC column (3 µm, 2.1 × 150 mm; Sequant, Merck).^170,171^ The flow rate was 0.2 mL/min over a 21-minute run with a binary gradient starting at 85:15 (A:B), changing to 36:64 at 10 min, then 64:36 at 12 min, and ending at 15:85 from 17–21 min. Mass detection was performed on a Q-Exactive Plus Orbitrap (Thermo Scientific) in both positive and negative electrospray modes over an m/z range of 60–200, with all other parameters set as previously described.^170,171^

A pooled mixture of all extracts served as a quality control (QC) to monitor signal reproducibility and analyte stability. Blank and QC samples were analyzed before and after the oocyte runs in both ionization modes, and QC samples were interspersed throughout the batch to ensure consistent measurements. Standards containing 50 µM of either lactate or lactamide were also run at the start of each sequence. Raw LC-MS data were imported into Skyline (version 25.1)^172,173^ for peak detection and quantification. Lactate and lactamide were identified based on retention time and exact mass by comparison with standards, and extracted ion chromatograms were generated for each compound (**Fig. S15; Table S7**). Peak areas were manually inspected and adjusted to ensure accurate integration. To evaluate uptake, peak areas from PfFNT-expressing oocytes were normalized to those from non-injected controls.

## Acknowledgements

This work was supported by resources and services provided by the National Computational Infrastructure (NCI), funded by the Australian Government. C.W. acknowledges support from an Australian Government Research Training Program (RTP) Ph.D. Scholarship and a Medical Advances Without Animals Honours Scholarship. K.P.G. acknowledges funding from an ANU Research School of Biology Seed Grant, as well as allocation of computing time and system administration support provided by the Faculty of Science, Agriculture, Business, and Law at the University of New England to the Linux cluster of the Karton group. *Xenopus* work was supported by funding obtained by G.v.D. from Australian Research Council (DP230100853). A.M.L. acknowledges funding from Australian National Health and Medical Research Council (GNT2028714). B.C. acknowledges funding from Australian Research Council (DP250100893).

We thank Josiah Bones and Elaine Tao for helping with setting up initial AMBER simulations, John Tanner for providing a script for generating Free Energy Surfaces, Dr Yiechang Lin for assistance with setting up initial flooding simulations, Ruitao Jin and Sitong He for running the GROMACS CpHMD simulations for H230 pK_a_ prediction, and Dr Deyun Qiu for guidance on performing minipreps and assistance with ordering compounds. We are also grateful to the *Xenopus laevis* Animal Husbandry Staff, the ANU Veterinary Services Team (Dr Jess McLeod and Dr Justin Clarke), as well as Angelika Bröer, Professor Stefan Bröer, Leqian Zhao, Dr Vicky Zhang, and Dr Courtney Winning for their assistance with *X. laevis* surgeries and oocyte preparation.

We further acknowledge the generous support of the ANU Joint Mass Spectrometry Facility Team (Dr Adam Carroll, Anitha Jeyasingham, and Joseph Boileau), and the ANU Biomolecular Resource Facility for sequencing.

## Contributions

All standard MD simulations, umbrella sampling simulations, AMBER CpHMD simulations, and associated simulation analyses: C.W.

Quantum chemical calculations and DFTB simulations: K.P.G.

*Xenopus laevis* oocyte surgeries: C.W. and S.J.F.

*Xenopus laevis* transport assays and analysis: C.W.

Wrote the original draft: C.W., K.P.G., A.M.L., B.C.

Contributed to writing and editing: C.W., K.P.G., G.v.D., A.M.L., B.C.

Contributed to figure-making: C.W., K.P.G., B.C., A.M.L., G.v.D.

Supervision: B.C., A.M.L., K.P.G., G.v.D., S.J.F.

Funding: B.C., A.M.L., G.v.D.

Ethics Approval: G.v.D.

Conceived the project: B.C, A.M.L, C.W.

## Supplementary Information

Supplementary Figures 1-16

Supplementary Tables 1-7

Supplementary Movies 1-4

## Data Availability

All computational and experimental data, input files, and analysis scripts are available upon request.

## References

1. Ashcroft, F., Gadsby, D. & Miller, C. The blurred boundary between channels and transporters: We dedicate this volume to the memory of Peter Läuger, a pioneer of the link between channels and pumps. Philosophical Transactions of the Royal Society B: Biological Sciences 364, 145 (2009).

2. Bihler, I. & Crane, R. K. Studies on the mechanism of intestinal absorption of sugars V. The influence of several cations and anions on the active transport of sugars, in vitro, by various preparations of hamster small intestine. BBA - Biochimica et Biophysica Acta 59, 78–93 (1962).

3. Hodgkin, A. L. & Huxley, A. F. A quantitative description of membrane current and its application to conduction and excitation in nerve. J Physiol 117, 500 (1952).

4. del Castillo, J. & Katz, B. Quantal components of the end-plate potential. J Physiol 124, 560 (1954).

5. Jardetzky, O. Simple Allosteric Model for Membrane Pumps. Nature 1966 211:5052 211, 969–970 (1966).

6. Jentsch, T. J. & Pusch, M. CLC chloride channels and transporters: Structure, function, physiology, and disease. Physiol Rev 98, 1493–1590 (2018).

7. Miller, C. ClC Chloride Channels Viewed Through a Transporter Lens. Nature 440, (2006).

8. Poroca, D. R., Pelis, R. M. & Chappe, V. M. ClC channels and transporters: Structure, physiological functions, and implications in human chloride channelopathies. Front Pharmacol 8, 1–25 (2017).

9. Yue, Z., Li, C. & Voth, G. A. The role of conformational change and key glutamic acid residues in the ClC-ec1 antiporter. Biophys J 122, 1068–1085 (2023).

10. Alper, S. L. & Sharma, A. K. The SLC26 gene family of anion transporters and channels. Mol Aspects Med 34, 494–515 (2013).

11. Chi, X. et al. Structural insights into the gating mechanism of human SLC26A9 mediated by its C-terminal sequence. Cell Discov 6, (2020).

12. Liu, F., Zhang, Z., Csanády, L., Gadsby, D. C. & Chen, J. Molecular Structure of the Human CFTR Ion Channel. Cell 169, 85–95.e8 (2017).

13. Khunweeraphong, N. & Kuchler, K. The human ABCG2 transporter engages three gates to control multidrug extrusion. iScience 28, 112125 (2025).

14. Windler, F. et al. The solute carrier SLC9C1 is a Na+/H+-exchanger gated by an S4-type voltage-sensor and cyclic-nucleotide binding. Nat Commun 9, 1–13 (2018).

15. Yeo, H., Mehta, V., Gulati, A. & Drew, D. Structure and electromechanical coupling of a voltage-gated Na+/H+ exchanger. Nature 623, 193–201 (2023).

16. Kalienkova, V., Peter, M. F., Rheinberger, J. & Paulino, C. Structures of a sperm-specific solute carrier gated by voltage and cAMP. Nature 623, 202–209 (2023).

17. Pal, K., Chowdhury, S. & Author, C. Architecture and rearrangements of a sperm-specific Na+/H+ exchanger. bioRxiv 2023.10.04.560940 (2023).

18. Fairman, W. A., Vandenberg, R. J., Arriza, J. L., Kavanaught, M. P. & Amara, S. G. An excitatory amino-acid transporter with properties of a ligand-gated chloride channel. Nature 375, 599–603 (1995).

19. Ryan, R. M. & Mindell, J. A. The uncoupled chloride conductance of a bacterial glutamate transporter homolog. Nat Struct Mol Biol 14, 365–371 (2007).

20. Lu, Y. et al. Structural basis for inositol pyrophosphate gating of the phosphate channel XPR1. Science (1979) 386, (2024).

21. Zhang, W. et al. Structural insights into the mechanism of phosphate recognition and transport by XPR1. Nature Communications 16, 1–10 (2025).

22. Wang, X. et al. KIDINS220 and InsP8 safeguard the stepwise regulation of phosphate exporter XPR1. Mol Cell 85, 3209–3224.e8 (2025).

23. Wang, Y. et al. Structure of the formate transporter FocA reveals a pentameric aquaporin-like channel. Nature 462, 467–472 (2009).

24. Suppmann, B. & Sawers, G. Isolation and characterization of hypophosphite-resistant mutants of Escherichia coli: identification of the FocA protein, encoded by the pfl operon, as a putative formate transporter. Mol Microbiol 11, 965–982 (1994).

25. Wiechert, M., Erler, H., Golldack, A. & Beitz, E. A widened substrate selectivity filter of eukaryotic formate-nitrite transporters enables high-level lactate conductance. FEBS Journal 284, 2663–2673 (2017).

26. Waight, A. B., Love, J. & Wang, D.-N. N. Structure and mechanism of a pentameric formate channel. Nat Struct Mol Biol 17, 31–38 (2010).

27. Lü, W. et al. The formate channel FocA exports the products of mixed-acid fermentation. Proc Natl Acad Sci U S A 109, 13254–13259 (2012).

28. Marchetti, R. V. et al. A lactate and formate transporter in the intraerythrocytic malaria parasite, Plasmodium falciparum. Nature Communications 2015 6:1 6, 1–7 (2015).

29. Erler, H., Ren, B., Gupta, N. & Beitz, E. The intracellular parasite Toxoplasma gondii harbors three druggable FNT-type formate and l-lactate transporters in the plasma membrane. Journal of Biological Chemistry 293, 17622–17630 (2018).

30. Zeng, J. M. et al. Identifying the major lactate transporter of Toxoplasma gondii tachyzoites. Sci Rep 11, 1–11 (2021).

31. Czyzewski, B. K. & Wang, D. N. Identification and characterization of a bacterial hydrosulphide ion channel. Nature 2012 483:7390 483, 494–497 (2012).

32. Lü, W. et al. Structural and functional characterization of the nitrite channel NirC from Salmonella typhimurium. Proc Natl Acad Sci U S A 109, 18395–18400 (2012).

33. Jia, W., Tovell, N., Clegg, S., Trimmer, M. & Cole, J. A single channel for nitrate uptake, nitrite export and nitrite uptake by Escherichia coli NarU and a role for NirC in nitrite export and uptake. Biochemical Journal 417, 297–304 (2009).

34. Waight, A. B., Love, J. & Wang, D.-N. N. Structure and mechanism of a pentameric formate channel. Nat Struct Mol Biol 17, 31–38 (2010).

35. Lü, W. et al. pH-dependent gating in a FocA formate channel. Science (1979) 332, 352–354 (2011).

36. Lü, W. et al. Structural and functional characterization of the nitrite channel NirC from Salmonella typhimurium. Proc Natl Acad Sci U S A 109, 18395–18400 (2012).

37. Czyzewski, B. K. & Wang, D. N. Identification and characterization of a bacterial hydrosulphide ion channel. Nature 2012 483:7390 483, 494–497 (2012).

38. Wang, Y. et al. Structure of the formate transporter FocA reveals a pentameric aquaporin-like channel. Nature 462, 467–472 (2009).

39. Lü, W. et al. pH-dependent gating in a FocA formate channel. Science (1979) 332, 352–354 (2011).

40. Padhi, S., Reddy, L. K. & Priyakumar, U. D. pH-mediated gating and formate transport mechanism in the Escherichia coli formate channel. 10.1080/08927022.2017.1353691 43, 1300–1306 (2017).

41. Feng, Z., Hou, T. & Li, Y. Concerted movement in pH-dependent gating of FocA from molecular dynamics simulations. J Chem Inf Model 52, 2119–2131 (2012).

42. Lv, X., Liu, H., Ke, M. & Gong, H. Exploring the pH-dependent substrate transport mechanism of FocA using molecular dynamics simulation. Biophys J 105, 2714–2723 (2013).

43. Mukherjee, M., Gupta, A. & Sankararamakrishnan, R. Is the E. coli Homolog of the Formate/Nitrite Transporter Family an Anion Channel? A Computational Study. Biophys J 118, 846–860 (2020).

44. Atkovska, K. & Hub, J. S. Energetics and mechanism of anion permeation across formate-nitrite transporters. Sci Rep 7, 1–14 (2017).

45. Marchetti, R. V. et al. A lactate and formate transporter in the intraerythrocytic malaria parasite, Plasmodium falciparum. Nature Communications 2015 6:1 6, 1–7 (2015).

46. Wu, B. et al. Identity of a Plasmodium lactate/H+ symporter structurally unrelated to human transporters. Nature Communications 2015 6:1 6, 1–8 (2015).

47. Cranmer, S. L., Conant, A. R., Gutteridge, W. E. & Halestrap, A. P. Characterization of the enhanced transport of L- and D-lactate into human red blood cells infected with Plasmodium falciparum suggests the presence of a novel saturable lactate proton cotransporter. Journal of Biological Chemistry 270, 15045–15052 (1995).

48. Elliott, J. L., Saliba, K. J. & Kirk, K. Transport of lactate and pyruvate in the intraerythrocytic malaria parasite, Plasmodium falciparum. Biochemical Journal 355, 733 (2001).

49. Kanaani, J. & Ginsburg, H. Transport of lactate in Plasmodium falciparum-infected human erythrocytes. J Cell Physiol 149, 469–476 (1991).

50. Golldack, A. et al. Substrate-analogous inhibitors exert antimalarial action by targeting the Plasmodium lactate transporter PfFNT at nanomolar scale. PLoS Pathog 13, 1–18 (2017).

51. Hapuarachchi, S. V. et al. The Malaria Parasite’s Lactate Transporter PfFNT Is the Target of Antiplasmodial Compounds Identified in Whole Cell Phenotypic Screens. PLoS Pathog 13, 1–24 (2017).

52. Miller, L. H., Ackerman, H. C., Su, X. Z. & Wellems, T. E. Malaria biology and disease pathogenesis: Insights for new treatments. Nat Med 19, 156–167 (2013).

53. MacRae, J. I. et al. Mitochondrial metabolism of sexual and asexual blood stages of the malaria parasite Plasmodium falciparum. BMC Biol 11, 1–10 (2013).

54. Mehta, M., Sonawat, H. M. & Sharma, S. Malaria parasite-infected erythrocytes inhibit glucose utilization in uninfected red cells. FEBS Lett 579, 6151–6158 (2005).

55. Wu, B. et al. Identity of a Plasmodium lactate/H+ symporter structurally unrelated to human transporters. Nature Communications 2015 6:1 6, 1–8 (2015).

56. Golldack, A. et al. Substrate-analogous inhibitors exert antimalarial action by targeting the Plasmodium lactate transporter PfFNT at nanomolar scale. PLoS Pathog 13, 1–18 (2017).

57. Hapuarachchi, S. V. et al. The Malaria Parasite’s Lactate Transporter PfFNT Is the Target of Antiplasmodial Compounds Identified in Whole Cell Phenotypic Screens. PLoS Pathog 13, 1–24 (2017).

58. Walloch, P., Hansen, C., Priegann, T., Schade, D. & Beitz, E. Pentafluoro-3-hydroxy-pent-2-en-1-ones Potently Inhibit FNT-Type Lactate Transporters from all Five Human-Pathogenic Plasmodium Species. ChemMedChem 16, 1283–1289 (2021).

59. Davies, H. et al. The Plasmodium Lactate/H+ Transporter PfFNT Is Essential and Druggable In Vivo. Antimicrob Agents Chemother 10.1128/AAC.00356-23 (2023) doi:10.1128/AAC.00356-23.

60. Wiechert, M. & Beitz, E. Mechanism of formate–nitrite transporters by dielectric shift of substrate acidity. EMBO J 36, 949–958 (2017).

61. Wang, N. et al. Structural basis of human monocarboxylate transporter 1 inhibition by anti-cancer drug candidates. Cell 184, 370–383.e13 (2021).

62. Xu, B. et al. Embigin facilitates monocarboxylate transporter 1 localization to the plasma membrane and transition to a decoupling state. Cell Rep 40, (2022).

63. Peng, X. et al. Structural characterization of the Plasmodium falciparum lactate transporter PfFNT alone and in complex with antimalarial compound MMV007839 reveals its inhibition mechanism. PLoS Biol 19, 1–19 (2021).

64. Lyu, M., Su, C., Kazura, J. W. & Yu, E. W. Structural basis of transport and inhibition of the Plasmodium falciparum transporter PfFNT. EMBO Rep 22, 1–12 (2021).

65. Helmstetter, F., Arnold, P., Höger, B., Petersen, L. M. & Beitz, E. Formate–nitrite transporters carrying nonprotonatable amide amino acids instead of a central histidine maintain pH-dependent transport. J Biol Chem 294, 623 (2019).

66. Schmidt, J. D. R. & Beitz, E. Mutational widening of constrictions in a formate–nitrite/H+ transporter enables aquaporin-like water permeability and proton conductance. Journal of Biological Chemistry 298, 101513 (2022).

67. Wiechert, M. & Beitz, E. Formate-nitrite transporters: Monoacids ride the dielectric slide. Channels 11, 365–367 (2017).

68. Olsson, M. H. M., SØndergaard, C. R., Rostkowski, M. & Jensen, J. H. PROPKA3: Consistent treatment of internal and surface residues in empirical p K a predictions. J Chem Theory Comput 7, 525–537 (2011).

69. Huang, Y., Henderson, J. A. & Shen, J. Continuous Constant pH Molecular Dynamics Simulations of Transmembrane Proteins. in Methods in Molecular Biology vol. 2302 275–287 (Humana Press Inc., 2021).

70. Martins de Oliveira, V., Liu, R. & Shen, J. Constant pH molecular dynamics simulations: Current status and recent applications. Current Opinion in Structural Biology vol. 77 Preprint at 10.1016/j.sbi.2022.102498 (2022).

71. Harris, R. C. & Shen, J. GPU-Accelerated Implementation of Continuous Constant pH Molecular Dynamics in Amber: pKa Predictions with Single-pH Simulations. J Chem Inf Model 59, 4821–4832 (2019).

72. Huang, Y., Harris, R. C. & Shen, J. Generalized Born Based Continuous Constant pH Molecular Dynamics in Amber: Implementation, Benchmarking and Analysis. J Chem Inf Model 58, 1372–1383 (2018).

73. Harris, J. A. et al. GPU-Accelerated All-Atom Particle-Mesh Ewald Continuous Constant pH Molecular Dynamics in Amber. J Chem Theory Comput 18, 7510–7527 (2022).

74. Woodrow, C. J., Penny, J. I. & Krishna, S. Intraerythrocytic Plasmodium falciparum expresses a high affinity facilitative hexose transporter. Journal of Biological Chemistry 274, 7272–7277 (1999).

75. Jiang, X. et al. Structural Basis for Blocking Sugar Uptake into the Malaria Parasite Plasmodium falciparum. Cell 183, 258–268.e12 (2020).

76. Penna-Coutinho, J., Cortopassi, W. A., Oliveira, A. A., França, T. C. C. & Krettli, A. U. Antimalarial Activity of Potential Inhibitors of Plasmodium falciparum Lactate Dehydrogenase Enzyme Selected by Docking Studies. PLoS One 6, e21237 (2011).

77. Krugliak, M. & Ginsburg, H. The evolution of the new permeability pathways in Plasmodium falciparum—infected erythrocytes—a kinetic analysis. Exp Parasitol 114, 253–258 (2006).

78. Wiechert, M. & Beitz, E. Mechanism of formate–nitrite transporters by dielectric shift of substrate acidity. EMBO J 36, 949–958 (2017).

79. Elliott, J. L., Saliba, K. J. & Kirk, K. Transport of lactate and pyruvate in the intraerythrocytic malaria parasite, Plasmodium falciparum. Biochemical Journal 355, 733 (2001).

80. Sawers, R. G. Formate and its role in hydrogen production in Escherichia coli. Biochem Soc Trans 33, 42–46 (2005).

81. Lü, W. et al. The formate channel FocA exports the products of mixed-acid fermentation. Proc Natl Acad Sci U S A 109, 13254–13259 (2012).

82. Kammel, M. & Sawers, R. G. The FocA channel functions to maintain intracellular formate homeostasis during Escherichia coli fermentation. 1–6 (2022) doi:10.1099/mic.0.001168.

83. Suppmann, B. & Sawers, G. Isolation and characterization of hypophosphite-resistant mutants of Escherichia coli: identification of the FocA protein, encoded by the pfl operon, as a putative formate transporter. Mol Microbiol 11, 965–982 (1994).

84. Erler, H., Ren, B., Gupta, N. & Beitz, E. The intracellular parasite Toxoplasma gondii harbors three druggable FNT-type formate and l-lactate transporters in the plasma membrane. Journal of Biological Chemistry 293, 17622–17630 (2018).

85. Lü, W. et al. The formate/nitrite transporter family of anion channels. Biol Chem 394, 715–727 (2013).

86. Schnell, J. R. & Chou, J. J. Structure and Mechanism of the M2 Proton Channel of Influenza A Virus. Nature 451, 591 (2008).

87. Zheng, W. et al. pH regulates potassium conductance and drives a constitutive proton current in human TMEM175. Sci Adv 8, 1568 (2022).

88. Ramsey, I. S., Moran, M. M., Chong, J. A. & Clapham, D. E. A voltage-gated proton-selective channel lacking the pore domain. Nature 2006 440:7088 440, 1213–1216 (2006).

89. Chamberlin, A. et al. Hydrophobic plug functions as a gate in voltage-gated proton channels. Proc Natl Acad Sci U S A 111, E273–E282 (2014).

90. Saotome, K. et al. Structures of the otopetrin proton channels Otop1 and Otop3. Nature Structural & Molecular Biology 2019 26:6 26, 518–525 (2019).

91. Scerri, E. The Born-Oppenheimer Approximation and its role in the reduction of chemistry. Foundations of Chemistry 2025 27:2 27, 183–197 (2025).

92. Zheng, R., Jing, Y., Chen, L. & Shi, Q. Theory of proton coupled electron transfer reactions: Assessing the Born–Oppenheimer approximation for the proton motion using an analytically solvable model. Chem Phys 379, 39–45 (2011).

93. Li, C., Requist, R. & Gross, E. K. U. Density functional theory of electron transfer beyond the Born-Oppenheimer approximation: Case study of LiF. J Chem Phys 148, (2018).

94. Marx, D. Proton Transfer 200 Years after von Grotthuss: Insights from Ab Initio Simulations. ChemPhysChem 7, 1848–1870 (2006).

95. Nanni, L. Modelling proton tunneling in hydrogen bonds through path integral method. Chem Phys 574, 112054 (2023).

96. Wang, N. et al. Structural basis of human monocarboxylate transporter 1 inhibition by anti-cancer drug candidates. Cell 184, 370–383.e13 (2021).

97. Xu, B. et al. Embigin facilitates monocarboxylate transporter 1 localization to the plasma membrane and transition to a decoupling state. Cell Rep 40, (2022).

98. Köpnick, A. L., Geistlinger, K. & Beitz, E. Cysteine 159 delineates a hinge region of the alternating access monocarboxylate transporter 1 and is targeted by cysteine-modifying inhibitors. FEBS J 288, 6052–6062 (2021).

99. Wilson, M. C., Meredith, D., Bunnun, C., Sessions, R. B. & Halestrap, A. P. Studies on the DIDS-binding site of monocarboxylate transporter 1 suggest a homology model of the open conformation and a plausible translocation cycle. Journal of Biological Chemistry 284, 20011–20021 (2009).

100. Rossmann, R., Sawers, G. & Böck, A. Mechanism of regulation of the formate-hydrogenlyase pathway by oxygen, nitrate, and pH: definition of the formate regulon. Mol Microbiol 5, 2807–2814 (1991).

101. Davies, H. et al. The Plasmodium Lactate/H+ Transporter PfFNT Is Essential and Druggable In Vivo. Antimicrob Agents Chemother 10.1128/AAC.00356-23 (2023) doi:10.1128/AAC.00356-23.

102. Jentsch, T. J. & Pusch, M. CLC chloride channels and transporters: Structure, function, physiology, and disease. Physiol Rev 98, 1493–1590 (2018).

103. Miller, C. ClC Chloride Channels Viewed Through a Transporter Lens. Nature 440, (2006).

104. Poroca, D. R., Pelis, R. M. & Chappe, V. M. ClC channels and transporters: Structure, physiological functions, and implications in human chloride channelopathies. Front Pharmacol 8, 1–25 (2017).

105. Yue, Z., Li, C. & Voth, G. A. The role of conformational change and key glutamic acid residues in the ClC-ec1 antiporter. Biophys J 122, 1068–1085 (2023).

106. Alper, S. L. & Sharma, A. K. The SLC26 gene family of anion transporters and channels. Mol Aspects Med 34, 494–515 (2013).

107. Chi, X. et al. Structural insights into the gating mechanism of human SLC26A9 mediated by its C-terminal sequence. Cell Discov 6, (2020).

108. Liu, F., Zhang, Z., Csanády, L., Gadsby, D. C. & Chen, J. Molecular Structure of the Human CFTR Ion Channel. Cell 169, 85–95.e8 (2017).

109. Khunweeraphong, N. & Kuchler, K. The human ABCG2 transporter engages three gates to control multidrug extrusion. iScience 28, 112125 (2025).

110. Fairman, W. A., Vandenberg, R. J., Arriza, J. L., Kavanaught, M. P. & Amara, S. G. An excitatory amino-acid transporter with properties of a ligand-gated chloride channel. Nature 375, 599–603 (1995).

111. Ryan, R. M. & Mindell, J. A. The uncoupled chloride conductance of a bacterial glutamate transporter homolog. Nat Struct Mol Biol 14, 365–371 (2007).

112. Lu, Y. et al. Structural basis for inositol pyrophosphate gating of the phosphate channel XPR1. Science (1979) 386, (2024).

113. Zhang, W. et al. Structural insights into the mechanism of phosphate recognition and transport by XPR1. Nature Communications 16, 1–10 (2025).

114. Wang, X. et al. KIDINS220 and InsP8 safeguard the stepwise regulation of phosphate exporter XPR1. Mol Cell 85, 3209–3224.e8 (2025).

115. Schrodinger. Maestro, Schrödinger Release 2022-1: Preprint at (2021).

116. Lomize, M. A., Pogozheva, I. D., Joo, H., Mosberg, H. I. & Lomize, A. L. OPM database and PPM web server: resources for positioning of proteins in membranes. Nucleic Acids Res 40, (2012).

117. Jo, S., Kim, T., Iyer, V. G. & Im, W. CHARMM-GUI: A web-based graphical user interface for CHARMM. J Comput Chem 29, 1859–1865 (2008).

118. Lee, J. et al. CHARMM-GUI Input Generator for NAMD, GROMACS, AMBER, OpenMM, and CHARMM/OpenMM Simulations Using the CHARMM36 Additive Force Field. J Chem Theory Comput 12, 405–413 (2016).

119. Kim, J. et al. Structure and Drug Resistance of the Plasmodium falciparum Transporter PfCRT. Nature 576, 315 (2019).

120. Sardar, R. et al. In-silico profiling and structural insights into the impact of nSNPs in the P. falciparum acetyl-CoA transporter gene to understand the mechanism of drug resistance in malaria. J Biomol Struct Dyn 39, 558–569 (2021).

121. Feng, Z., Hou, T. & Li, Y. Concerted movement in pH-dependent gating of FocA from molecular dynamics simulations. J Chem Inf Model 52, 2119–2131 (2012).

122. Lv, X., Liu, H., Ke, M. & Gong, H. Exploring the pH-dependent substrate transport mechanism of FocA using molecular dynamics simulation. Biophys J 105, 2714–2723 (2013).

123. Atkovska, K. & Hub, J. S. Energetics and mechanism of anion permeation across formate-nitrite transporters. Sci Rep 7, 1–14 (2017).

124. Mukherjee, M., Gupta, A. & Sankararamakrishnan, R. Is the E. coli Homolog of the Formate/Nitrite Transporter Family an Anion Channel? A Computational Study. Biophys J 118, 846–860 (2020).

125. Winterberg, M. & Kirk, K. A high-sensitivity HPLC assay for measuring intracellular Na+ and K+ and its application to Plasmodium falciparum infected erythrocytes. Scientific Reports 2016 6:1 6, 1–6 (2016).

126. Mauritz, J. M. A. et al. X-Ray Microanalysis Investigation of the Changes in Na, K, and Hemoglobin Concentration in Plasmodium falciparum-Infected Red Blood Cells. Biophys J 100, 1438 (2011).

127. Henry, R. I. et al. An acid-loading chloride transport pathway in the intraerythrocytic malaria parasite, Plasmodium falciparum. Journal of Biological Chemistry 285, 18615–18626 (2010).

128. Humphrey, W., Dalke, A. & Schulten, K. VMD: visual molecular dynamics. J Mol Graph 14, 33–38 (1996).

129. O’Boyle, N. M. et al. Open Babel: An Open chemical toolbox. J Cheminform 3, 1–14 (2011).

130. O’Boyle, N. M. et al. Open Babel. J Cheminform 3, 1–14 (2011).

131. Wang, J., Wolf, R. M., Caldwell, J. W., Kollman, P. A. & Case, D. A. Development and testing of a general amber force field. J Comput Chem 25, 1157–1174 (2004).

132. Baaden, M., Bemy, F. & Wipff, G. The chloroform / TBP / Aqueous Nitric Acid Interfacial System: a Molecular Dynamics Investigation. J Mol Liq 90, 9 (2001).

133. Case, D. A., et al. Amber20. Preprint at (2021).

134. Bouysset, C. & Fiorucci, S. ProLIF: a library to encode molecular interactions as fingerprints. J Cheminform 13, (2021).

135. Tanner, J. D., Richards, S. N. & Corry, B. Molecular basis of the functional conflict between chloroquine and peptide transport in the Malaria parasite chloroquine resistance transporter PfCRT. Nature Communications 16, 1–16 (2025).

136. Roe, D. R. & Cheatham, T. E. PTRAJ and CPPTRAJ: Software for Processing and Analysis of Molecular Dynamics Trajectory Data. J Chem Theory Comput 10.1021/ct400341p (2013) doi:10.1021/ct400341p.

137. E, W., Ren, W. & Vanden-Eijnden, E. Simplified and improved string method for computing the minimum energy paths in barrier-crossing events. Journal of Chemical Physics 126, (2007).

138. Wiechert, M., Erler, H. & Beitz, E. A widened substrate selectivity filter of eukaryotic formate-nitrite transporters enables high-level lactate conductance. 10.1111/febs.14117 doi:10.1111/febs.14117.

139. Helmstetter, F., Arnold, P., Höger, B., Petersen, L. M. & Beitz, E. Formate–nitrite transporters carrying nonprotonatable amide amino acids instead of a central histidine maintain pH-dependent transport. J Biol Chem 294, 623 (2019).

140. Adamson, L. S. R. et al. Pore structure controls stability and molecular flux in engineered protein cages. Sci Adv 8, (2022).

141. Schrodinger LLC. The PyMOL Molecular Graphics System, Version 1.8. Preprint at (2015).

142. Landrum, G. RDKit: Open-Source Cheminformatics Software. Preprint at (2016).

143. Peverati, R. & Truhlar, D. G. Communication: A global hybrid generalized gradient approximation to the exchange-correlation functional that satisfies the second-order density-gradient constraint and has broad applicability in chemistry. J Chem Phys 135, 191102 (2011).

144. Grimme, S., Antony, J., Ehrlich, S. & Krieg, H. A consistent and accurate ab initio parametrization of density functional dispersion correction (DFT-D) for the 94 elements H-Pu. Journal of Chemical Physics 132, (2010).

145. Becke, A. D. & Johnson, E. R. A density-functional model of the dispersion interaction. Journal of Chemical Physics 123, (2005).

146. Weigend, F. & Ahlrichs, R. Balanced basis sets of split valence, triple zeta valence and quadruple zeta valence quality for H to Rn: Design and assessment of accuracy. Physical Chemistry Chemical Physics 7, 3297–3305 (2005).

147. Frisch, M. J. et al. Gaussian 16, Revision C.01. Preprint at (2016).

148. Goerigk, L. et al. A look at the density functional theory zoo with the advanced GMTKN55 database for general main group thermochemistry, kinetics and noncovalent interactions. Physical Chemistry Chemical Physics 19, 32184–32215 (2017).

149. Peng, C. & Bernhard Schlegel, H. Combining Synchronous Transit and Quasi-Newton Methods to Find Transition States. Isr J Chem 33, 449–454 (1993).

150. Fukui, K. The path of chemical reactions - the IRC approach. Acc Chem Res 14, 363–368 (2002).

151. Fukui, K. Formulation of the reaction coordinate. Journal of Physical Chemistry 74, 4161 (2002).

152. Gaus, M., Cui, Q. & Elstner, M. DFTB3: Extension of the Self-Consistent-Charge Density-Functional Tight-Binding Method (SCC-DFTB). J Chem Theory Comput 7, 931–948 (2011).

153. Kubillus, M., Kubař, T., Gaus, M., Řezáč, J. & Elstner, M. Parameterization of the DFTB3 Method for Br, Ca, Cl, F, I, K, and Na in Organic and Biological Systems. J Chem Theory Comput 11, 332–342 (2014).

154. Hourahine, B. et al. DFTB+, a software package for efficient approximate density functional theory based atomistic simulations. Journal of Chemical Physics 152, 124101 (2020).

155. Liu, R., Yue, Z., Tsai, C. C. & Shen, J. Assessing Lysine and Cysteine Reactivities for Designing Targeted Covalent Kinase Inhibitors. J Am Chem Soc 141, 6553–6560 (2019).

156. Henderson, J. A., Verma, N., Harris, R. C., Liu, R. & Shen, J. Assessment of proton-coupled conformational dynamics of SARS and MERS coronavirus papain-like proteases: Implication for designing broad-spectrum antiviral inhibitors. Journal of Chemical Physics 153, 1–10 (2020).

157. Peeples, C. A., Liu, R. & Shen, J. Force Field Limitations of All-Atom Continuous Constant pH Molecular Dynamics. bioRxiv 2024.09.03.611076 (2024) doi:10.1101/2024.09.03.611076.

158. Henderson, J. A. et al. A Guide to the Continuous Constant pH Molecular Dynamics Methods in Amber and CHARMM [Article v1.0]. Living J Comput Mol Sci 4, 1563–1563 (2022).

159. Suh, D., Zhang, H. & Im, W. CHARMM-GUI constant-pH simulator for the constant-pH molecular dynamics simulations. Biophys J 123, 423a (2024).

160. Aho, N. et al. Scalable Constant pH Molecular Dynamics in GROMACS. J Chem Theory Comput 18, 6148–6160 (2022).

161. Huang, J. et al. CHARMM36m: An Improved Force Field for Folded and Intrinsically Disordered Proteins. Nat Methods 14, 71 (2017).

162. Michaud-Agrawal, N., Denning, E. J., Woolf, T. B. & Beckstein, O. MDAnalysis: A toolkit for the analysis of molecular dynamics simulations. J Comput Chem 32, 2319–2327 (2011).

163. Harris, C. R. et al. Array programming with NumPy. Nature 2020 585:7825 585, 357–362 (2020).

164. Mckinney, W. Data Structures for Statistical Computing in Python. (2010).

165. Hunter, J. Matplotlib: A 2D Graphics Environment. Comput Sci Eng 9, (2007).

166. Virtanen, P. et al. SciPy 1.0: fundamental algorithms for scientific computing in Python. Nature Methods 2020 17:3 17, 261–272 (2020).

167. Smart, O. S., Neduvelil, J. G., Wang, X., Wallace, B. A. & Sansomt, M. S. P. HOLE: A Program for the Analysis of the Pore Dimensions of Ion Channel Structural Models. http://www.golden.com/golden/for (1996).

168. Gracia, L. Luisico/clustering: VMD plugin to calculate and visualize clusters of conformations for a trajectory. https://github.com/luisico/clustering.

169. Zeng, J. M. et al. Identifying the major lactate transporter of Toxoplasma gondii tachyzoites. Sci Rep 11, 1–11 (2021).

170. Fairweather, S. J. et al. A GC-MS/Single-Cell Method to Evaluate Membrane Transporter Substrate Specificity and Signaling. Front Mol Biosci 8, 646574 (2021).

171. Fairweather, S. J. et al. Coordinated action of multiple transporters in the acquisition of essential cationic amino acids by the intracellular parasite Toxoplasma gondii. PLoS Pathog 17, e1009835 (2021).

172. Adams, K. J. et al. Skyline for Small Molecules: A Unifying Software Package for Quantitative Metabolomics. J Proteome Res 19, 1447–1458 (2020).

173. MacLean, B., et al. Skyline: an open source document editor for creating and analyzing targeted proteomics experiments. Bioinformatics 26, 966–968 (2010).

