## Supplementary Information for "A proton transfer mechanism in the malaria parasite lactate/H^+^ symporter reveals a channel-like transporter without conformational changes"

### Supplementary Movies

**Movie S1. H230 can protonate via a Grotthuss mechanism.** A representative trajectory from DFTB simulations of a 5 Å region within the transport cavity, showing a proton 'hopping' into the cavity to protonate H230 at the δ nitrogen. The region simulated in DFTB simulations was aligned and superimposed back into the transport cavity of a PfFNT subunit (shown in a light grey cartoon representation).

**Movie S2. Binding events in the $His^{+}/Lac^{-}$ system. a)** A representative trajectory from $His^{+}/Lac^{-}$ flooding simulations where two $Lac^{-}$ molecules were observed to bind within the transport cavities of different subunits of PfFNT. A cutaway of the PfFNT pentamer is shown in a light grey cartoon representation. Pink dashed circles are added to highlight binding events. **b)** A close-up view of a representative binding event from the $His^{+}/Lac^{-}$ flooding simulations. $Lac^{-}$ can be observed to pass the intracellular constriction and bind to $His^{+}$ once inside.

**Movie S3. $His^{+}$ favourably transfers a proton to $Lac^{-}$, and the newly formed $Lac^{0}$ dissociates. a)** A representative trajectory from DFTB simulations of a 5 Å region within the transport cavity with bound $Lac^{-}$. The movie shows H230 transferring a proton to $Lac^{-}$ from the δ NH on the H230 sidechain. The region simulated in DFTB simulations was aligned and superimposed back into the transport cavity of a PfFNT subunit (shown in a light grey cartoon representation). **b)** A representative trajectory from MD simulations following manual proton transfer, where $Lac^{0}$ dissociates from the transport cavity. H230 and $Lac^{-}/Lac^{0}$ are shown in a ball and stick representation and are coloured by atom type. Other constriction residues are shown in a white ball and stick representation.

**Movie S4. PfFNT can function as a formic acid channel when H230 is charged.** A representative trajectory from flooding simulations of the $His^{+}/For^{0}$ system, showing neutral formic acid molecules passively diffusing through the PfFNT transport cavity.

#### Supplementary Figures and Tables

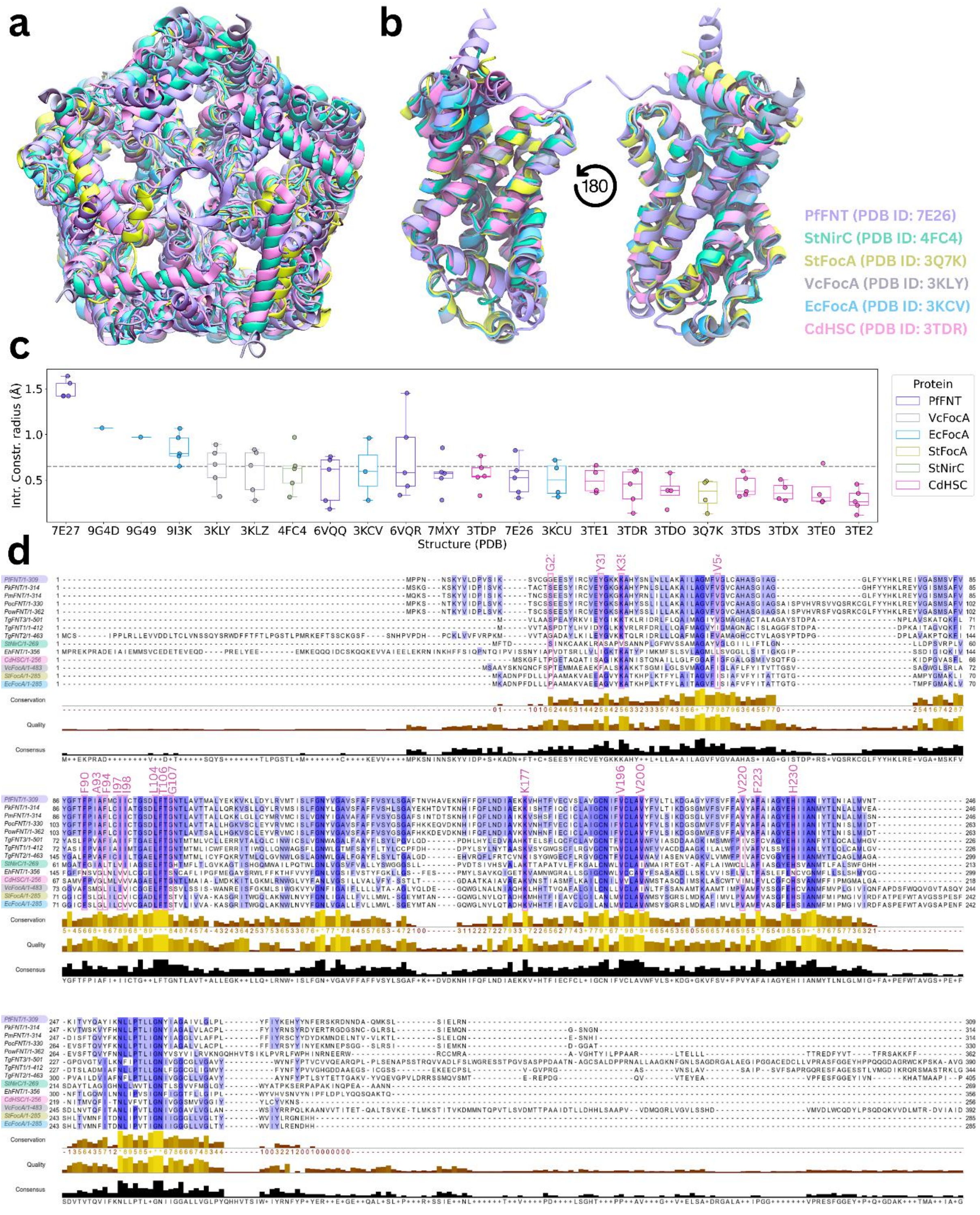


**Figure S1. FNTs share similarity in structure and sequence.** Structural alignments of **a)** the whole pentamer and **b)** a single subunit of PfFNT (PDB ID: 7E26), StNirC (PDB ID: 4FC4), StFocA (PDB ID: 3Q7K), VcFocA (PDB ID: 3KLY), EcFocA (PDB ID: 3KCV), and CdHSC (PDB ID: 3TDR). Proteins were aligned using the MultiSeq Stamp Structural Alignment plugin within VMD[1]. The average percent identity across all proteins was 30.39%. Comparisons for percentage identity of PfFNT relative to other proteins are given in Table S1. **c)** Boxplots of the average intracellular constriction radius from each subunit of the 22 available FNT structures. Transport pathway radius was measured using HOLE. Structures are coloured by protein. Average radius is shown as a grey line. **d)** Sequence alignments for VcFocA (F0M16_11295), StFocA (G1X15_10175), EcFocA (ECs0987), CdHSC (SAMEA1710456_03117), StNirC (STM3476), PfFNT (PF3D7_0316600), as well as the *P. knowlesi* FNT (PKNH_0825200), *P.* malariae FNT (PmFNT, PMLGA01_080027300), *P. ovale curtisi* FNT (PocFNT, POVCU1_016520), *P. ovale wallikeri* FNT (PowFNT, POVWA1_022090), and the three *Toxoplasma gondii* FNTs, TgFNT1 (TGGT1_209800), TgFNT2 (TGGT1_292110) and TgFNT3 (TGGT1_229170). The sequence of another FNT, EhFNT from *Entamoeba histolytica* (EHI5A_201460), lacking the central histidine was also included in the alignment. Sequences are coloured by percent identity from lowest (white) to highest (purple). Alignment was made using Clustal Omega[2,3] and visualised using JalView[4].

**Table S1. RMSD, sequence, and structural identity comparisons between PfFNT and closely related FNTs.** RMSD and percent structural identity were calculated using the MultiSeq Stamp Structural Alignment tool in VMD. Percent sequence identity was calculated using the pairwise alignment tool in JalView.

| Protein Comparison | RMSD | Percent Structural Identity | Percent Sequence Identity |
|---|---|---|---|
| PfFNT (7E26) – StNirC (4FC4) | 1.9361 | 20.34 | 23.69 |
| PfFNT (7E26) – StFocA (3Q7K) | 2.2109 | 20.85 | 27.18 |
| PfFNT (7E26) – VcFocA (3KLY) | 1.8268 | 18.47 | 21.97 |
| PfFNT (7E26) – EcFocA (3KCV) | 2.2965 | 20.71 | 27.27 |
| PfFNT (7E26) – CdHSC (3TDR) | 1.6837 | 22.45 | 29.78 |
| PfFNT – PkFNT | - | - | 72.82 |
| PfFNT – PmFNT | - | - | 72.49 |
| PfFNT – PocFNT | - | - | 70.55 |
| PfFNT – PowFNT | - | - | 64.09 |
| PfFNT – PvFNT | - | - | 73.05 |
| PfFNT – TgFNT1 | - | - | 32.54 |
| PfFNT – TgFNT2 | - | - | 33.76 |
| PfFNT – TgFNT3 | - | - | 36.00 |

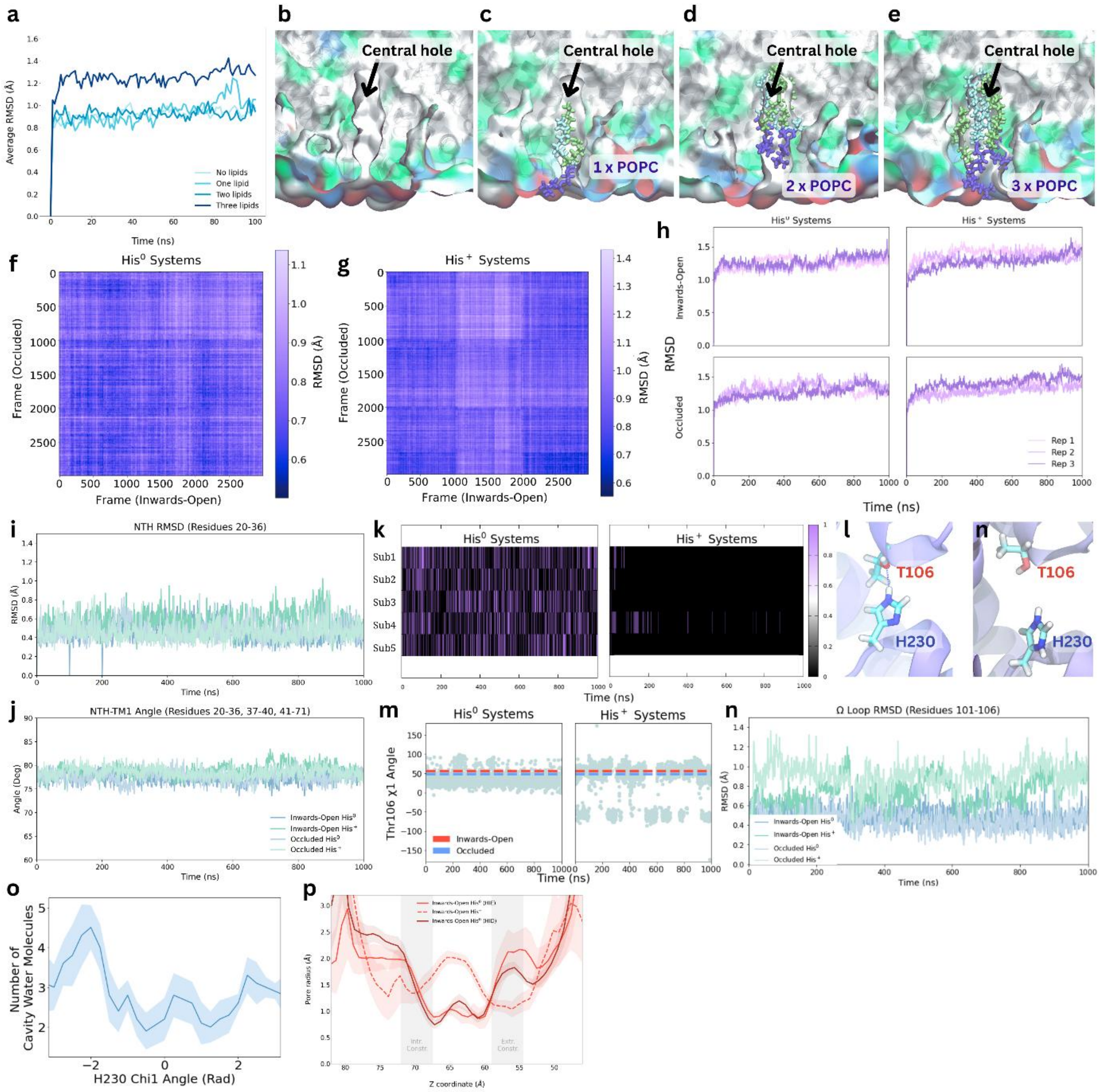


**Figure S2. Dynamics of the PfFNT central hole and transport cavities. a)** Comparison of the RMSD of the central hole lining residues over time following insertion of one, two, or three lipids into the central hole. The RMSD of the central hole lining residues without inserted lipids is a reference. **(b-e)** Visualisations of the central hole when no lipids, one lipid, two lipids, and three lipids have been inserted. When inspecting the trajectories of the simulations with three lipids, we saw that one lipid appeared to progressively get 'pushed' out of the central hole by the other two lipids. When only one lipid was in the central hole, the lipid could not fully occupy the central hole: leaving a partial vacuum of space within the protein. We observed that two lipids could stabilise each other to occupy the central hole best. Thus, we include two lipids within the central hole of PfFNT in all subsequent simulations. **(f-g)** Frame-by-frame comparisons of RMSD between 'inwards-open' and 'occluded' systems where H230 was neutral or charged. **h)** RMSD of the whole protein backbone compared between replicates for each simulation system without ligands present. Measurements of **i)** RMSD of residues forming the N-terminal helix (NTH) and **j)** the angle formed between the N-terminal helix and transmembrane helix 1 (TM1) over time in the inwards-open His$^0$, inwards-open His$^+$, occluded His$^0$, and occluded His$^+$ systems. The N-terminal helix appears stable in all systems and does not substantially fluctuate in position. **k)** Heatmaps of the His$^0$ and His$^+$ systems showing the presence or absence of a hydrogen bond (HB) between H230 and T106 in each subunit of a representative trajectory. **l)** Hydrogen bond between H230 and T106 in the His$^0$ system. Electrostatic repulsion between the OH group of the T106 sidechain and the NH group of the H230 sidechain in the $His^+$ system disrupts the hydrogen bond normally present between H230 and T106. **m)** Measured T106 χ1 dihedral angles over time in the His0 and His+ systems. Dashed lines are added to show the dihedral angles adopted by T106 in the inwards-open (red) and occluded (blue) cryo-EM structures. **n)** RMSD of the entire Ω loop (formed by residues 101 to 106) of each subunit for the Inwards-Open $His^0$, Inwards-Open $His^+$, Occluded $His^0$, and Occluded $His^+$ systems over time. **o)** Average number of water molecules (± SEM) within 5 Å of the His0 sidechain from 1D US simulations where H230 was restrained at different χ1 angles. **p)** Average transport pathway radius (± SEM) from $His^0$ and $His^+$ simulations.

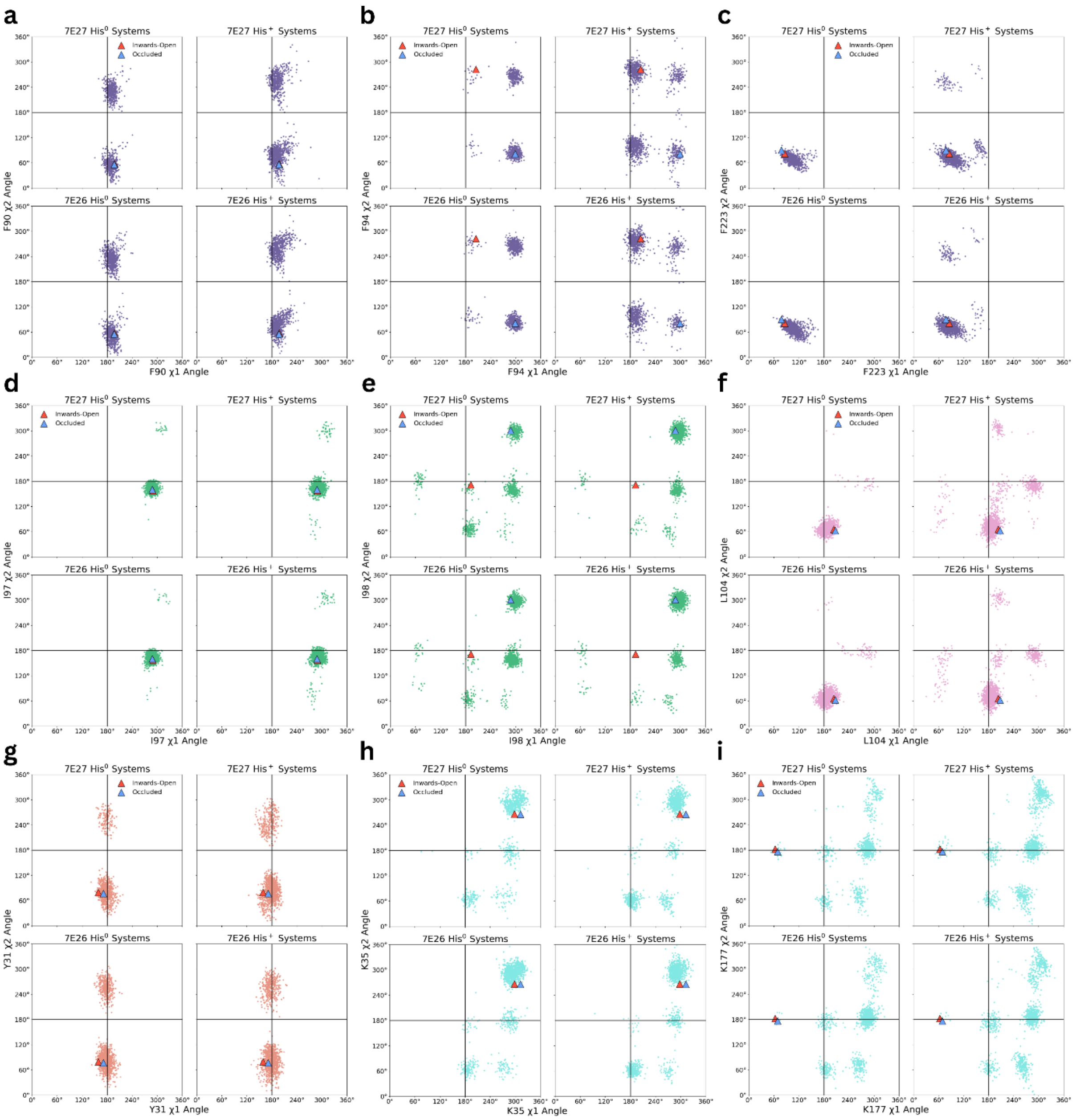


**Figure S3. χ1 and χ2 dihedral angles of key residues from different PfFNT systems.** The dihedral angles for **a)** F90, **b)** F94, **c)** F223, **d)** I97, **e)** I98, **f)** L104, **g)** Y31, **h)** K35, and **i)** K177 from $His^0$ and $His^+$ systems, starting from either the 7E26 or 7E27 structures.

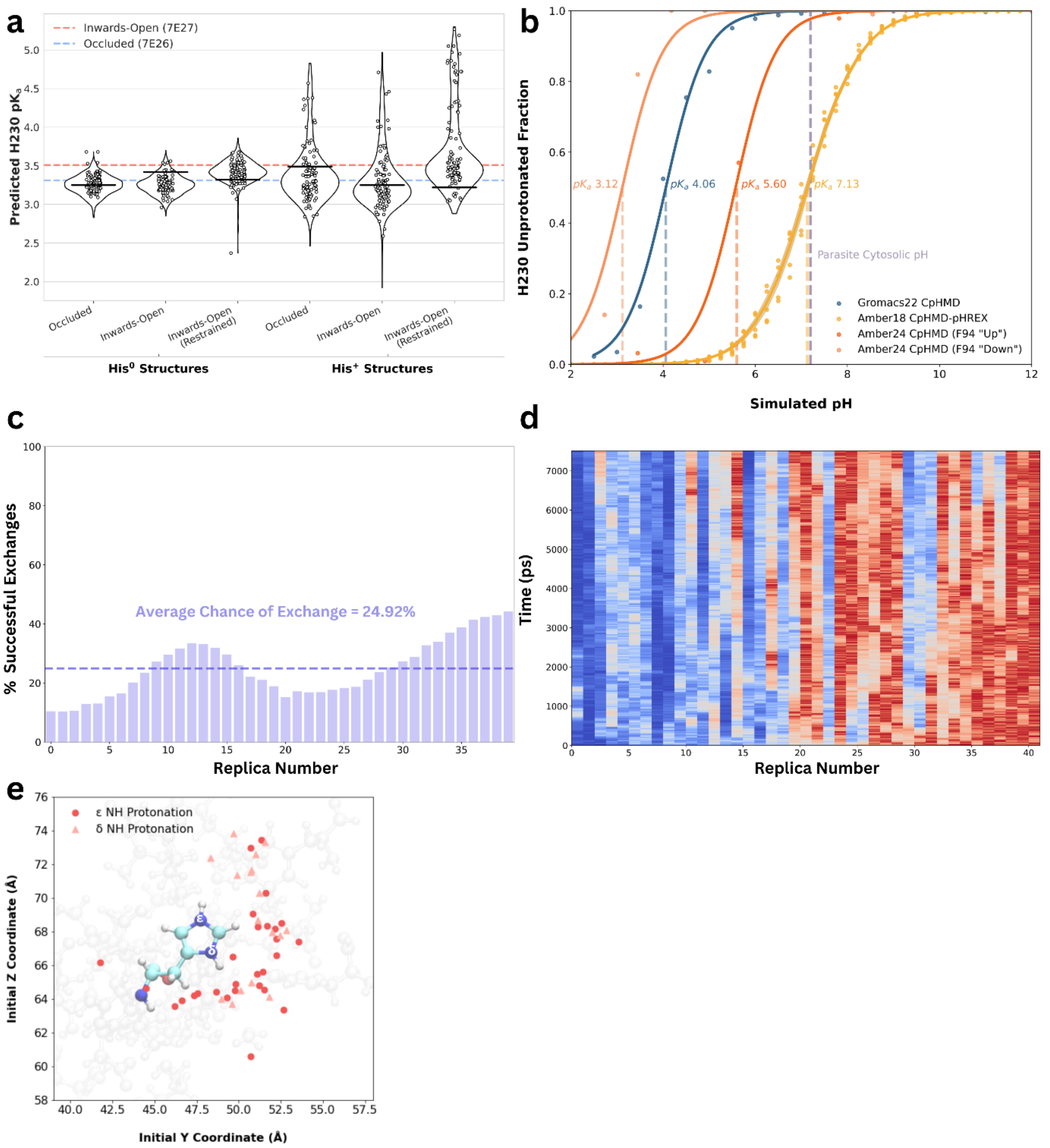


**Figure S4. H230 protonation is plausible, but p$K_a$ predictions are inconsistent. a)** H230 p$K_a$ values predicted using PROPKA. PROPKA calculations were performed on the cryo-EM structures (dashed lines) as well as on frames taken from MD simulations under different conditions in the absence of substrates. **b)** H230 p$K_a$ values predicted using all-atom CpHMD simulations in Gromacs22 and Amber24, as well as CpHMD pH replica exchange simulations (CpHMD-pHREX) performed in implicit solvent in Amber18. The p$K_a$ obtained from each method is shown next to the corresponding titration curve. A dashed purple line at pH 7.2 is used to indicate parasite cytosolic pH. **c)** Average exchange frequency over time between adjacent replicas in implicit solvent CpHMD-pHREX simulations. Exchange frequency was determined by measuring the number of successful exchanges between each window out of the total number of exchanges attempted. A dashed line is added to represent the overall average chance of exchange between replicas (~24.92%). **d)** 2D heatmap showing exchange between replicas over time. **e)** Starting positions for protons from DFTB simulations.

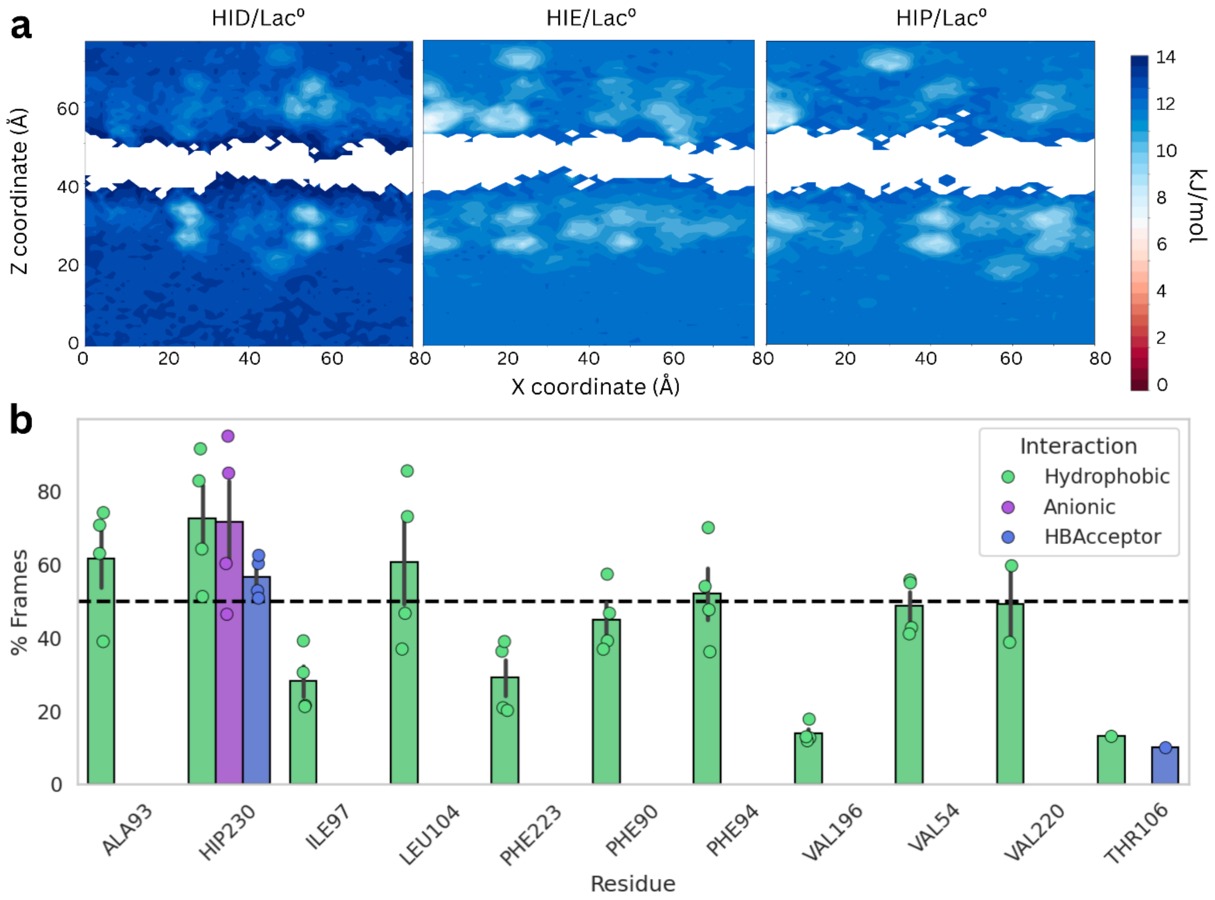


**Figure S5. Lactic acid does not enter the cavity, but lactate binds to charged H230. a)** Free energy surfaces for lactic acid for different H230 protonation states. HIE = neutral and protonated at the ε nitrogen, HID = neutral and protonated at the δ nitrogen, HIP = protonated at both. **b)** The average residue contacts observed across all lactate binding events were determined using ProLIF and the percentage of frames that each residue contacted substrates were averaged across all binding events for that substrate. Only frames where the substrate was inside the transport cavity were included for analysis. Residues that interact with substrates are coloured by the type of interaction occurring. A dashed line is added to all plots to indicate which residues interact with the respective substrate in over 50% of all frames. Error bars represent standard error.

**Table S2. Binding events observed across all simulations.** The total number of binding events (out of a possible five) observed in each replicate for the lactate, formate, nitrate, and iodide $His^+$ systems is shown below. Other = all other compounds ($citrate^{2-}$, $citrate^{3-}$, lactamide, propionamide, acetamide, formamide) with either $His^+$ or $His^0$, in which no binding events were observed. In the case of co-flooding simulations, binding events are listed for $Lac^-$, only.

| **Replicate** | **$Lac^-/His^+$** | **$For^-/His^+$** | **$Nit^-/His^+$** | **$I^-/His^+$** | **$Lac^-$ & $I^-/His^+$** | **$Lac^-$ & $Lam^0/His^+$** | **Other*** |
|---|---|---|---|---|---|---|---|
| 1 | 3 | 3 | 3 | 4 | 0 | 2 | 0 |
| 2 | 3 | 4 | 4 | 5 | 0 | 1 | 0 |
| 3 | 2 | 3 | 3 | 4 | 0 | 2 | 0 |
| 4 | 3 | 4 | - | - | 0 | 2 | - |
| 5 | 3 | 3 | - | - | 0 | 2 | - |
| 6 | 2 | 3 | - | - | - | - | - |
| 7 | 3 | - | - | - | - | - | - |
| 8 | 3 | - | - | - | - | - | - |
| **Mean** | 2.75 | 3.33 | 3.33 | 4.33 | 0 | 1.80 | 0 |
| **SEM** | 0.15 | 0.21 | 0.21 | 0.21 | 0 | 0.18 | 0 |

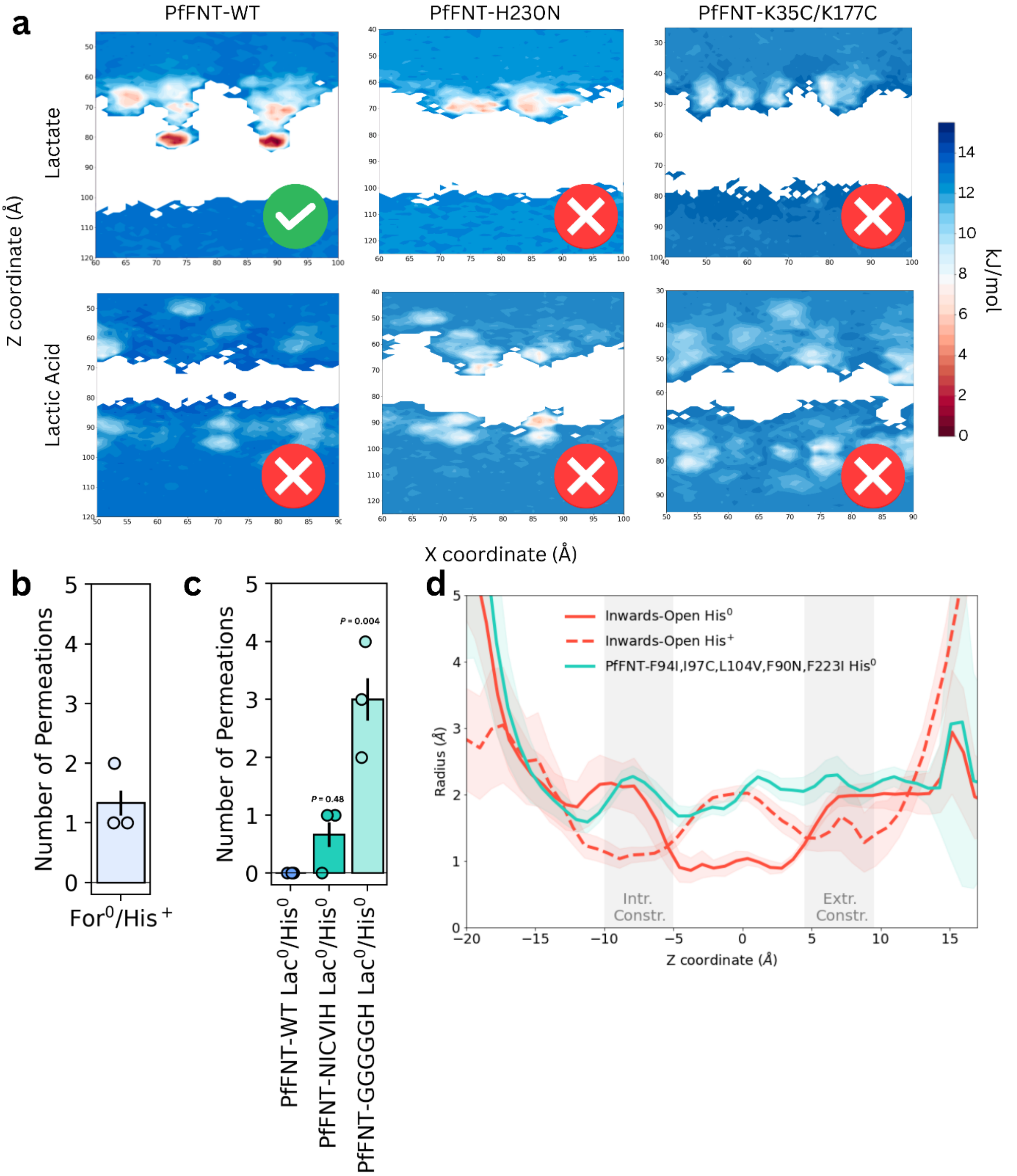


**Figure S6. Mutations to key residues alter binding and transport in simulations. a)** Free energy surfaces for Lac$^-$ and Lac$^0$ from PfFNT-WT, PfFNT-H230N, and PfFNT-K35C/K177C simulations. Green ticks indicate systems where substrate binding occurred, while red crosses indicate systems where binding did not occur. H230 was charged in all systems (except for H230N where it had been mutated to asparagine). **b)** Average number of permeation events for the For$^0$/His$^+$ system (± SEM). **c)** Average permeation events for Lac$^0$ in PfFNT-WT and mutant systems. PfFNT-NICVIH = PfFNT-F94I/I97C/L104V/F90N/F223I mutant (± SEM). PfFNT-GGGGGH = PfFNT-F94G/I97G/L104G/F90G/F223G mutant. **d)** Average transport pathway radius of PfFNT-WT systems and the PfFNT-F94I/I97C/L104V/F90N/F223I mutant.

**Table S3. Permeation events from flooding simulations.** The number of permeation events measured from flooding simulations for formic acid with wildtype PfFNT with charged H230 ($For^0/His^+$), as well as for lactic acid in mutant simulations (PfFNT-GGGGGH = PfFNT-F94G/I97G/L104G/F90G/F223G mutant, PfFNT-NICVIH = PfFNT-F94I/I97C/L104V/F90N/F223I mutant) across different replicates. In both mutant systems, H230 was neutral ($His^0$).

| **Replicate** | **$For^0/His^+$** | **$Lac^0$/PfFNT-GGGGGH $His^0$** | **$Lac^0$/PfFNT-NICVIH $His^0$** |
|---|---|---|---|
| 1 | 2 | 3 | 1 |
| 2 | 1 | 4 | 1 |
| 3 | 1 | 3 | 0 |
| **Mean** | 1.33 | 3.33 | 0.66 |
| **SEM** | 0.21 | 0.21 | 0.21 |

**Table S4. Dissociation events observed from lactate and lactic acid systems.** The total number of dissociation events (out of a possible five) observed in each replicate of each system is summarized for the lactate and lactic acid simulations. A dissociation event was counted if the centre of mass of the substrate passed the centre of mass of either constriction to exit the transport cavity. The direction of dissociation (intracellular or extracellular) is recorded next to the number of dissociation events observed within the simulation.

| | **Number of Dissociation Events** | | | | |
|---|---|---|---|---|---|
| **Replicate** | **$Lac^-/His^+$** | **$Lac^0/His^+$** | **$Lac^-/His^0$** | **$Lac^0/His^0$** | **$Lac^0/His^0$ (Proton Transfer)** |
| 1 | 0 | 1 (intracellular) | 3 (intracellular) | 0 | 2 (2 intracellular) |
| 2 | 0 | 0 | 5 (intracellular) | 0 | 4 (1 intracellular, 3 extracellular) |
| 3 | 0 | 2 (intracellular) | 1 (intracellular) | 1 (extracellular) | 0 |
| 4 | 0 | 2 (1 intracellular, 1 extracellular) | 3 (intracellular) | 0 | 1 (1 extracellular) |
| 5 | 0 | 2 (intracellular) | 4 (intracellular) | 0 | 2 (1 intracellular, 1 extracellular) |
| 6 | 0 | 1 (intracellular) | 4 (intracellular) | 1 (intracellular) | 2 (intracellular) |
| 7 | 0 | 3 (2 intracellular, 1 extracellular) | 5 (intracellular) | 1 (extracellular) | 1 (intracellular) |
| 8 | 0 | 4 (1 intracellular, 3 extracellular) | 5 (intracellular) | 0 | 5 (4 intracellular, 1 extracellular) |
| 9 | 0 | 4 (2 intracellular, 2 extracellular) | 3 (intracellular) | 3 (2 intracellular, 1 extracellular) | 5 (3 intracellular, 2 extracellular) |
| **Mean** | 0 | 2.11 | 3.67 | 0.67 | 2.44 |
| **SEM** | 0 | 0.45 | 0.44 | 0.33 | 0.60 |

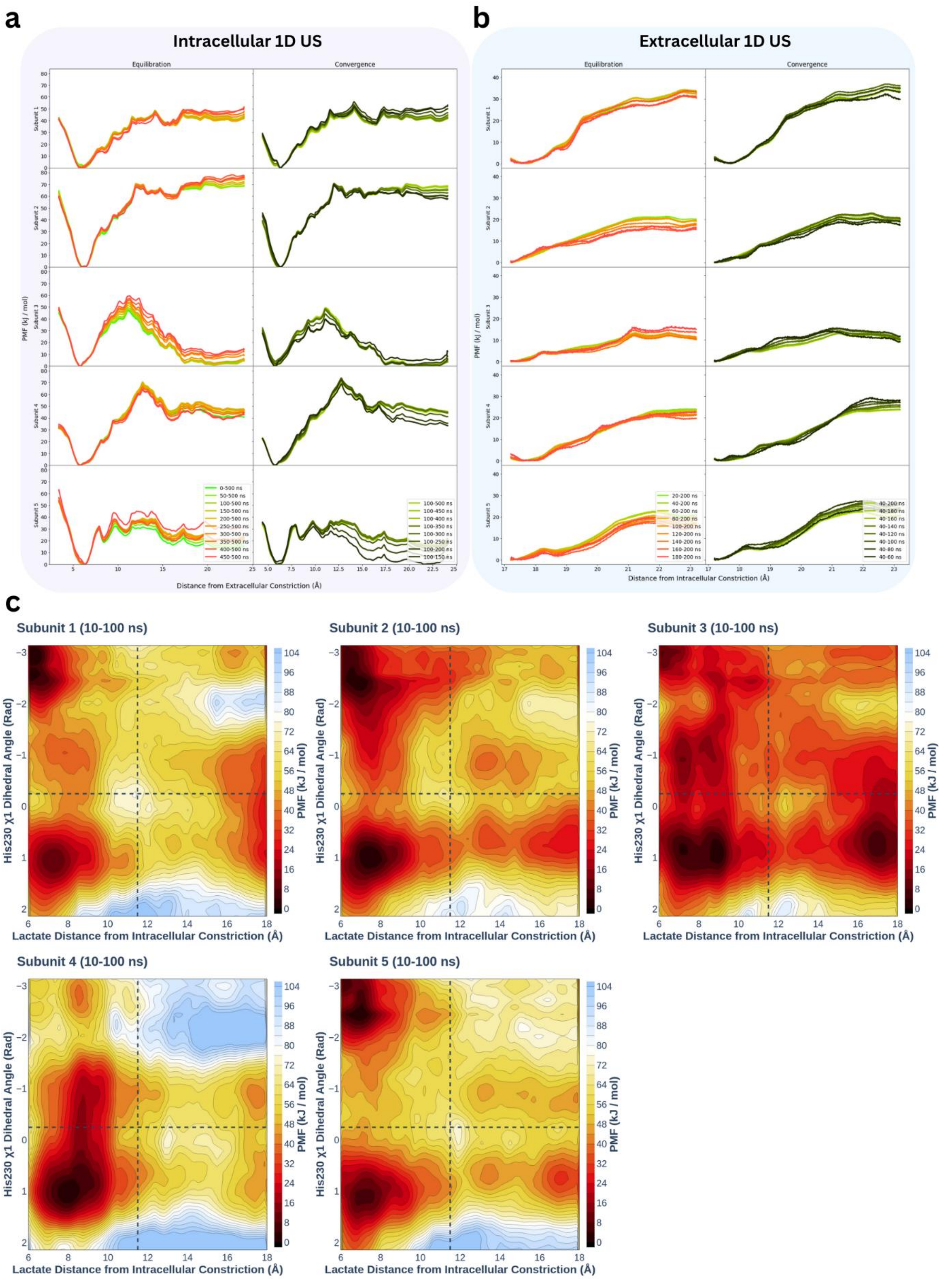


**Figure S7. Convergence and reproducibility of Lac$^-$/His$^+$ intracellular and extracellular umbrella sampling.** PMFs for each of the five subunits for the **a)** intracellular and **b)** extracellular 1D umbrella sampling constructed from different lengths of simulation time are graphed together to show how much simulation time is required for the system to equilibrate and converge. The Y-axis represents the potential of mean force (PMF) in kJ/mol as the substrate moves through the protein. For the intracellular side, the X-axis represents the distance from the extracellular constriction site (Å) as lactate moves from the bulk solution into the transport cavity. For the extracellular side, the X-axis represents the distance from the intracellular constriction site (Å) as lactate moves from the bulk solution into the transport cavity. **c)** 2D umbrella sampling was performed where the distance between lactate and the backbone atoms of the intracellular constriction, as well as the $\chi 1$ dihedral angle of the H230 sidechain, were biased. PMFs include data obtained between 10 and 100 ns of simulation, with the first 10 ns having been discarded as equilibration time. In all subunits, an energy minimum can be observed inside the cavity when the H230 sidechain is angled downwards ($x < 10$, $y > 1$). An additional minimum can be observed in subunits 1, 2, 3, and 5 where the H230 sidechain is angled upwards ($x < 10$, $y < -2$).

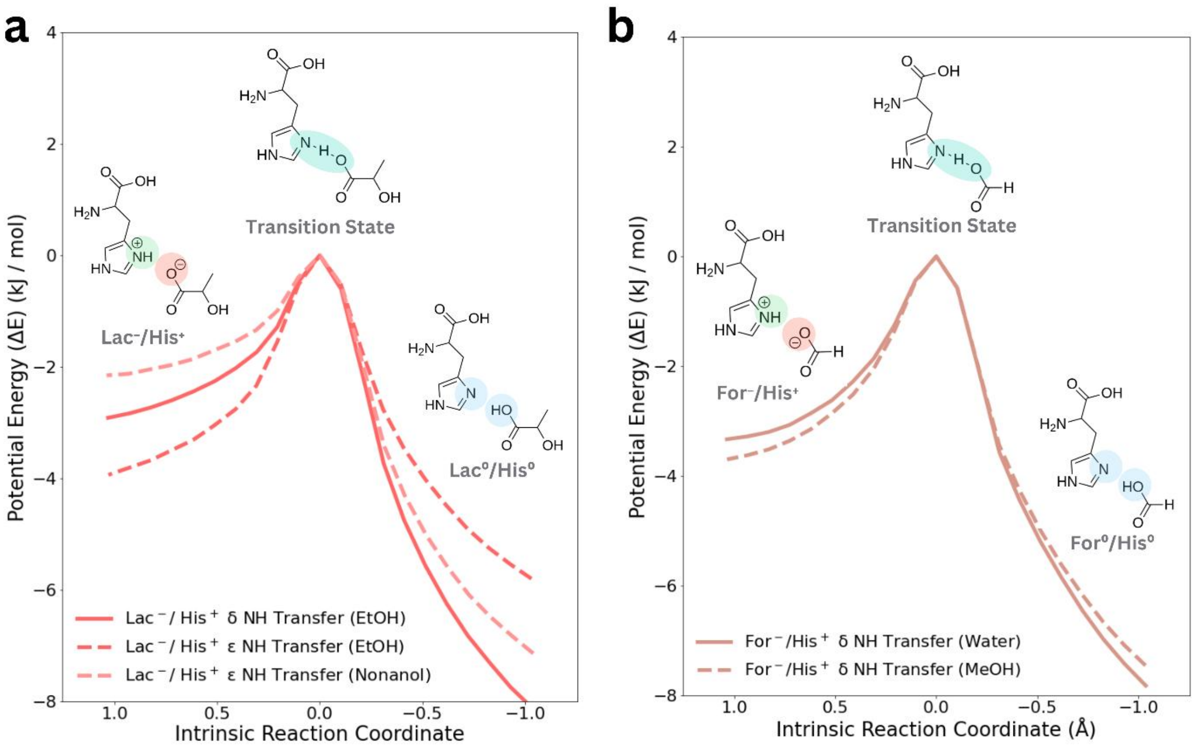


**Figure S8. Proton transfer can occur under different solvent conditions. a)** Energetics of $His^{+}$ donating a proton to $Lac^{-}$ in implicit ethanol and nonanol environments. **b)** Energetics of $His^{+}$ donating a proton to $For^{-}$ in implicit water or MeOH environments.

**Table S5. Summary of quantum chemical calculations performed in this study.** Quantum chemical calculations performed using SOGGA11X(D3-BJ)/def2-TZVPP to determine whether proton transfer can occur from His+ to lactate, formate, nitrate, and MMV007839 in the presence and absence of different solvents. Additional calculations were performed to determine whether H3O+ can donate a proton to His0.

| Substrate | PCM solvent | OPT (H-His start) | OPT (H-substrate start) | QST2 |
|---|---|---|---|---|
| **Lactate-δHis** | None | Proton Transfer from His+ to Lac- (5 of 5 replicates) | - | Barrierless |
| | MeOH | Proton Transfer from His+ to Lac- (5 of 5 replicates) | - | Barrierless |
| | Water | No transfer (5 of 5 replicates) | Lac0 stays protonated (5 of 5 replicates) | Transition state found |
| **Lactate-εHis** | None | Proton Transfer from His+ to Lac- (3 of 5 replicates) | - | Barrierless |
| | MeOH | Proton Transfer from His+ to Lac- (3 of 5 replicates) | - | Barrierless |
| | Water | No Transfer (5 of 5 replicates) | Lac0 stays protonated (5 of 5 replicates) | Transition state found |
| **Formate- δHis** | None | Proton Transfer from His+ to For- (4 of 4 replicates) | - | Barrierless |
| | Water | No Transfer (5 of 5 replicates) | For0 stays protonated (5 of 5 replicates) | Transition state found |
| **Nitrate- δHis** | None | Proton Transfer from His+ to Nit- (3 of 3 replicates) | - | Barrierless |
| | MeOH | No Transfer (3 of 3 replicates) | - | Barrierless |
| | Water | No Transfer (5 of 5 replicates) | Nit0 transfers proton back to His0 (5 of 5 replicates) | Barrierless |
| **Iodide-εHis** | None | No Transfer (5 of 5 replicates) | - | No TS found |
| | Water | No Transfer (5 of 5 replicates) | - | No TS found |
| **MMV007839-εHis** | None | Proton Transfer from His+ to MMV007839- (2 of 5 replicates) | - | No TS found |
| | Water | No Transfer (5 of 5 replicates) | MMV007839$^{0}$ transfers proton | No TS found |

| | | | |
|---|---|---|---|
| | | | back to His0 (5 of 5 replicates) |
| $H_3O^+$ | None | - | $H_3O^+$ transfers proton to His0 (1 of 1 replicate) |
| | MeOH | - | $H_3O^+$ transfers proton to His0 (1 of 1 replicate) |
| | Water | His+ stays protonated (1 of 1 replicate) | No Transfer (1 of 1 replicate) |

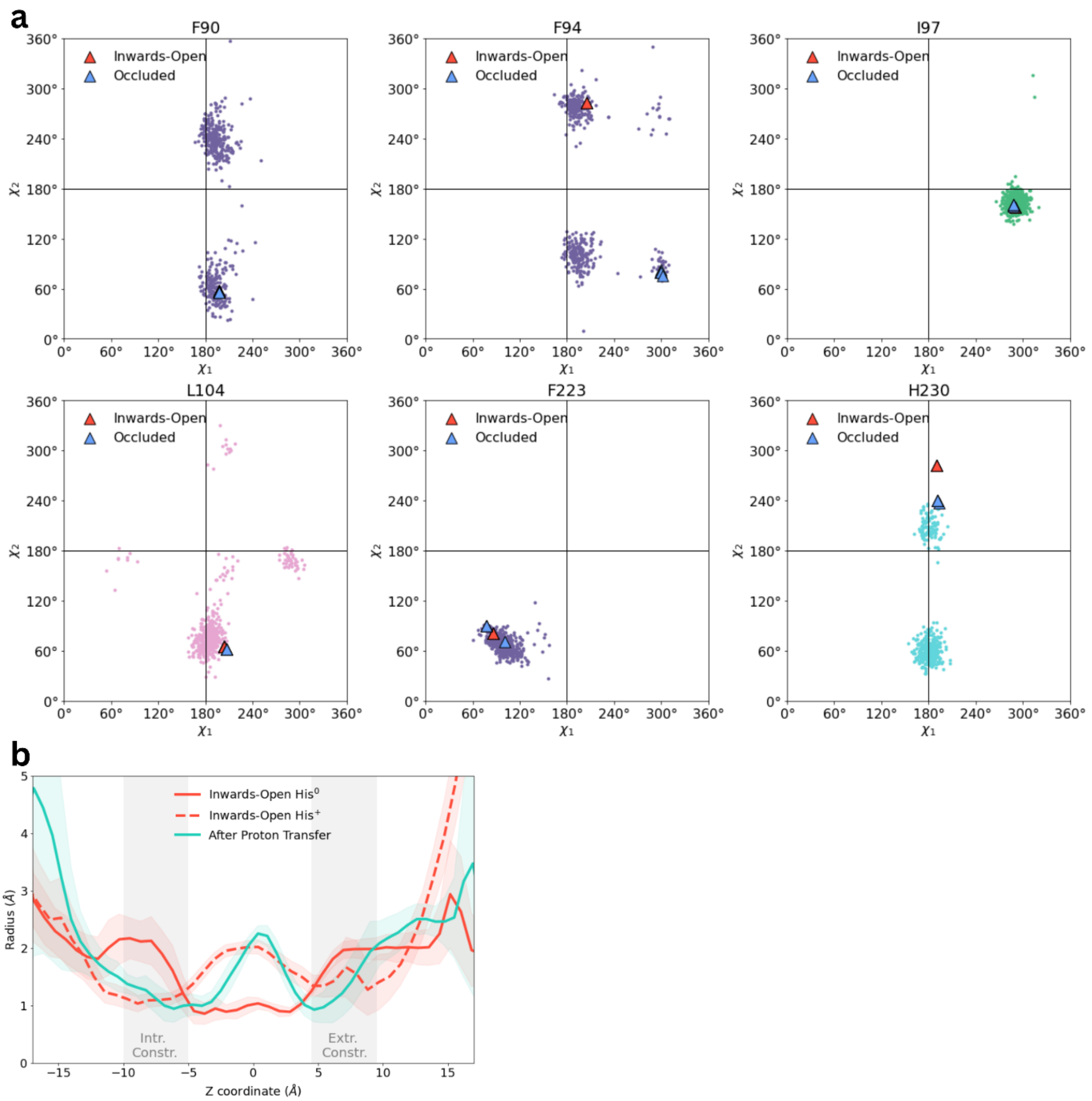


**Figure S9. The transport cavity remains open following proton transfer. a)** χ1- χ2 Janin plots for residues surrounding the PfFNT transport cavity. Janin plots are shown for F90, F94, I97, L104, F223, and H230 following proton transfer. Data from all five subunits and all three replicates for both inwards-open and occluded systems were combined on the same plot. Triangles are added to represent the starting dihedral angle from the inwards-open (red) and occluded (blue) cryo-EM structures. **b)** Measurements of average pore radius following proton transfer. The average pore radius (± SEM) across each frame of the ensemble trajectory of all subunits and replicates was measured using HOLE. Average radius for the Inwards-Open $His^0$ and Inwards-Open $His^+$ systems are shown for reference.

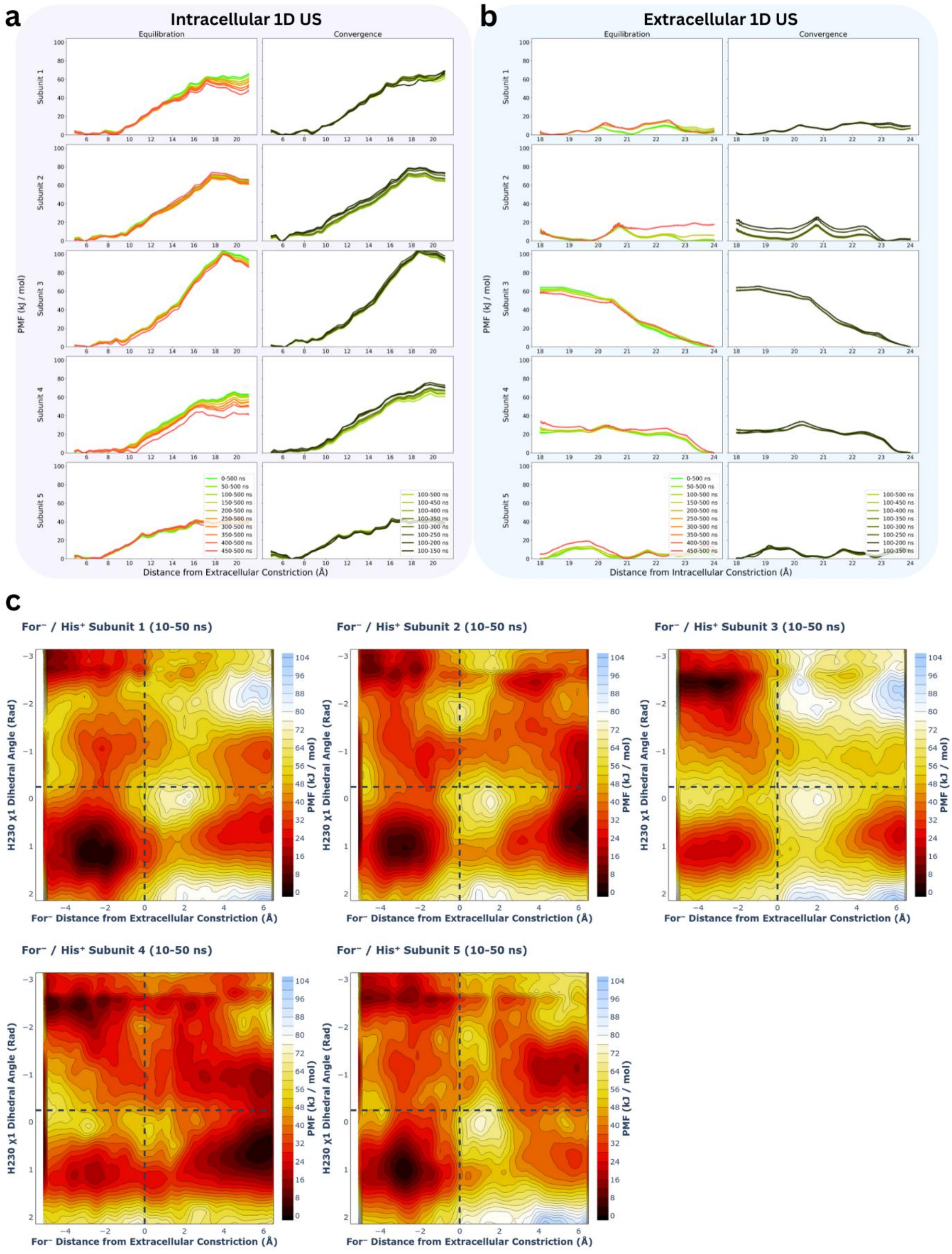


**Figure S10. Convergence and reproducibility of $For^-/His^+$ intracellular and extracellular umbrella sampling.** PMFs for each of the five subunits for the **a)** intracellular and **b)** extracellular 1D umbrella sampling constructed from different lengths of simulation time are graphed together to show how much simulation time is required for the system to equilibrate and converge. The Y-axis represents the potential of mean force (PMF) in kJ/mol as the substrate moves through the protein. For the intracellular side, the X-axis represents the distance from the extracellular constriction site (Å) as formate moves from the bulk solution into the transport cavity. For the extracellular side, the X-axis represents the distance from the intracellular constriction site (Å) as formate moves from the bulk solution into the transport cavity. **c)** 2D umbrella sampling was performed where the distance between formate and the backbone atoms of the intracellular constriction, as well as the $\chi 1$ dihedral angle of the H230 sidechain, were biased. PMFs include data obtained between 10 and 100 ns of simulation, with the first 10 ns having been discarded as equilibration time.

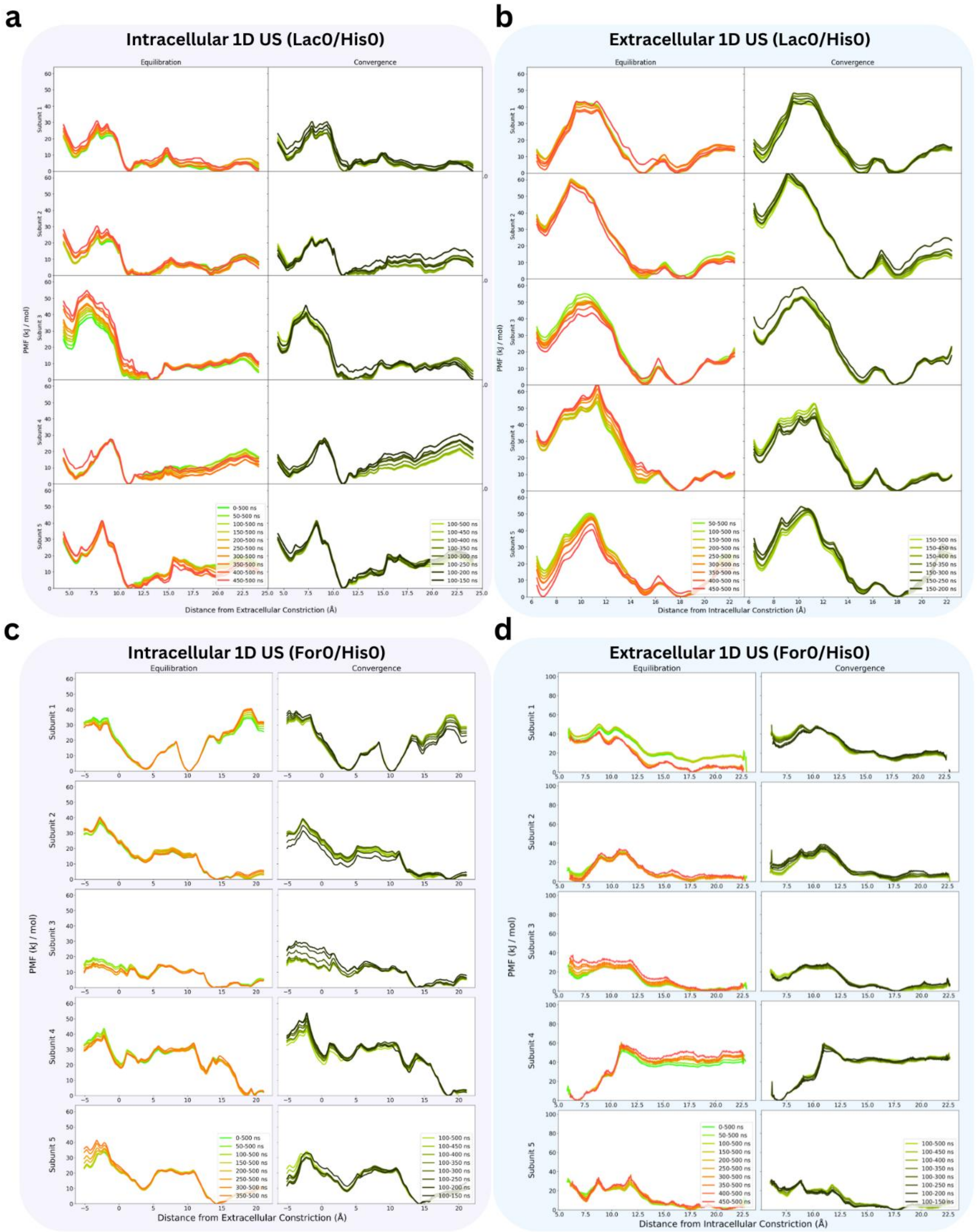


**Figure S11. Convergence and reproducibility of the 1D Lac$^0$/His$^0$ and For$^0$/His$^0$ intracellular and extracellular umbrella sampling.** PMFs for each of the five subunits for Lac$^0$/His$^0$ **a)** intracellular and **b)** extracellular, and For$^0$/His$^0$ **c)** intracellular and **d)** extracellular umbrella sampling constructed from different lengths of simulation time are graphed together to show how much simulation time is required for the system to equilibrate and converge. The Y-axis represents the potential of mean force (PMF) in kJ/mol as the substrate moves through the protein. For the intracellular side, the X-axis represents the distance from the extracellular constriction site (Å) as the substrate moves from the bulk solution into the transport cavity. For the extracellular side, the X-axis represents the distance from the intracellular constriction site (Å) as the substrate moves from the bulk solution into the transport cavity.

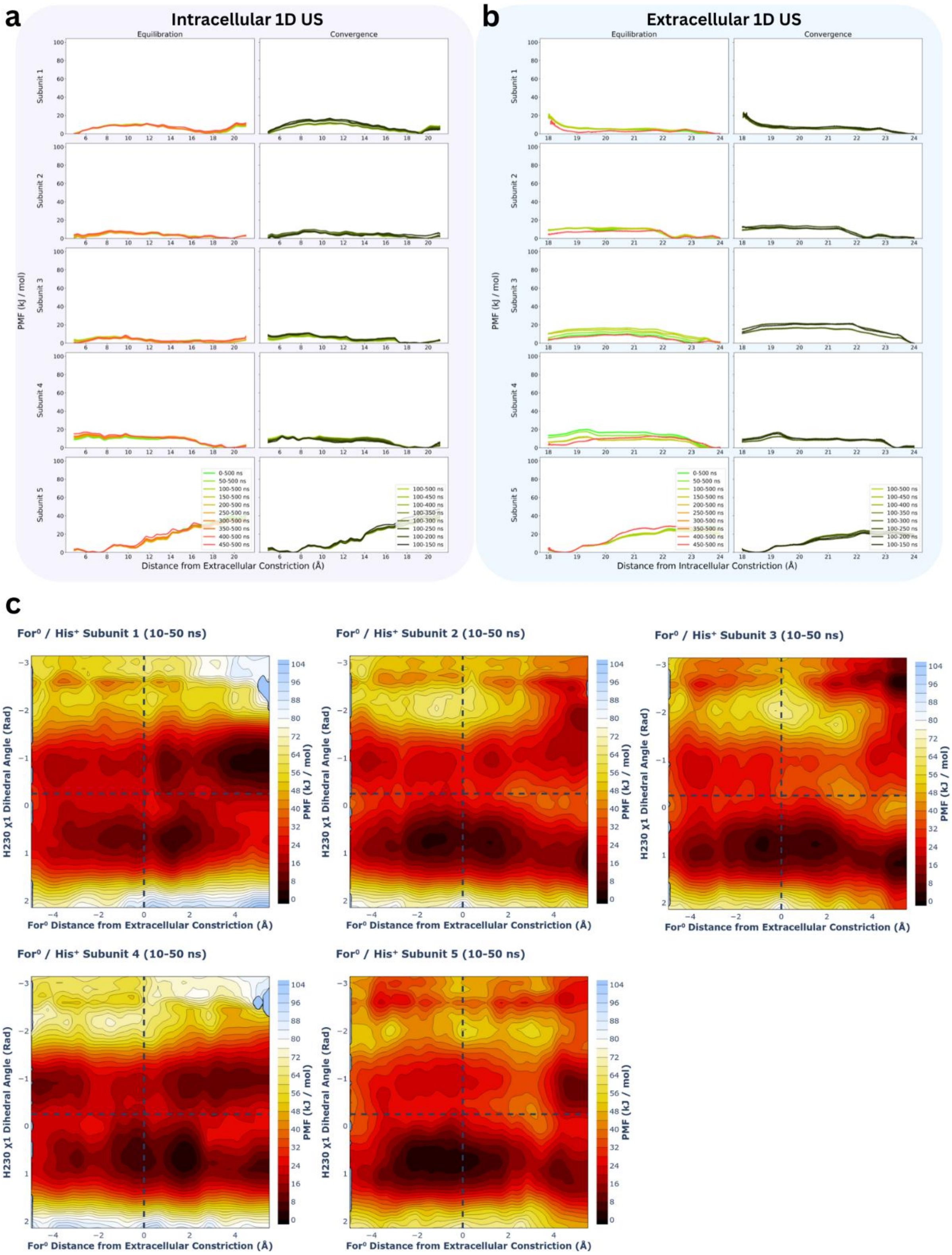


**Figure S12. Convergence and reproducibility of For$^0$/His$^+$ intracellular and extracellular umbrella sampling.** PMFs for each of the five subunits for the **a)** intracellular and **b)** extracellular 1D umbrella sampling constructed from different lengths of simulation time are graphed together to show how much simulation time is required for the system to equilibrate and converge. The Y-axis represents the potential of mean force (PMF) in kJ/mol as the substrate moves through the protein. For the intracellular side, the X-axis represents the distance from the extracellular constriction site (Å) as formic acid moves from the bulk solution into the transport cavity. For the extracellular side, the X-axis represents the distance from the intracellular constriction site (Å) as formic acid moves from the bulk solution into the transport cavity. **c)** 2D umbrella sampling was performed where the distance between formic acid and the backbone atoms of the intracellular constriction, as well as the $\chi 1$ dihedral angle of the H230 sidechain, were biased. PMFs include data obtained between 10 and 100 ns of simulation, with the first 10 ns having been discarded as equilibration time.

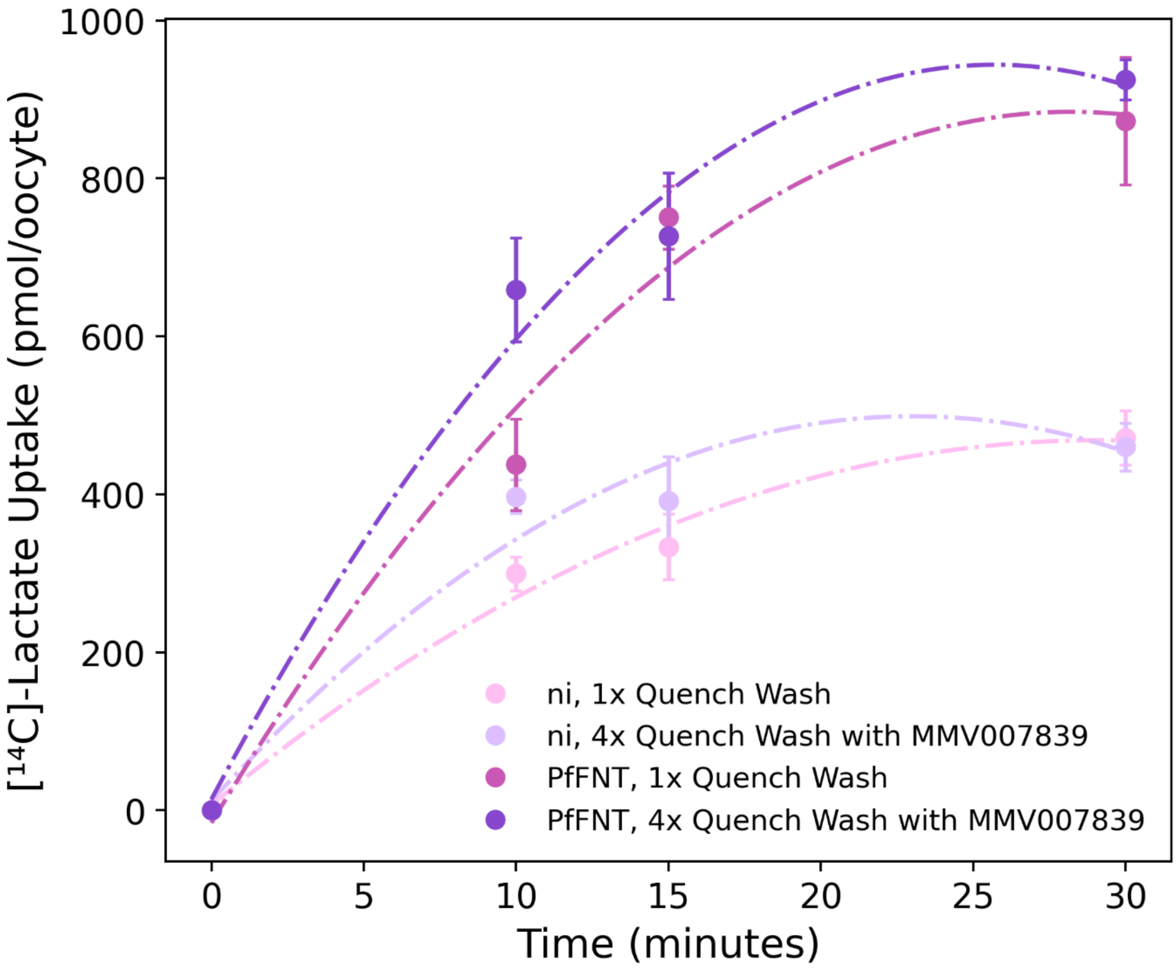


**Figure S13. Timecourses for [$^{14}$C]-lactate uptake in non-injected and PfFNT-injected oocytes.** Experiments were performed with PfFNT-expressing (PfFNT) and non-injected (ni) oocytes, using an extracellular lactate concentration of 1 mM at pH 6.4 at 27.5 °C (n=3). Each replicate used 7-10 individual oocytes per timepoint per condition. Experiments were performed on separate days, using oocytes from separate frogs. The quench washes for non-injected (ni) and PfFNT-injected (PfFNT) oocytes were performed either by aspirating radiolabelled substrate after the desired incubation period and washing once in ice-cold ND96 (1x Quench Wash) or by washing four times in ice-cold ND96 containing 2 µM MMV007839 (4x Quench Wash).

**Table S6. [$^{14}$C]-lactate uptake by non-injected and PfFNT-expressing oocytes from competition assays.** Data are presented for each individual repeat (corresponding to independent experimental days). The test compounds included in each repeat are listed. All conditions contained 1 mM lactate; N/A denotes the control condition in which only 1 mM lactate was present. For each repeat, the average [$^{14}$C]-lactate uptake across 8–10 non-injected oocytes and across PfFNT-expressing oocytes is shown and was used to calculate the PfFNT-induced uptake, defined as the uptake in PfFNT-expressing oocytes minus that in non-injected oocytes. The PfFNT-induced uptake (% of control) was calculated for each repeat relative to the N/A condition, where the average PfFNT-induced uptake in the presence of each test compound was divided by the average PfFNT-induced uptake under the N/A control.

| Repeat | Test Compound | Uptake (pmol/oocyte/15 min) | | | |
|---|---|---|---|---|---|
| | | Average Non-Injected Uptake | Average PfFNT-Expressing Uptake | Average PfFNT-Induced Uptake | PfFNT-induced uptake (% of Control) |
| 1 | N/A | 107.63 | 375.05 | 267.42 | 100.00 |
| 1 | 10 mM Pyruvate | 76.65 | 148.16 | 71.51 | 26.74 |
| 1 | 10 mM Propionate | 103.33 | 217.25 | 113.92 | 42.60 |
| 1 | 10 mM Acetate | 116.59 | 205.98 | 89.39 | 33.43 |
| 1 | 10 mM Formate | 103.15 | 141.50 | 38.34 | 14.34 |
| 1 | 10 mM Citrate | 106.06 | 144.67 | 38.61 | 14.44 |
| 1 | 10 mM Nitrate | 140.39 | 209.74 | 69.35 | 25.93 |
| 1 | 10 mM Iodide | 128.54 | 88.07 | -40.47 | -15.13 |
| 1 | 10 mM Lactamide | 199.44 | 308.58 | 109.14 | 40.81 |
| 1 | 10 mM Propionamide | 134.50 | 101.41 | -33.08 | -12.37 |
| 1 | 10 mM Acetamide | 188.81 | 288.91 | 100.10 | 37.43 |
| 1 | 10 mM Formamide | 134.39 | 161.35 | 26.95 | 10.08 |
| 1 | 2 µM MMV007839 | 92.84 | 67.14 | -25.70 | -9.61 |
| 2 | N/A | 209.66 | 509.12 | 299.45 | 100.00 |
| 2 | 10 mM Pyruvate | 189.77 | 243.09 | 53.32 | 17.81 |
| 2 | 10 mM Propionate | 173.57 | 236.17 | 62.61 | 20.91 |
| 2 | 10 mM Acetate | 263.81 | 323.77 | 59.97 | 20.03 |
| 2 | 10 mM Formate | 234.28 | 312.13 | 77.84 | 25.99 |
| 2 | 10 mM Citrate | 255.74 | 264.00 | 8.25 | 2.76 |

| | | | | | |
|---|---|---|---|---|---|
| 2 | 10 mM Nitrate | 254.42 | 344.85 | 90.44 | 30.20 |
| 2 | 10 mM Iodide | 189.60 | 166.27 | -23.32 | -7.79 |
| 2 | 10 mM Lactamide | 168.33 | 374.34 | 206.01 | 68.79 |
| 2 | 10 mM Propionamide | 185.08 | 248.70 | 63.62 | 21.24 |
| 2 | 10 mM Acetamide | 225.31 | 272.68 | 47.37 | 15.82 |
| 2 | 10 mM Formamide | 182.05 | 209.46 | 27.41 | 9.15 |
| 2 | 2 µM MMV007839 | 192.84 | 199.95 | 7.10 | 2.37 |
| 3 | N/A | 127.99 | 419.91 | 291.92 | 100.00 |
| 3 | 10 mM Pyruvate | 120.05 | 219.91 | 99.86 | 34.21 |
| 3 | 10 mM Propionate | 105.08 | 183.12 | 78.04 | 26.73 |
| 3 | 10 mM Acetate | 155.32 | 189.72 | 34.40 | 11.78 |
| 3 | 10 mM Formate | 136.51 | 227.36 | 90.85 | 31.12 |
| 3 | 10 mM Citrate | 139.32 | 149.26 | 9.95 | 3.41 |
| 3 | 10 mM Nitrate | 124.14 | 210.28 | 86.14 | 29.51 |
| 3 | 10 mM Iodide | 128.34 | 140.46 | 12.12 | 4.15 |
| 3 | 10 mM Lactamide | 147.05 | 244.60 | 97.55 | 33.42 |
| 3 | 10 mM Propionamide | 103.90 | 199.27 | 95.37 | 32.67 |
| 3 | 10 mM Acetamide | 136.80 | 163.59 | 26.80 | 9.18 |
| 3 | 10 mM Formamide | 155.99 | 205.38 | 49.39 | 16.92 |
| 3 | 2 µM MMV007839 | 215.89 | 197.34 | -18.55 | -6.36 |
| 4 | N/A | 117.70 | 225.21 | 107.51 | 100.00 |
| 4 | 10 mM Pyruvate | 66.05 | 94.55 | 28.50 | 26.51 |
| 4 | 10 mM Propionate | 92.95 | 124.27 | 31.32 | 29.13 |
| 4 | 10 mM Acetate | 82.10 | 102.18 | 20.08 | 18.68 |
| 4 | 10 mM Formate | 78.89 | 94.70 | 15.81 | 14.71 |
| 4 | 10 mM Citrate | 108.42 | 135.33 | 26.92 | 25.04 |

| | | | | | |
|---|---|---|---|---|---|
| 4 | 10 mM Nitrate | 108.48 | 125.89 | 17.41 | 16.19 |
| 4 | 10 mM Iodide | 132.58 | 116.90 | -15.68 | -14.58 |
| 4 | 10 mM Lactamide | 94.20 | 151.45 | 57.25 | 53.25 |
| 4 | 10 mM Propionamide | 122.21 | 153.56 | 31.35 | 29.16 |
| 4 | 10 mM Acetamide | 91.04 | 106.07 | 15.03 | 13.98 |
| 4 | 10 mM Formamide | 111.59 | 120.23 | 8.64 | 8.03 |
| 4 | 2 µM MMV007839 | 107.94 | 108.75 | 0.80 | 0.75 |
| 5 | N/A | 199.16 | 464.89 | 265.73 | 100.00 |
| 5 | 10 mM Pyruvate | 171.59 | 281.78 | 110.18 | 41.47 |
| 5 | 10 mM Propionate | 217.06 | 348.14 | 131.08 | 49.33 |
| 5 | 10 mM Acetate | 212.27 | 410.58 | 198.31 | 74.63 |
| 5 | 10 mM Formate | 175.54 | 259.01 | 83.46 | 31.41 |
| 5 | 10 mM Citrate | 204.07 | 219.87 | 15.80 | 5.95 |
| 5 | 10 mM Nitrate | 171.32 | 207.57 | 36.24 | 13.64 |
| 5 | 10 mM Iodide | 225.74 | 204.80 | -20.94 | -7.88 |
| 5 | 10 mM Lactamide | 184.98 | 321.63 | 136.65 | 51.42 |
| 5 | 10 mM Propionamide | 175.99 | 272.67 | 96.68 | 36.38 |
| 5 | 10 mM Acetamide | 198.72 | 252.15 | 53.44 | 20.11 |
| 5 | 10 mM Formamide | 241.55 | 283.74 | 42.19 | 15.88 |
| 5 | 2 µM MMV007839 | 179.35 | 173.45 | -5.90 | -2.22 |

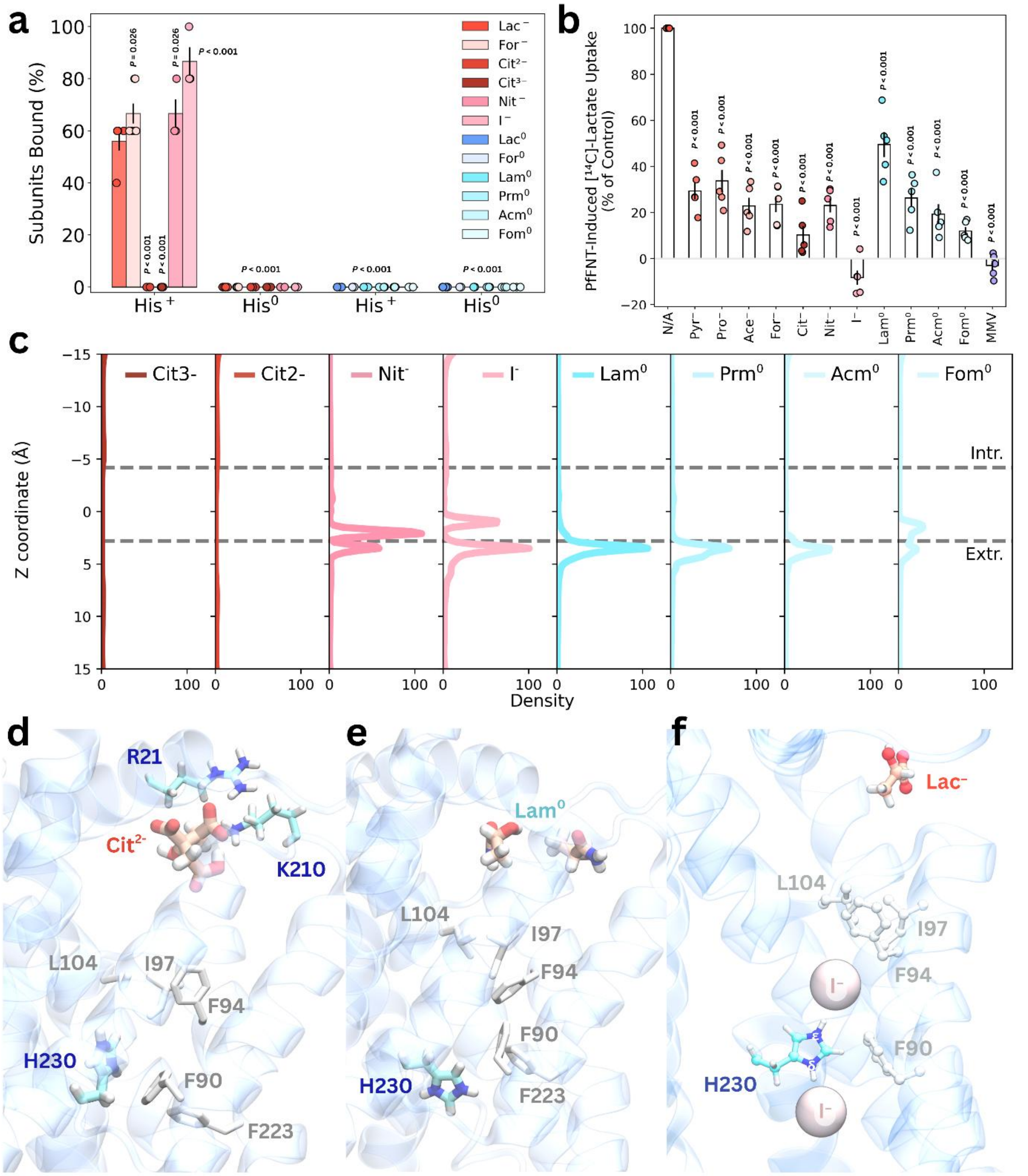


**Figure S14. Molecules tested in flooding simulations and X. laevis uptake assays.** **a)** Average binding events (± SEM) from flooding simulations with lactate (Lac⁻), formate (For⁻), citrate²⁻ (Cit²⁻), citrate³⁻ (Cit³⁻), nitrate (Nit⁻), iodide (I⁻), lactic acid (Lac⁰), formic acid (For⁰), lactamide (Lam⁰), propionamide (Prm⁰), acetamide (Acm⁰), and formamide (Fom⁰), where H230 was either neutral or charged. **b)** Effect of competing substrates on PfFNT-mediated lactate uptake (1 mM; of which a small portion was [$^{14}$C]-labelled) in X. laevis oocytes, from five independent experiments performed on different days using oocytes from different frogs. Uptake assays were performed at pH 6.4 with 1 mM lactate present in all conditions; test compounds were added at 10 mM (pyruvate, propionate, acetate, formate, citrate, nitrate, iodide, lactamide, propionamide, acetamide, formamide) or 2 μM (MMV007839). For each condition, uptake was measured over 15 minutes in both non-injected and PfFNT-injected oocytes, and the reported values represent PfFNT-induced uptake, defined as: (uptake in PfFNT-injected oocytes) - (uptake in non-injected oocytes). PfFNT-induced uptake is expressed as a percentage relative to the PfFNT-induced uptake measured in the absence of test compounds (N/A control). All compounds were added simultaneously with radiolabelled and unlabelled lactate. Each experiment used 8-10 oocytes per condition. Statistical significance was assessed using a one-way ANOVA blocking by experimental day, followed by a post hoc Tukey test. **c)** Average substrate density across z for $Cit^{2-}$, $Cit^{3-}$, $Nit^{-}$, $I^{-}$, $Lac^{0}$, $For^{0}$, $Lam^{0}$, $Prm^{0}$, $Acm^{0}$, and $Fom^{0}$ from flooding simulations. **d–f)** Representative snapshots of $Cit^{2-}$, $Lam^{0}$, and $I^{-}$ from flooding simulations.

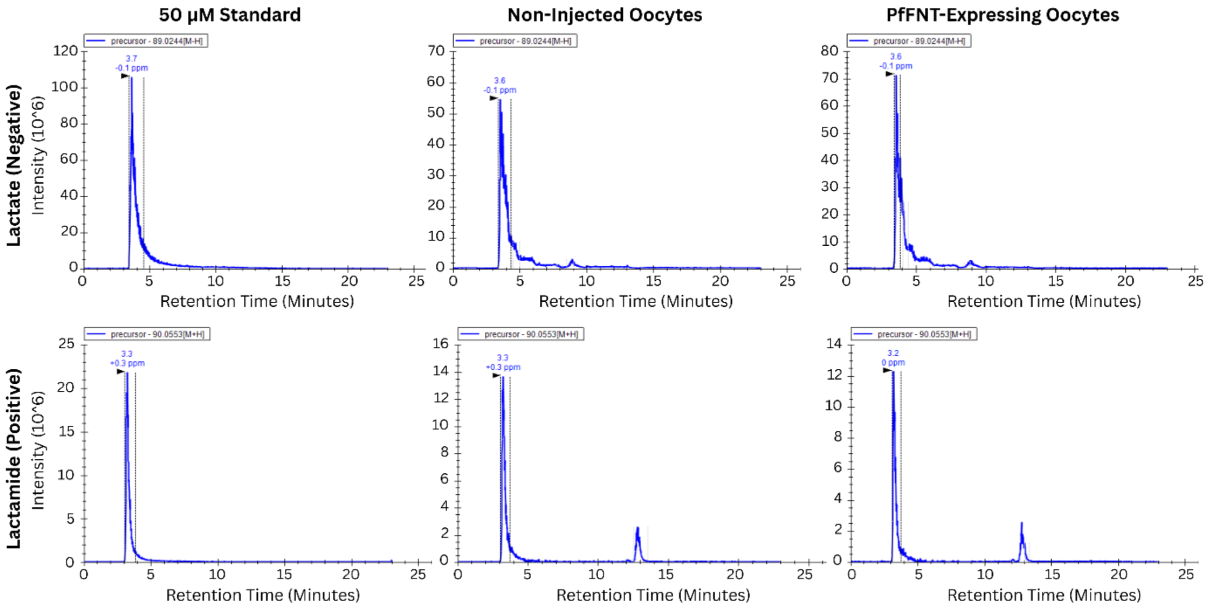


**Figure S15. Extracted ion chromatograms for lactate and lactamide from samples run in positive and negative electrospray modes.** Output files from LC-MS runs for standards of lactate and lactamide, as well as samples extracted from non-injected and PfFNT-expressing oocytes, were analysed in Skyline to obtain the extracted ion chromatograms. Peak areas for each chromatogram were extracted and analysed. Dashed vertical lines signify the borders of the peak used for calculating peak area. Blue numbers above each peak refer to the retention time of the compound. The key shows the expected size of the precursor mass used for peak identification within skyline.

**Table S7. Raw peak areas from extracted ion chromatograms for lactate and lactamide from non-injected and PfFNT-expressing oocyte samples.** Lactate data is reported from samples run in negative electrospray mode, while lactamide data is reported from samples run in positive electrospray mode.

| Compound | Oocyte | Electrospray Mode | Peak Area |
|---|---|---|---|
| Lactate | Non-injected | Negative | $8.21 \times 10^8$ |
| Lactate | Non-injected | Negative | $7.98 \times 10^8$ |
| Lactate | Non-injected | Negative | $8.54 \times 10^8$ |
| Lactamide | Non-injected | Negative | - |
| Lactamide | Non-injected | Negative | - |
| Lactamide | Non-injected | Negative | - |
| Lactate | PfFNT | Negative | $1.36 \times 10^9$ |
| Lactate | PfFNT | Negative | $1.25 \times 10^9$ |
| Lactate | PfFNT | Negative | $1.46 \times 10^9$ |
| Lactamide | PfFNT | Negative | - |
| Lactamide | PfFNT | Negative | - |
| Lactamide | PfFNT | Negative | - |
| Lactate | Non-injected | Positive | - |
| Lactate | Non-injected | Positive | - |
| Lactate | Non-injected | Positive | - |
| Lactamide | Non-injected | Positive | $1.53 \times 10^8$ |
| Lactamide | Non-injected | Positive | $1.48 \times 10^8$ |
| Lactamide | Non-injected | Positive | $1.55 \times 10^8$ |
| Lactate | PfFNT | Positive | - |
| Lactate | PfFNT | Positive | - |
| Lactate | PfFNT | Positive | - |
| Lactamide | PfFNT | Positive | $1.54 \times 10^8$ |
| Lactamide | PfFNT | Positive | $1.63 \times 10^8$ |
| Lactamide | PfFNT | Positive | $1.36 \times 10^8$ |

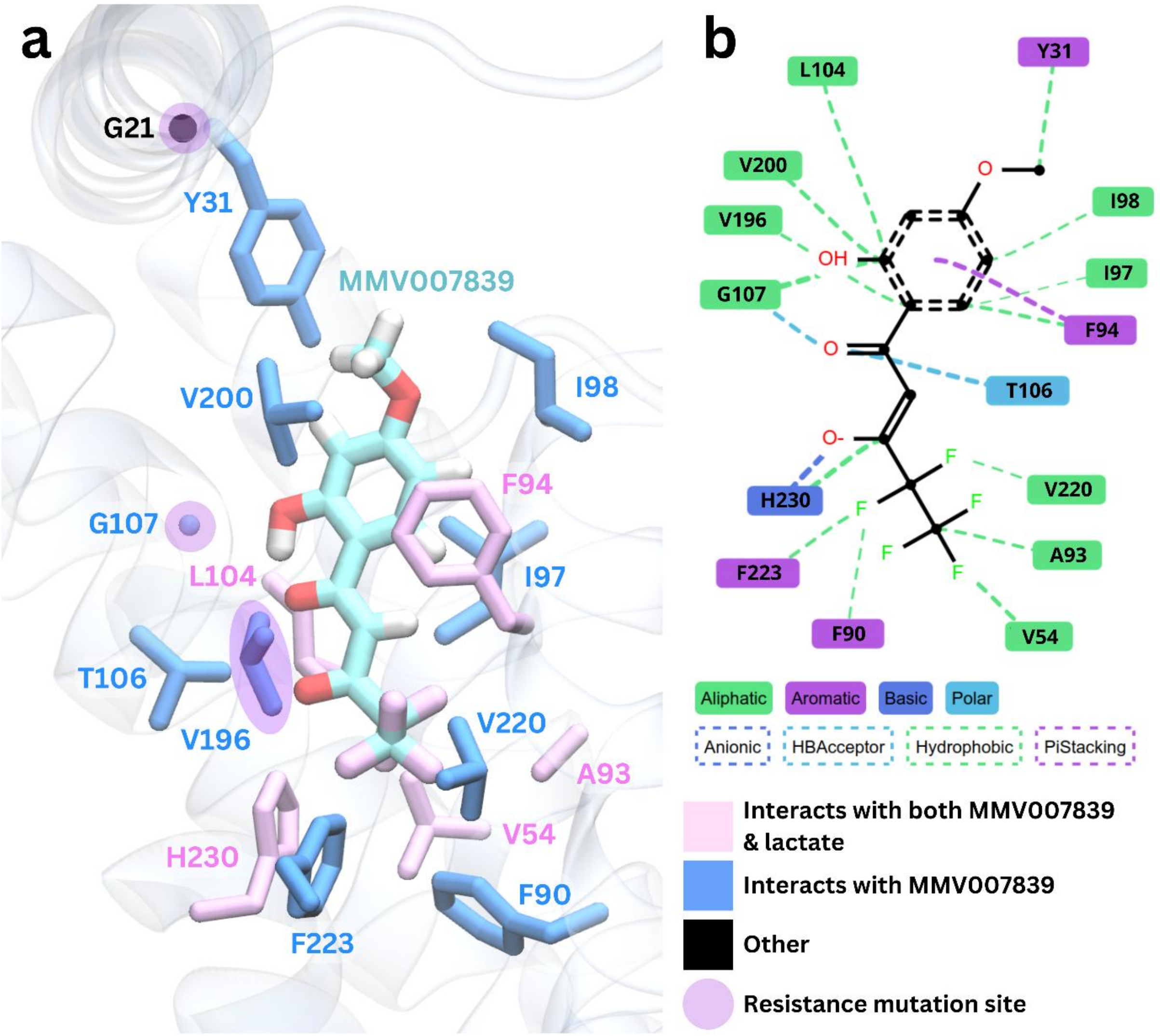


**Figure S16. Overlap between the MMV007839 and lactate binding sites. a)** Snapshot of MMV007839 bound within the transport cavity. Residues identified to contact both MMV007839 and lactate are coloured in pink licorice, residues that contact MMV007839 but do not form strong interactions with lactate are coloured in blue. Residues that fall into neither category are coloured in black. Purple circles are added to signify sites of known resistance mutations. **b)** ProLIF contact map for MMV007839 binding.

### Supplementary References

1. Roberts, E., Eargle, J., Wright, D. & Luthey-Schulten, Z. MultiSeq: Unifying sequence and structure data for evolutionary analysis. *BMC Bioinformatics* **7**, 1–11 (2006).

2. Goujon, M. *et al.* A new bioinformatics analysis tools framework at EMBL–EBI. *Nucleic Acids Res* **38**, W695–W699 (2010).

3. Sievers, F. *et al.* Fast, scalable generation of high-quality protein multiple sequence alignments using Clustal Omega. *Mol Syst Biol* **7**, 539 (2011).

4. Waterhouse, A. M., Procter, J. B., Martin, D. M. A., Clamp, M. & Barton, G. J. Jalview Version 2-A multiple sequence alignment editor and analysis workbench. *Bioinformatics* **25**, 1189–1191 (2009).
